# DANTE and DANTE_LTR: Lineage-centric annotation pipelines for long terminal repeat retrotransposons in plant genomes

**DOI:** 10.1101/2024.04.17.589915

**Authors:** Petr Novák, Nina Hoštáková, Pavel Neumann, Jiří Macas

## Abstract

Long terminal repeat (LTR) retrotransposons constitute a predominant class of repetitive DNA elements in most plant genomes. With the increasing number of sequenced plant genomes, there is an ongoing demand for computational tools facilitating efficient annotation and classification of LTR retrotransposons in plant genome assemblies. Herein, we introduce DANTE, a computational pipeline for Domain-based ANnotation of Transposable Elements, designed for sensitive detection of these elements via their conserved protein domain sequences. The identified protein domains are subsequently inputted into the DANTE_LTR pipeline to annotate complete element sequences by detecting their structural features, such as LTRs, in adjacent genomic regions. Leveraging domain sequences allows for precise classification of elements into phylogenetic lineages, offering a more granular annotation compared with coarser conventional superfamily-based classification methods. The efficiency and accuracy of this approach were evidenced via annotation of LTR retrotransposons in 93 plant genomes. Results were benchmarked against several established pipelines, showing that DANTE_LTR is capable of identifying significantly more intact LTR retrotransposons. DANTE and DANTE_LTR are provided as user-friendly Galaxy tools accessible via a public server (https://repeatexplorer-elixir.cerit-sc.cz), installable on local Galaxy instances from the Galaxy tool shed or executable from the command line.

## Introduction

Long terminal repeat retrotransposons (LTR-RTs) constitute a predominant proportion of repetitive DNA in most higher plant genomes, comprising up to 75% of their nuclear DNA (1). Differential amplification of LTR-RTs is a key driver of genome size evolution (2), contributing to substantial expansion of genomes in species conducive to retrotransposon accumulation (3, 4). Although LTR-RTs have traditionally been considered genomic parasites, they can confer benefits to their hosts by providing regulatory genetic elements (5), driving rapid genomic changes (6), or forming integral parts of specific genomic regions, such as centromeres (7, 8). Investigating these aspects of LTR-RT biology relies on the availability of genome sequencing data across diverse plant taxa as well as computational tools for identifying and accurately annotating LTR-RTs in these data. Although advancements in sequencing and assembly technologies have largely addressed data availability, substantial challenges remain in accurately annotating mobile elements in plant genome assemblies.

LTR-RT annotation entails two primary tasks: identifying element sequences in the assembly and classifying these sequences. Given the extensive element diversity and limited conservation of nucleotide sequences among plant taxa, most current tools employ structure-based methods for LTR-RT identification. Typically, these tools search for pairs of direct repeats representing LTRs and subsequently screen suitable candidates for additional features characteristic of LTR-RTs, such as primer binding sites (PBSs), polypurine tracts, target site duplication (TSD), and protein-coding regions. This methodology is employed by most tools currently used for LTR-RT annotation, including LTR_STRUC (9), LTR_FINDER (10), LTRharvest (11), LTRdigest (12), LtrDetector (13) and MGEScan3 (14). Despite their shared approach, these programs may yield divergent results. For example, some programs excel in identifying true LTR-RTs but also generate a substantial number of false positives, with their performance often depending on the selected parameters and studied organism (15). Consequently, postprocessing tools, such as LTR_retriever (16) and LTRpred (17), have been developed to combine results from multiple tools, enhancing the sensitivity and accuracy of element annotations. Nevertheless, structure-based approaches are computationally intensive and generally less efficient in genomes rich in tandem repeats or nested LTR-RT insertions (16).

Plant LTR-RTs are divided into two superfamilies, Ty1/copia and Ty3/gypsy, distinguished structurally by the arrangement of their protein-coding domains. These superfamilies are further subdivided into families, representing groups of elements with substantial nucleotide sequence similarities [i.e., the 80-80-80 rule, (18)]. However, this system is limited by the broad classification into superfamilies and the highly similar elements within families, typically shared only among closely related species. To address these limitations, we proposed a classification system based on the phylogenetic analysis of conserved domains extracted from LTR-RT polyprotein sequences (19).

Autonomous LTR-RTs encode a polyprotein with at least five protein domains: GAG, protease (PROT), reverse transcriptase (RT), ribonuclease H (RH), and integrase (INT). Cleavage of the polyprotein is due to PROT domain activity, releasing mature proteins essential for the replication and integration of new element copies into the genome. These protein sequences are sufficiently conserved to be efficiently detected in LTR-RTs across phylogenetically distant plant species based on similarities while exhibiting sufficient variation to distinguish specific lineages within the Ty1/copia and Ty3/gypsy superfamilies. Lineages, primarily established through phylogenetic analysis of corresponding protein domains, are further supported by lineage-specific sequence and structural features of elements. This classification system has been implemented in REXdb, a database containing over 75,000 protein sequences from representative green plant genomes. REXdb serves as a reference for consistent, comprehensive LTR-RT annotation across various plant genomes (19). It was originally developed for use in the RepeatExplorer pipeline (20) but later adapted for LTR-RT classification using other tools, such as REPET (21), TEsorter (22) and InpactorDB (23). However, none of these tools primarily employ conserved protein domains for LTR-RT identification in genome assemblies.

Herein, we introduce a novel, protein domain–centric approach for LTR-RT identification and annotation. Instead of primarily searching for LTR-RTs’ structural features, we initially detect and annotate their conserved protein domains. To this end, we perform similarity searches against REXdb and evaluate the resulting hits using the Domain-based ANnotation of Transposable Elements (DANTE) pipeline. Subsequently, we identify clusters of domains from the same lineage and detect LTR-RTs’ structural features in their vicinity using the DANTE_LTR tool. This approach efficiently identifies and annotates intact elements, minimizing false detections and enabling lineage-level classification of identified elements.

## Materials and Methods

### Principles of DANTE and DANTE_LTR

The principles underlying detection and classification of protein domains, as implemented in DANTE, are illustrated schematically in Figure 1A–C. Initially, the program lastal (24) is used to perform a sensitive and frameshift-tolerant DNA–protein similarity search of the query sequence in REXdb. Regions of the query sequence covered by similarity hits exceeding an initial threshold are recorded. Given the database’s redundancy and high search sensitivity, most retrotransposon coding regions in the query sequence are covered by multiple hits. For each region, the hit with the highest similarity score is identified and used to determine the classification score threshold at 80% of the best hit’s similarity score (Figure 1A). Hits surpassing this threshold (“top hits”) are used in DANTE’s classification step. Using multiple similarity hits in this step improves classification robustness by minimizing classification errors due to false similarity hits.

**Figure 1.**
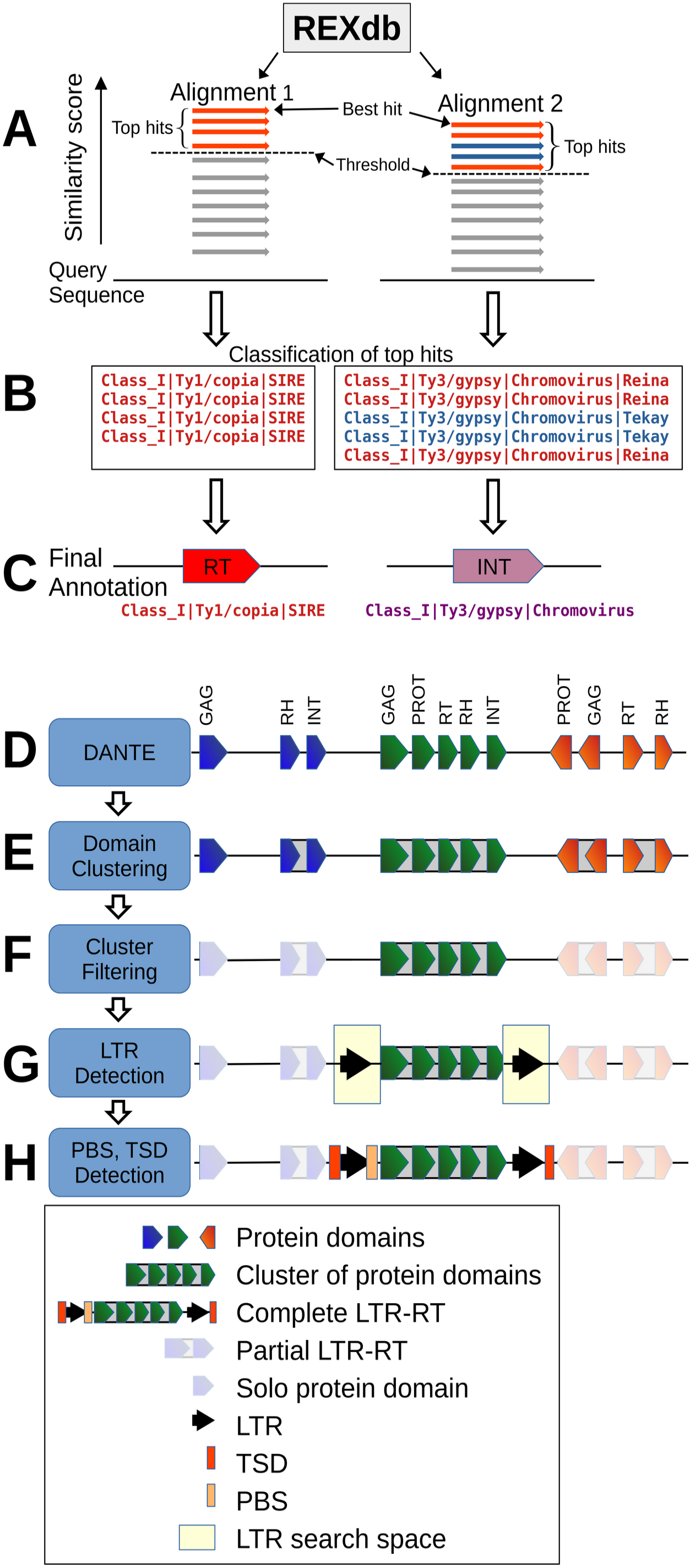
Principles of LTR-retrotransposon annotation achieved via DANTE and DANTE_LTR. Refer to the main text for further details. *Abbreviations*: INT, integrase; LTR, long terminal repeat; PBS, primer binding site; PROT, protease; RH, ribonuclease H; RT, reverse transcriptase; TSD, target site duplication.

In the classification step, DANTE employs a hierarchical classification system for mobile elements implemented in REXdb (Supplementary Figure S1). If all top hits point to a single lineage, this classification is reported as final (illustrated by alignment 1 in Figure 1B and C). If the top hits contain sequences from multiple lineages, the next highest classification level (formally, the lowest common ancestor) including all top hits is reported (alignment 2 in Figure 1B and C). DANTE then outputs domain positions, classifications, and protein sequences as GFF3 and FASTA files.

The algorithm used by DANTE_LTR to identify the nucleotide sequences of complete LTR-RTs is depicted schematically in Figure 1D–H. Initially, DANTE’s output is analyzed to identify clusters of adjacent protein domains likely belonging to the gag-pol coding region of a single element (Figure 1D–F). These clusters must meet specific criteria: (1) all protein domains within a cluster must belong to the same lineage, (2) domains must share the same orientation (strand), (3) domains must be correctly ordered, and (4) distances between neighboring domains must not exceed predefined thresholds. This threshold, along with the typical number and order of domains, was determined for each lineage by analyzing all elements in REXdb. Complete LTR-RTs typically comprise GAG, PROT, RT, RH, and INT domains, with certain lineages encoding additional domains, such as chromodomains (i.e., CHD and CHDCR) from chromoviruses (8) or ancestral RNase H from Tat elements (19). Clusters lacking two or more domains are excluded from further analysis, as they likely represent truncated, rearranged, or nonautonomous elements.

In the next step, DANTE_LTR is used to analyze nucleotide sequences from genome regions upstream and downstream of the domain clusters, aiming to determine the presence of LTRs (Figure 1G). The size of analyzed flanking regions is limited to 2–14 kb based on typical LTR length distributions for different lineages, increased by 25% to accommodate potential size variations. If flanking regions contain additional annotated protein domains indicating another mobile element’s presence, they are truncated to exclude these domains. Nucleotide sequences of the 5′ and 3′ flanking regions are then compared using BLASTN to identify direct repeats. The closest direct repeat to the gag-pol coding region, possessing at least 80% of the shortest LTR length for a given LTR-RT lineage in REXdb, is considered the LTR sequence. If present, TG/CA boundaries typical of LTRs are also reported. Finally, TSD is detected as a short (typically 5 bp) direct repeat surrounding the element sequence, and the element’s PBS is identified based on similarity to a custom plant tRNA sequence database (available as part of the DANTE_LTR GitHub repository).

### Implementation

The DANTE and DANTE_LTR pipelines were developed in Python and R programming languages and are accessible as conda packages. These pipelines operate on the Linux operating system, requiring a minimum hardware configuration of 8 CPUs and 32 GB RAM.

DANTE is designed to analyze FASTA-formatted nucleotide sequences, ranging from a few hundred nucleotides up to whole genome assemblies. The resulting GFF3 file provides the sequence coordinates of identified protein domains, alongside each domain’s type and classification, the name of the REXdb entry providing the best similarity hit, and the translated protein sequence of the annotated region. Notably, DANTE reports all protein domain types present in REXdb, including not only LTR-RTs but also domains encoded by LINEs, DNA transposons, and helitrons (https://doi.org/10.5281/zenodo.10160280). In addition to the complete output, DANTE generates a filtered GFF3 file containing only domains meeting strict filter criteria (45% similarity over at least 80% of the reference sequence, allowing a maximum of 3 frameshifts or stop codons per 100 amino acids), alongside corresponding protein sequences in FASTA format. The filtered output is intended for specific downstream analyses, such as phylogenetic tree generation for domains, whereas DANTE’s full output is used for LTR-RT annotation in DANTE_LTR.

DANTE_LTR is executed using the same input sequences as DANTE, but it requires an additional input: a GFF3 file containing DANTE’s complete output. It produces a GFF3 output file containing detailed annotations of the identified LTR-RTs and a summary table enumerating the identified elements. Optionally, a graphical summary can be generated to provide a detailed overview of LTR-RT structure and size distributions, including their constituent parts (Supplementary File S1).

### Genome assemblies for testing and benchmarking

DANTE_LTR was tested using the genome assemblies of maize (*Zea mays*) and rice (*Oryza sativa*), which have high-quality repeat annotations serving as reliable references. The reference annotation of the maize assembly (acc. no. GCF_000005005.1) was downloaded from MaizeGDB (https://download.maizegdb.org/B73_RefGen_v3/), whereas that of the rice genome (acc. no. GCF_001433935.1) was generated following recommendations (25) using the standard library v7.0.0 from https://github.com/oushujun/EDTA and RepeatMasker version 4.1.1 (parameters: “-e ncbi -pa 36 -q -no_is -norna -nolow -div 40 -cutoff 225”).

As both reference assemblies’ repeat annotations contained overlapping features, resulting in multiple annotations being assigned to portions of genomic repeat regions, these conflicting annotations were resolved as follows. Overlaps between “Ty1/copia” and “Ty3/gypsy” were converted to “LTR-RT unspecified,” whereas other conflicting annotations (e.g., DNA transposons versus LTR-RTs) were converted to “Unknown.” If annotations of overlapping features were identical, they were merged into a single feature to avoid redundancy. These corrections were made using a custom script ensuring that each base in the reference assembly was annotated uniquely or left unannotated. For comparison with annotations generated via tested methods, each base in the reference genome was assigned to one of the following exclusive categories: “Ty1/copia,” “Ty3/gypsy,” “LTR-RT unspecified,” “Other” (including all repeat types other than LTR-RTs), and “Not Annotated.”

To evaluate DANTE_LTR’s performance across diverse plant taxa, additional tests were performed on a set of 91 plant genome assemblies with available LTR-RT annotations selected from a dataset compiled by Zhou et al. (26). Only GenBank accessions that unambiguously matched those provided in (26) were used for testing. The species analyzed and links to their assembly and annotation data are detailed in Supplementary Table S1.

### Performance comparison: DANTE_LTR versus Inpactor2 and EDTA

For comparison with other LTR-RT annotation tools, DANTE (ver. 0.1.9) was executed with its default settings. DANTE_LTR (ver. 0.3.5) was run with the option “-M 1” to tolerate one missing protein domain when identifying intact LTR-RT elements. Although DANTE_LTR identifies both intact and partial LTR-RTs elements, partial elements missing LTR sequences were excluded from the analysis when comparing DANTE_LTR with other tools. Inpactor2 (commit version 3bd8954) (27) was executed with the options “-t 5 -m 35000 -C 3.” The option “-m” set the maximum length of LTR-RTs to match the value used in DANTE_LTR, and the “-C 3” option was employed to perform the detection process in three cycles, with this multicycle analysis aimed at mitigating potential element splitting issues when analyzed sequences are partitioned (27). LTR_retriever was used as implemented in the Extensive *de-novo* TE Annotator (EDTA) pipeline (ver. 2.0.1) (25) with the parameters “--species others --overwrite 0 --step all --sensitive 1.” Annotations of complete elements generated by DANTE_LTR, Inpactor2, and EDTA were preprocessed to remove overlapping features within each output file and converted into a unified classification containing only superfamily-level categories: “Ty1/copia,” “Ty3/gypsy,” and “LTR-RT unspecified.” This step ensured annotation comparability, as EDTA lacks classification of identified elements at the lineage level, unlike DANTE_LTR and Inpactor2. The modified annotations were then analyzed to determine the total number of elements identified by each tool and the proportion of elements identified by multiple tools simultaneously (i.e., annotations overlapping >90% of their length).

To compare annotations generated by each tool with reference repeat annotations from the rice and maize genomes, each base in the respective genome was assigned to one of four categories based on the match between the tested and reference annotation: true positive, true negative, false positive, and false negative. The number of bases assigned to each category was used to calculate the benchmarking metrics of sensitivity, specificity, accuracy, false discovery rate (FDR), precision, and F1 score (Supplementary Figure S2) using custom R scripts.

For similarity-based annotation of LTR-RTs, reference databases containing nucleotide sequences of elements identified by individual tools were compiled and used for a RepeatMasker search with the parameters “-e ncbi -pa 36 -q -no_is -norna -nolow -div 40 - cutoff 225.” Resulting annotations were processed and evaluated as described earlier.

Benchmarking of analysis execution times was conducted on a computer with Intel(R) Xeon(R) CPU E5-2630 v4 @ 2.20GHz, 32 CPU, and 64 GB RAM running Ubuntu 16.4 Linux. In addition to executing the programs detailed earlier, Inpactor2 was run with the “-C 1” option to use settings identical to those previously described (27). Benchmarking was performed with the whole genome assemblies of *O. sativa* and *Z. mays* as well as the centromere 6 assembly of *Pisum sativum* [GCA_947076115.1 (28)].

## Results

We assessed the performance of DANTE and DANTE_LTR using assemblies from 93 flowering plant species (Magnoliopsida). In the initial testing phase, we analyzed rice and maize genome assemblies to evaluate the sensitivity and accuracy of LTR-RT identification. These assemblies were chosen owing to their detailed and high-quality annotations of all repeat types, offering suitable references for benchmarking. In the subsequent testing phase, we examined DANTE_LTR’s capacity to identify LTR-RTs across 91 species, varying in genome assembly size (42–19,000 Mb), representing diverse taxonomic groups of flowering plants (Supplementary Table S1). These species’ assemblies were previously analyzed for the presence of LTR-RTs by Zhou et al. (26) using a combination of LTRharvest and LTRdigest. Additionally, we compared DANTE_LTR’s performance with those of two established pipelines run with the same set of assemblies: Inpactor2 (27), employing a machine learning approach, and EDTA, using a combination of structure-based methods for LTR-RT annotation (25).

### Evaluation of DANTE_LTR using reference annotations from rice and maize genomes

Analysis of the rice and maize genome assemblies using DANTE_LTR resulted in the identification and annotation of 2771 and 48,276 complete elements, respectively. Elements were considered complete if they contained both LTRs and an internal protein-coding region with conserved protein domains recognized by DANTE in the correct order and orientation, given tolerance for one missing domain. Primer binding sites and target site duplications were also detected and annotated. Elements were classified into the phylogenetic lineages of LTR-RTs defined in REXdb (19), revealing varying frequencies of elements from individual lineages in the analyzed genomes (Table 1 and Supplementary Table S2).

**Table 1.**
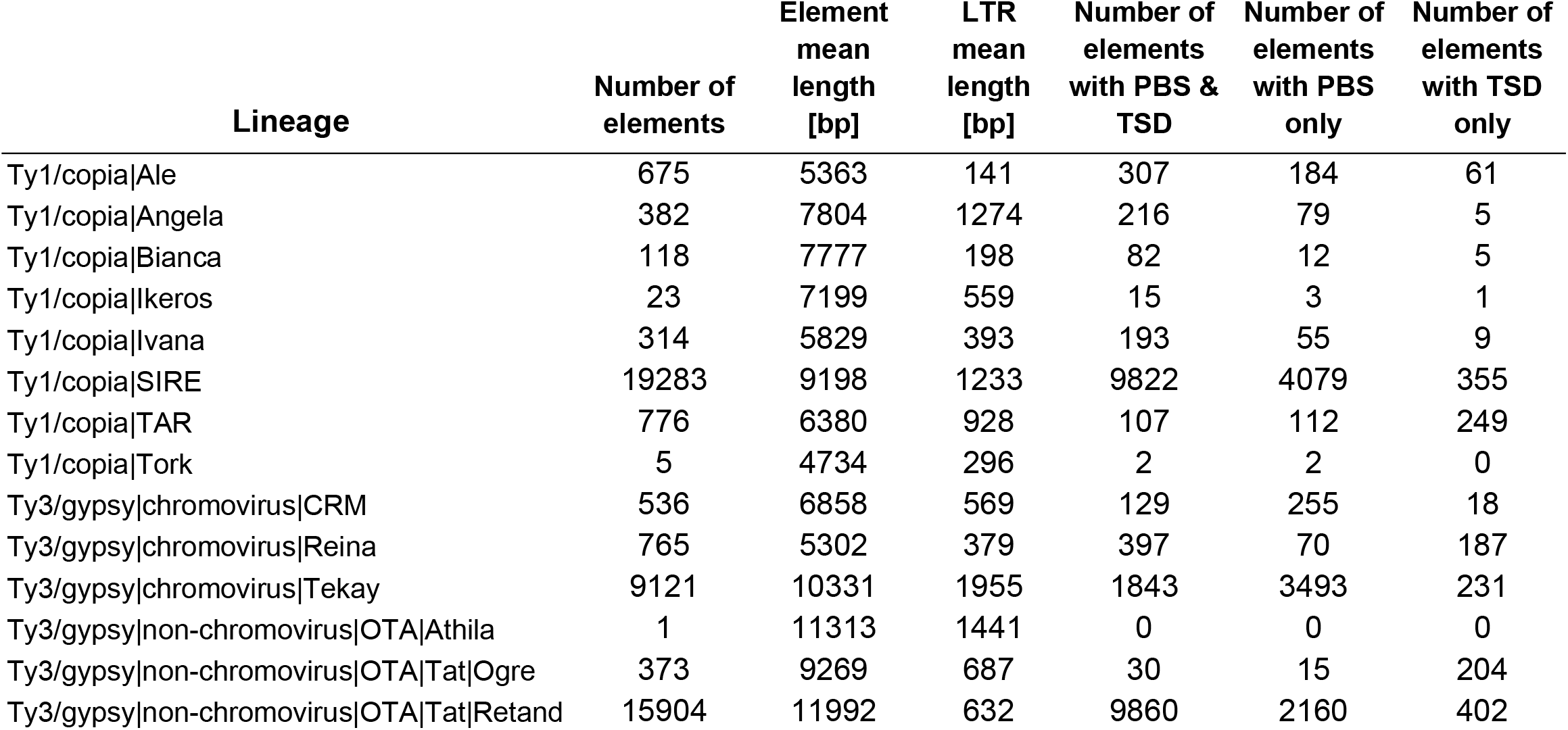
Number of elements identified in the *Z. mays* genome using DANTE_LTR.

DANTE_LTR not only provides LTR-RT annotations in GFF3 files but also generates a summary report and a graphical representation of the detected elements’ various features. In addition to the number of identified elements (Table 1 and Supplementary Table S2), it provides an overview of their consensus structures (Figure 2A) as well as the size and structural variations of elements within individual lineages (Figure 2B–D). Each lineage is further analyzed in terms of sequence divergence of LTRs in individual elements and primer binding site types (Figure 2E and F). Complete summary reports for rice and maize, including data for all identified LTR-RT lineages, are provided in Supplementary File S1.

**Figure 2.**
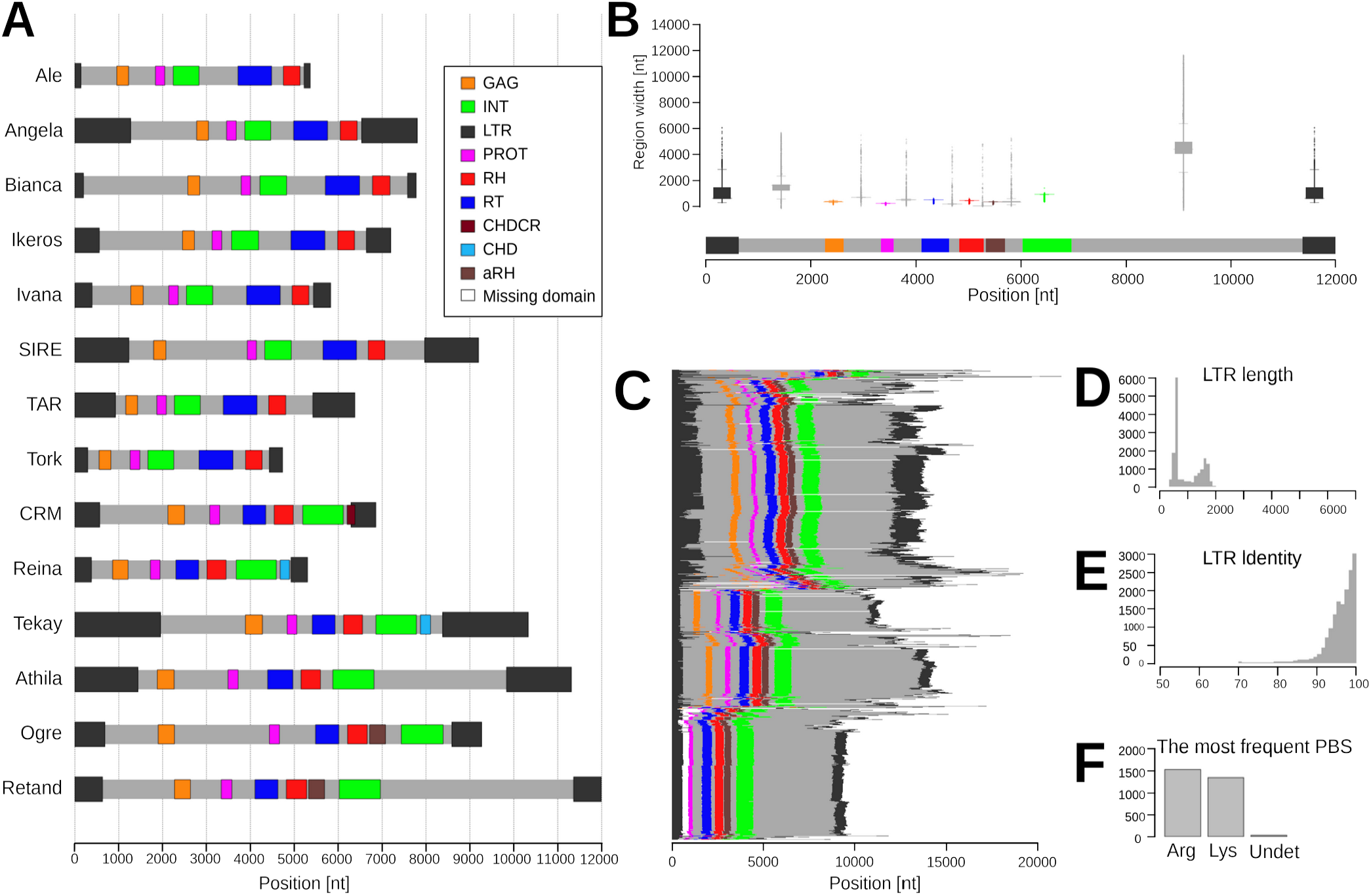
Examples of information provided in DANTE_LTR summary reports. (**A**) Consensus structure of all LTR-RT lineages identified in the analyzed genome (*Zea mays*). (**B–F**) Detailed information on individual lineages, using the example of Ty3/gypsy Retand from the *Z. mays* genome. The report includes the consensus structure and a boxplot depicting length variations of element features derived from all identified elements (**B**), the structures of which are shown in (**C**) as horizontal lines, with colors indicating sequence and structural feature positions. Histograms illustrate the LTR size distribution (**D**) and sequence similarities within each element (**E**) as well as the occurrences of the most common PBS types detected in each lineage (**F**). Complete summary reports are provided in Supplementary File S1.

Comparison of LTR-RT annotations generated by DANTE_LTR with reference annotations revealed high specificity and a low FDR (Figure 3A). DANTE_LTR outperformed both Inpactor2 and EDTA in terms of the number of complete elements identified (Figure 3B). Furthermore, DANTE_LTR exclusively identified 31.8% and 39.7% more elements in the rice and maize assemblies, respectively, compared with the other tools (Figure 3C). This higher element count was attributed to DANTE_LTR’s superior sensitivity, coupled with consistently low FDRs (1.4% and 6.2% for rice and maize, respectively). Additionally, DANTE_LTR exhibited a superior F1 score, indicating a balanced trade-off between precision and sensitivity, while achieving higher overall annotation accuracy (Figure 3A; Supplementary Table S3).

**Figure 3.**
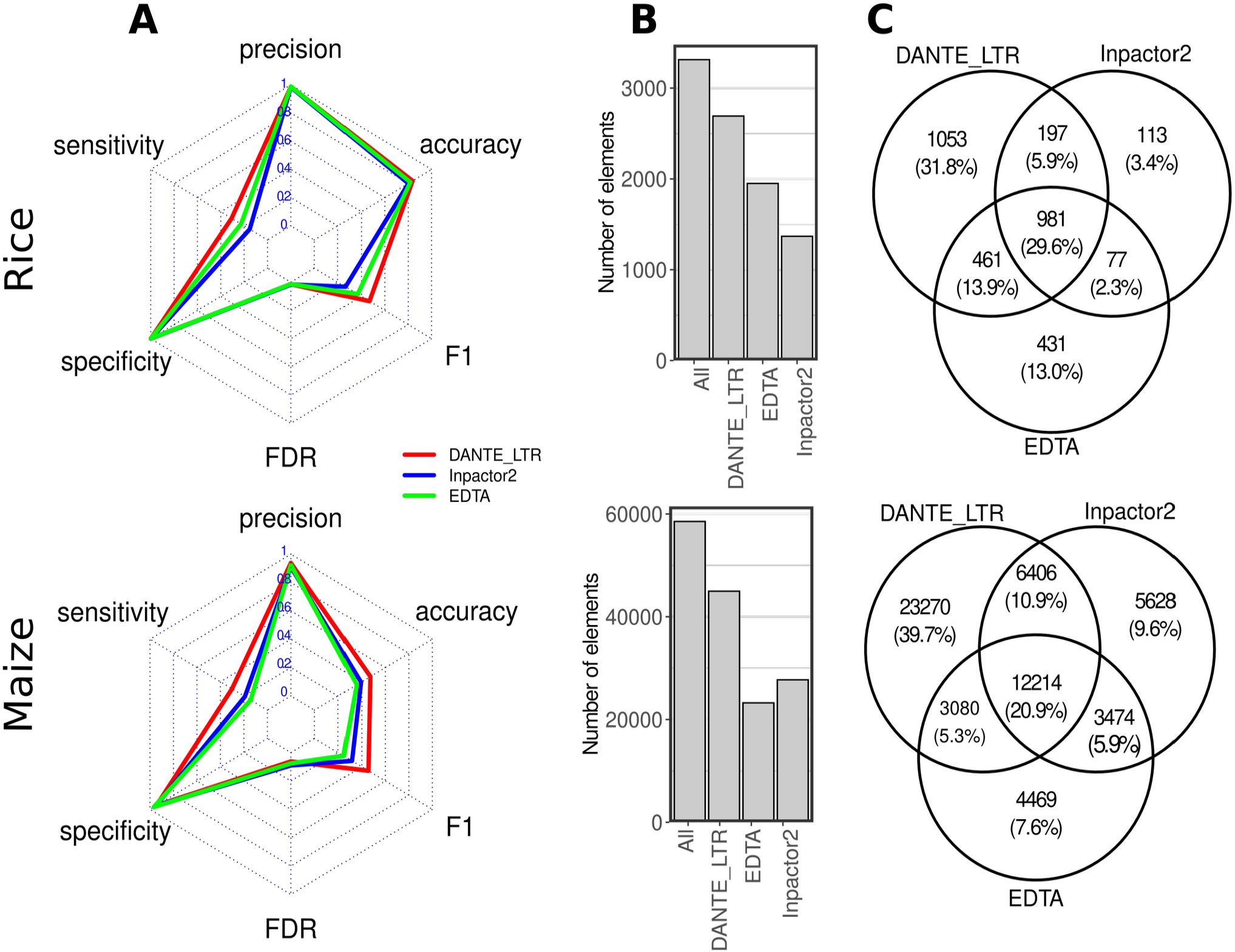
Comparison of LTR-RT annotations generated using DANTE_LTR, Inpactor2, and EDTA for the rice and maize genomes. (**A**) Annotation performance metrics of the three pipelines, calculated by comparing their results with reference annotations. (**B**) Number of identified elements. “All” bar corresponds to elements identified by at least one method. **(C)** Quantities (and percentages) of unique elements and those identified by two or all pipelines simultaneously. Refer to Supplementary Figure S2 for a detailed metrics explanation.

As the tools developed for identifying complete elements are inherently unable to annotate nonautonomous elements and fragmented LTR-RT sequences, they often include an additional annotation step based on sequence similarity. This approach was tested using RepeatMasker similarity search with reference libraries containing nucleotide sequences from the complete elements identified in previous analysis (Figure 4A and B). The sensitivity of LTR-RT sequence detection was slightly lower for the reference libraries generated from DANTE_LTR output compared with those generated using EDTA and Inpactor2: 75%, 96%, and 91% for the rice genome and 84%, 85%, and 94% for the maize genome, respectively. However, the higher sensitivities of EDTA and Inpactor2 were associated with 2–4-fold higher FDRs (Figure 3C). The specificity of similarity-based annotation was influenced by the proportion of misclassified sequences in the reference libraries. Although such contamination was minor, it led to a considerable increase in false positives in subsequent annotations. This effect was observed across all three pipelines, but DANTE_LTR maintained the lowest FDR (Supplementary Figure S3).

**Figure 4.**
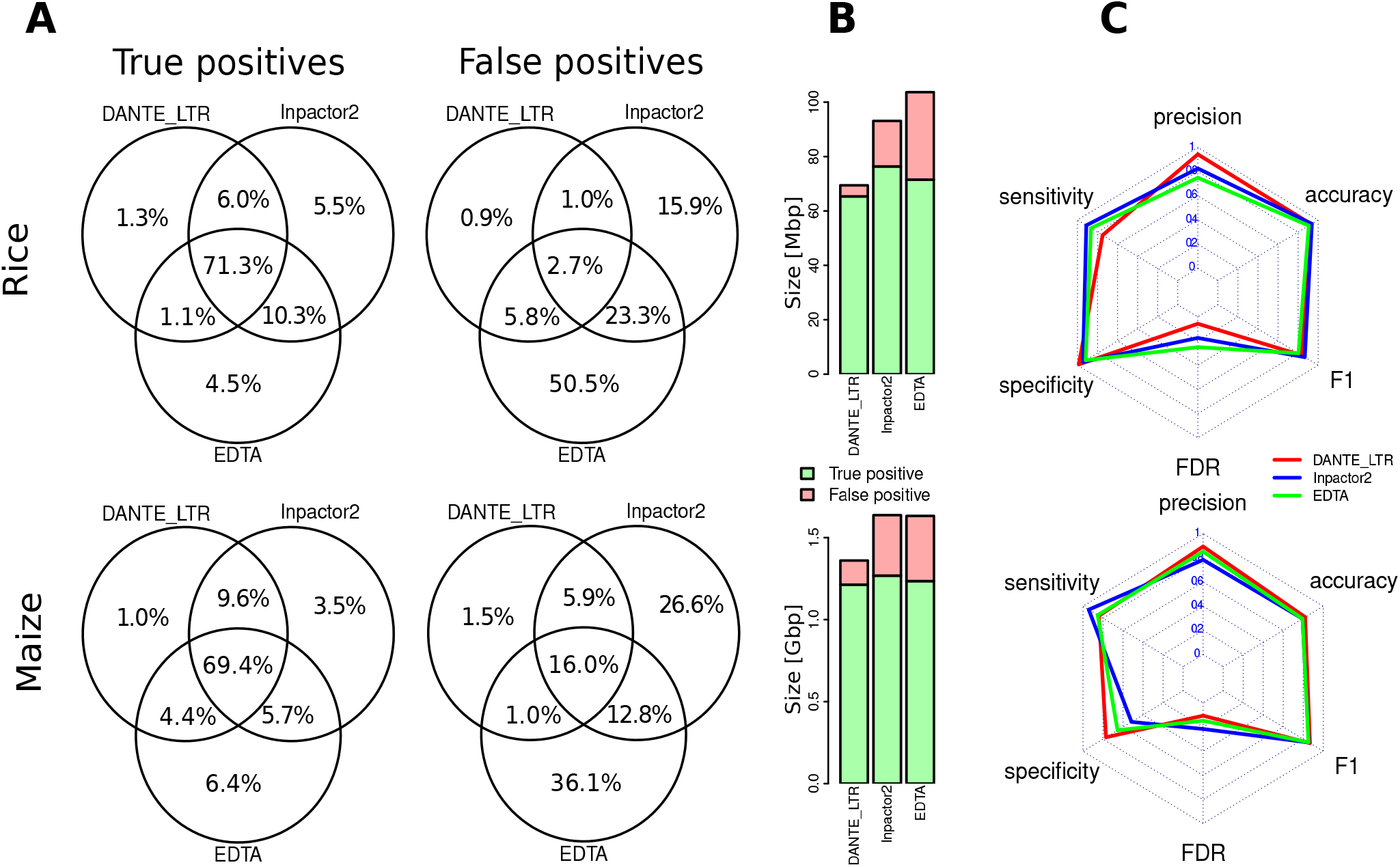
Similarity-based LTR-RT annotations using libraries of elements identified in the rice and maize genomes using DANTE_LTR, Inpactor2, and EDTA, evaluated against reference annotations of the rice and maize genomes. (**A**) Overlaps between true- and false-positive annotations generated by the compared pipelines. (**B**) Total length of annotated regions. (**C**) Annotation performance metrics of the three pipelines, calculated by comparing their results with reference annotations. Refer to Supplementary Figure S2 for a detailed metrics explanation.

### LTR-RT detection in a set of 91 flowering plant species

Given that DANTE_LTR’s performance relies on recognizing and accurately classifying the retrotransposon protein domains, we initially verified that the domains reported by DANTE were unambiguously assigned to specific LTR-RT lineages defined in REXdb (Supplementary Figure S1). An opposing scenario, where a large proportion of domains are only assigned to a general category as a superfamily, would suggest that corresponding elements from phylogenetically diverse plant species are not adequately represented in REXdb, hindering efficient identification. However, this scenario was ruled out, as most of the 21,054,179 identified domains were unambiguously classified for both the Ty1/copia (97.7%) and Ty3/gypsy (97.0%) lineages (Supplementary Figure S4A). The number of identified domains generally increased with assembly size, reflecting a higher proportion of LTR-RTs in species with larger genomes (Supplementary Figure S5).

Subsequent analysis of identified domains and their adjacent genomic regions by DANTE_LTR resulted in the annotation of 819,175 complete LTR-RT sequences (67– 166,487 per species). Complete element sequences were defined by the presence of LTRs and the gag-pol region. Of the identified protein domains, 38.7% and 34.8% were assigned to complete elements of the Ty1/copia and Ty3/gypsy lineages, respectively, with the remaining domains being either solo domains or part of partial elements (Supplementary Figure S4B). A strong positive correlation was observed between the number of protein domains and the number of complete elements identified in individual species, with relatively consistent R^2^ values ranging from 0.8 (Bianca lineage) to 0.97 (Retand lineage) observed across lineages (Supplementary Figure S6).

Compared with the other tools, DANTE_LTR identified significantly more complete elements for both Ty1/copia and Ty3/gyspy (Figure 5A and B). The number of identified elements was comparable to that reported by Zhou et al. (26); however, only 43%–44% of elements were identified in both analyses, with 17.3%–17.7% of elements identified only by DANTE_LTR and 11.2%–15.8% being specific to the Zhou et al. annotation (Figure 5C and D). Similar partial overlap was observed in the other two tools’ annotations, with only 5.1%–9.1% of elements common to all four annotations (Figure 5C and D).

**Figure 5.**
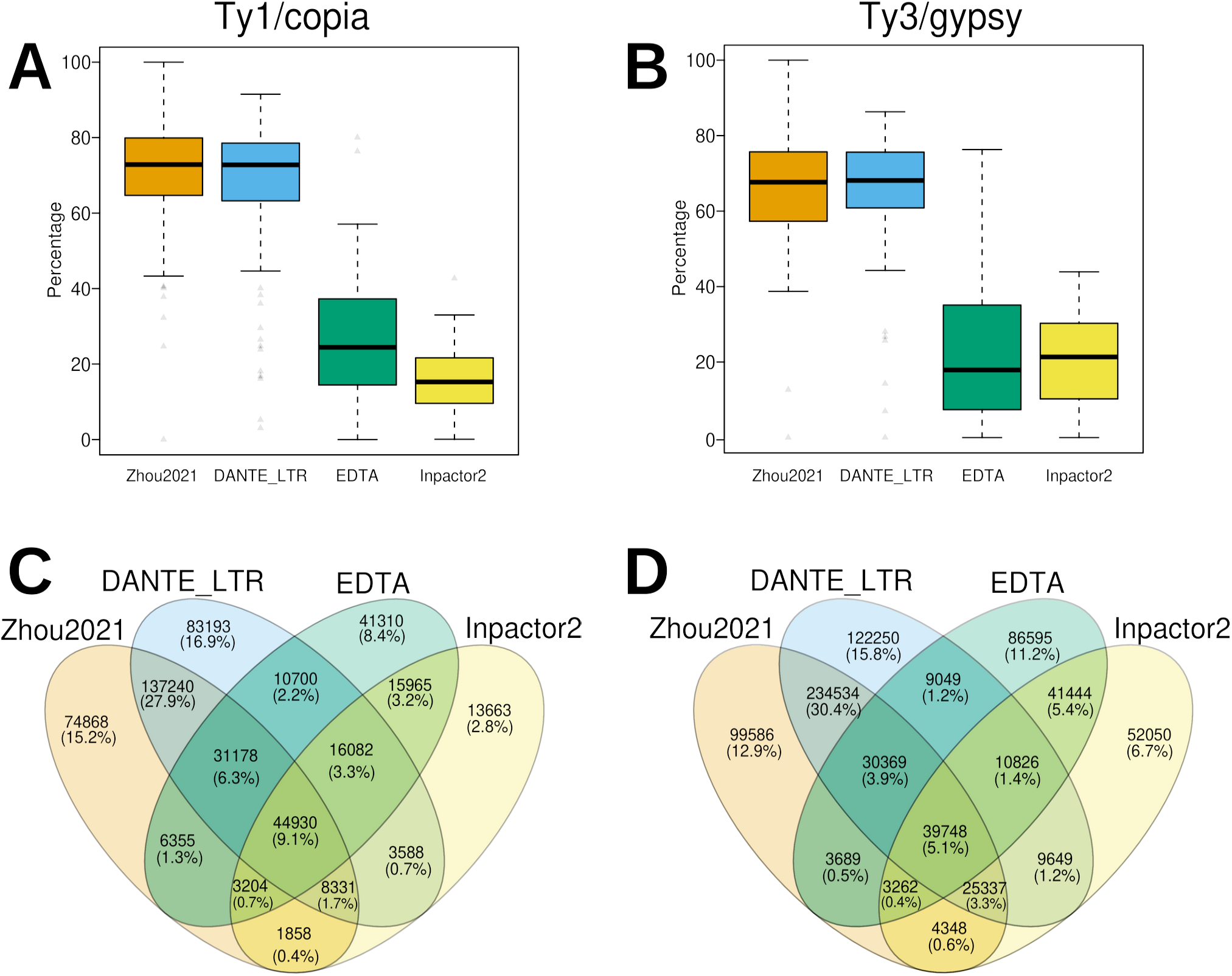
LTR-RT annotation across 91 plant species. Proportions of Ty1/copia (**A**) and Ty3/gypsy (**B**) elements annotated in the analyzed assemblies via the tested pipelines and by Zhou et al. (26) (referred to as “Zhou2021”). Proportions calculated as percentages of the combined number of elements identified using all methods. (**C, D**) Quantities (and percentages) of unique elements and those identified simultaneously by two or more pipelines, separately calculated for Ty1/copia (**C**) and Ty3/gypsy (**D**) elements.

### Execution time

Benchmarking of execution time for DANTE_LTR, EDTA, and Inpactor2 was performed using the rice and maize genome assemblies as well as the 178 Mb assembly of the *P. sativum* chromosome 6 centromeric region (28). This region is rich in tandemly organized satellite DNA, posing a major challenge for structure-based tools regarding LTR-RT annotation. DANTE_LTR demonstrated faster performance compared with EDTA and Inpactor2 across all three assemblies, except for one instance where Inpactor2 was run without iterations (option “-C 1”), resulting in the quickest analysis of the maize assembly (Table 2). However, when Inpactor2 was executed with three iterations, as recommended (option “-C 3”), its execution time increased markedly, particularly in maize genome analysis (a 36-fold increase), likely due to the heightened complexity of output processing. Nevertheless, excluding this exceptional case, the three tools’ execution times remained relatively comparable, fluctuating within a 1.3–3.7-fold range. Measured in megabases of analyzed assemblies per minute (Mbp/min), analysis of the rice genome was markedly faster for all tools compared with the LTR-RT–rich genomes of maize and *P. sativum*. Notably, DANTE_LTR exhibited the highest efficiency when analyzing the satellite DNA–rich centromere of *P. sativum* (Table 2).

**Table 2.**
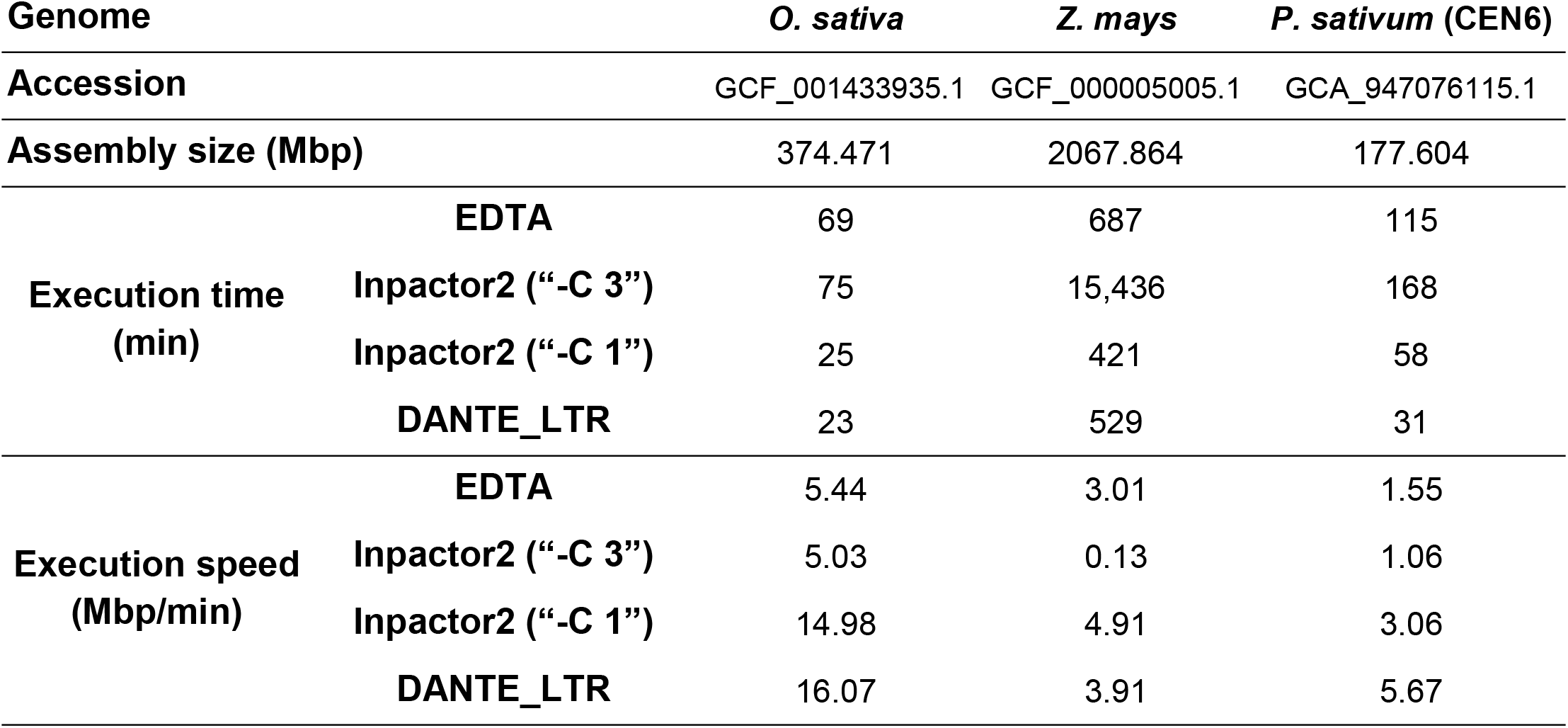
Execution time (min) benchmarks for DANTE_LTR, Inpactor2, and EDTA.

| Genome |  | *O. sativa* | *Z. mays* | *P. sativum* (CEN6) |
| --- | --- | --- | --- | --- |
| Accession |  | GCF_001433935.1 | GCF_000005005.1 | GCA_947076115.1 |
| Assembly size (Mbp) |  | 374.471 | 2067.864 | 177.604 |
| Execution time (min) | EDTA | 69 | 687 | 115 |
|  | Inpactor2 (“-C 3”) | 75 | 15,436 | 168 |
|  | Inpactor2 (“-C 1”) | 25 | 421 | 58 |
|  | DANTE_LTR | 23 | 529 | 31 |
| Execution speed (Mbp/min) | EDTA | 5.44 | 3.01 | 1.55 |
|  | Inpactor2 (“-C 3”) | 5.03 | 0.13 | 1.06 |
|  | Inpactor2 (“-C 1”) | 14.98 | 4.91 | 3.06 |
|  | DANTE_LTR | 16.07 | 3.91 | 5.67 |

## Discussion

In this study, we introduced and validated a new method for annotating LTR-RTs, focusing on their highly conserved protein-coding regions. Besides enhancing element detection sensitivity, conserved protein domain sequences enable precise classification into phylogenetic lineages, surpassing the level of detail in conventional superfamily classification (Ty1/copia and Ty3/gypsy), with general applicability across plant taxa (19). Moreover, during individual element annotation, an abundance of data are collected, including element structure and LTR, PBS, and TSD region properties, with this information being summarized for each LTR-RT lineage. These data not only provide an overview of element population variability in studied species but also facilitate various downstream studies, such as dating element insertions based on LTR sequence divergence. These features are unique to DANTE_LTR compared with similar tools, which typically offer only genome annotation files.

Two potential limitations of the DANTE_LTR approach to identifying LTR-RTs must be acknowledged. First, elements with protein sequences divergent from those in REXdb may not be identified. However, owing to the relatively high conservation of LTR-RT protein domains, efficient recognition in elements from related plant taxa is feasible with a limited number of representative reference sequences. Although the current REXdb version (ver. 3.0) fully covers flowering plants (angiosperms), other green plant groups are under-represented, potentially limiting DANTE_LTR’s sensitivity; therefore, database updates are required in the future. Second, DANTE_LTR cannot identify nonautonomous elements lacking protein-coding domains. However, this issue can be mitigated by performing a similarity search in the second step of the annotation procedure, using an identified element library from the first step as a reference. In this case, only element lineages lacking a complete element in the analyzed genome would be overlooked.

Compared with other tools designed for LTR-RT annotation in plant genomes, DANTE_LTR demonstrated higher sensitivity, identifying significantly more elements compared with EDTA and Inpactor2. Additionally, DANTE_LTR exhibited greater specificity given its evaluation of multiple patterns, including protein domain type and arrangement, LTRs, TSDs, PBSs, and TG/CA boundaries. Notably, limited overlap was observed in LTR-RT annotations generated through the three pipelines as well as those generated by Zhou at al. (26), with most elements identified by only one or a few tools, with those identified by all tools simultaneously being rare. This may be attributed to high sequence and structural variation in elements across plant genomes, alongside varying efficiencies among approaches for identifying specific element variants. This result emphasizes the importance of using multiple tools with different recognition principles, rather than relying on a single tool, to achieve LTR-RT annotations in plant assemblies.

Importantly, benchmarking genome annotation tools, including those for repeat annotation, inherently suffers from the lack of a definitive “ground truth.” Reference annotations are typically considered the best approximation to the ground truth and serve as benchmarks for new annotations. However, this approach assumes the completeness and accuracy of reference annotations, which may not always hold true. Thus, benchmarking primarily compares the relative performance of tools against each other and existing reference annotations, offering valuable insights into method strengths and weaknesses but lacking an absolute measure of annotation reliability.

Some limitations of structure-based LTR-RT annotation approaches can be addressed by an additional step using nucleotide sequence similarity detection in a reference library of complete elements (25, 27). This step notably enhanced DANTE_LTR output, markedly increasing the proportion of annotated LTR-RT sequences by detecting nonautonomous or fragmented element sequences. A similar effect was observed in the comparison pipelines, EDTA and Inpactor2 (Figure 4). Therefore, performing this step generally improves analysis results. However, the reference library of complete elements must be error-free, i.e., free from chimeric or incorrectly annotated elements. As some degree of contamination is inevitable, identifying and removing such sequences is crucial to prevent further erroneous annotations (Supplementary Figure S3). Among the compared pipelines, DANTE_LTR was the least affected by this issue; nevertheless, thorough validation and curation of the reference library are recommended. Graphical summary reports for each identified element lineage aid in this process, helping identify and remove elements with atypical structures or unexpectedly long noncoding regions potentially containing unrelated element insertions, such as MITEs or other DNA transposons.

## Data availability

DANTE and DANTE_LTR source code are available under GPLv3 license at https://github.com/kavonrtep/dante and https://github.com/kavonrtep/dante_ltr, respectively. DANTE and DANTE_LTR source code for Galaxy implementation are available at https://toolshed.g2.bx.psu.edu/view/petr-novak/dante and https://toolshed.g2.bx.psu.edu/view/petr-novak/dante_ltr, respectively. Custom scripts for annotation data analysis are available at https://github.com/kavonrtep/dante_ltr_evaluation. Output from DANTE_LTR, Inpactor2, and EDTA annotations of 93 analyzed species is available in Zenodo at https://zenodo.org/doi/10.5281/zenodo.10891048. Both DANTE and DANTE_LTR programs can be used on the public Galaxy server (https://repeatexplorer-elixir.cerit-sc.cz/), which is provided by the ELIXIR-CZ project.

## Author contributions

JM, PNo, and PNe designed the study. PNo and NH programmed the pipelines, PNo analyzed the data, and JM and PNo wrote the manuscript. All authors read and approved the final manuscript.

## Funding

This work was supported by the ELIXIR-CZ Research Infrastructure Project (LM2023055).

## Notes

### Competing Interest Statement

The authors have declared no competing interest.

### Summary of Updates

This version of the manuscript has been revised to change its formatting and improve writing style. Some references have also been corrected. The data presented has not been changed.

