## Supplementary Figures for "DANTE and DANTE_LTR: Lineage-centric annotation pipelines for long terminal repeat retrotransposons in plant genomes"

```
LTR-RT
  ├── Ty1/copia
  │     ├── Ale
  │     ├── Alesia
  │     ├── Angela
  │     ├── Bianca
  │     ├── Bryco
  │     ├── Lyco
  │     ├── Gymco-I
  │     ├── Gymco-II
  │     ├── Gymco-III
  │     ├── Gymco-IV
  │     ├── Ikeros
  │     ├── Ivana
  │     ├── Osser
  │     ├── SIRE
  │     ├── TAR
  │     ├── Tork
  │     └── Ty1-outgroup
  │
  └── Ty3/gypsy
        ├── non-chromovirus
        │     ├── non-chromo-outgroup
        │     ├── Phygy
        │     ├── Selgy
        │     └── OTA
        │          ├── Athila
        │          └── Tat
        │               ├── TatI
        │               ├── TatII
        │               ├── TatIII
        │               ├── Ogre
        │               └── Retand
        │
        └── chromovirus
              ├── Chlamyvir
              ├── Tcn1
              ├── chromo-outgroup
              ├── CRM
              ├── Galadriel
              ├── Tekay
              ├── Reina
              └── chromo-unclass
```

**Supplementary Figure S1**. REXdb LTR-retrotransposon classification system

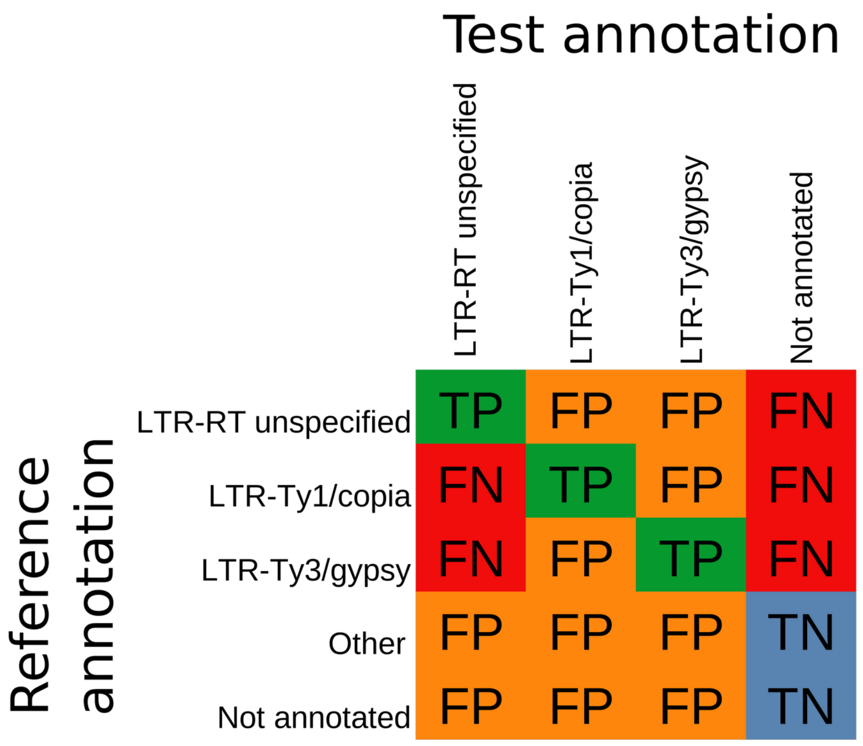


$$Sensitivity = \frac{TP}{TP + FN} \qquad Accuracy = \frac{TP + TN}{TP + TN + FP + FN}$$

$$Specificity = \frac{TN}{TN + FP} \qquad Precision = \frac{TP}{TP + FP}$$

$$F1 = 2 \times \frac{Precision \cdot Sensitivity}{Precision + Sensitivity} \qquad FDR = \frac{FP}{TP + FP}$$

**Supplementary Figure S2**. Assignment of bases in the assembly to true positive (TP), true negative (TN), false positive (FP), and false negative (FN) categories according to annotation in the reference and annotation under the test. Formulas show definitions of sensitivity, specificity, F1, accuracy, precision and false discovery rate (FDR).

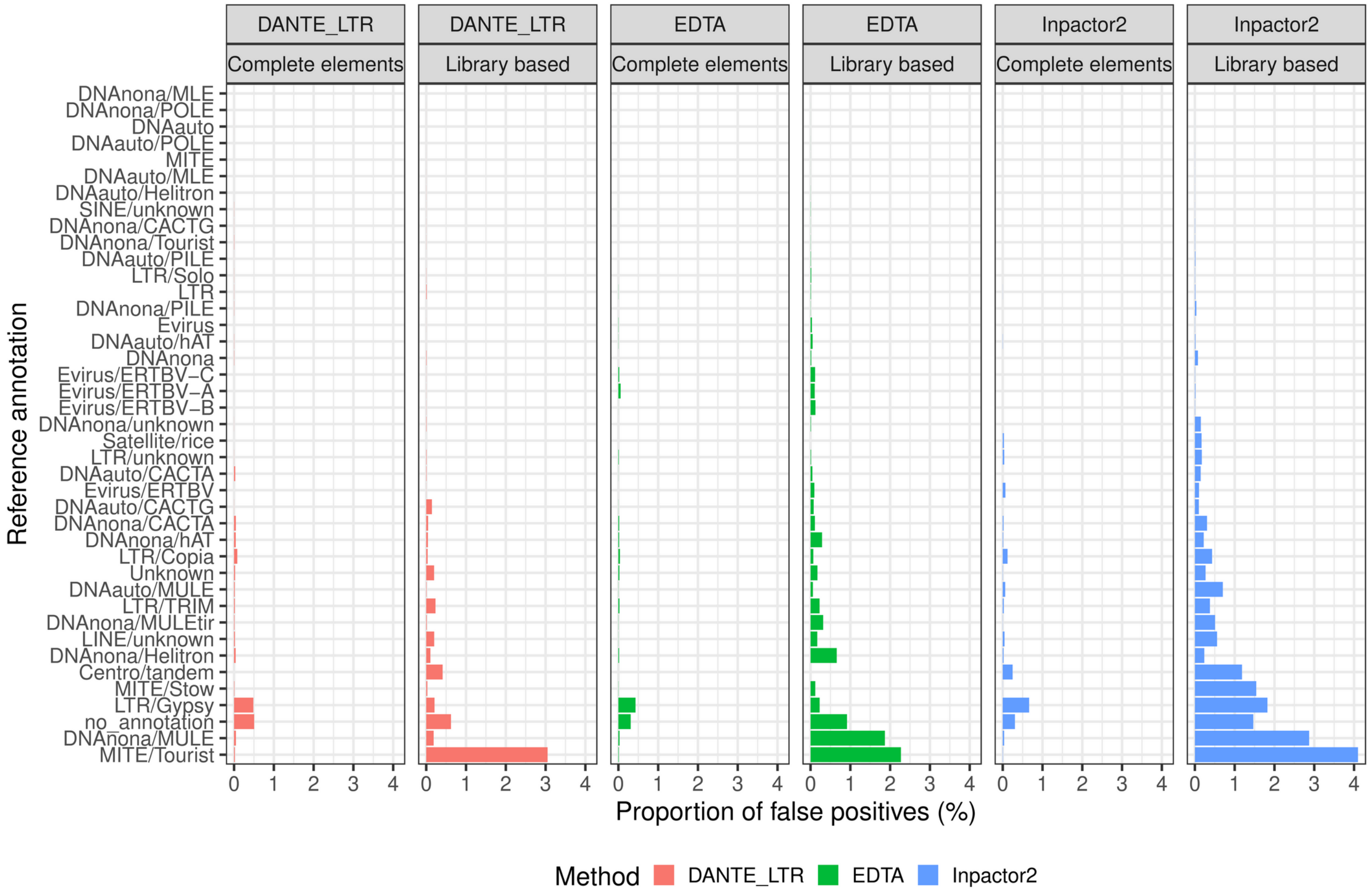


**Supplementary Figure S3**. Comparison of the structure-based annotation of complete elements using DANTE_LTR, EDTA, and Inpactor2 with library-based annotation. The plot shows the classification of false positive regions as per the reference annotation in *O.sativa*. Bar plots illustrate the proportion of these false positive regions relative to the total annotated regions. Each bar represents the percentage of regions incorrectly annotated compared to the reference annotation.

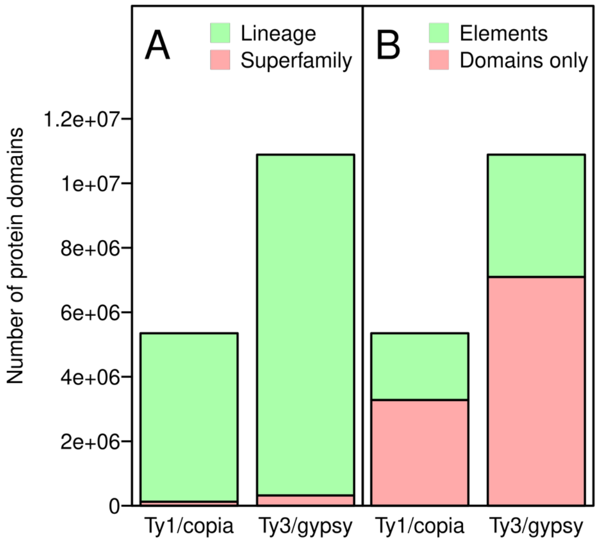


**Supplementary Figure S4**. Annotation of 91 *Magnoliopsida* assemblies by DANTE and DANTE_LTR (**A**) Barplots show the proportions and numbers of proteins domains identified by DANTE which could be assigned to lineage level classification and those which could be classified only to a superfamily level. (**B**) Numbers of protein domains assigned by DANTE_LTR to  elements and number of domains without assignment.

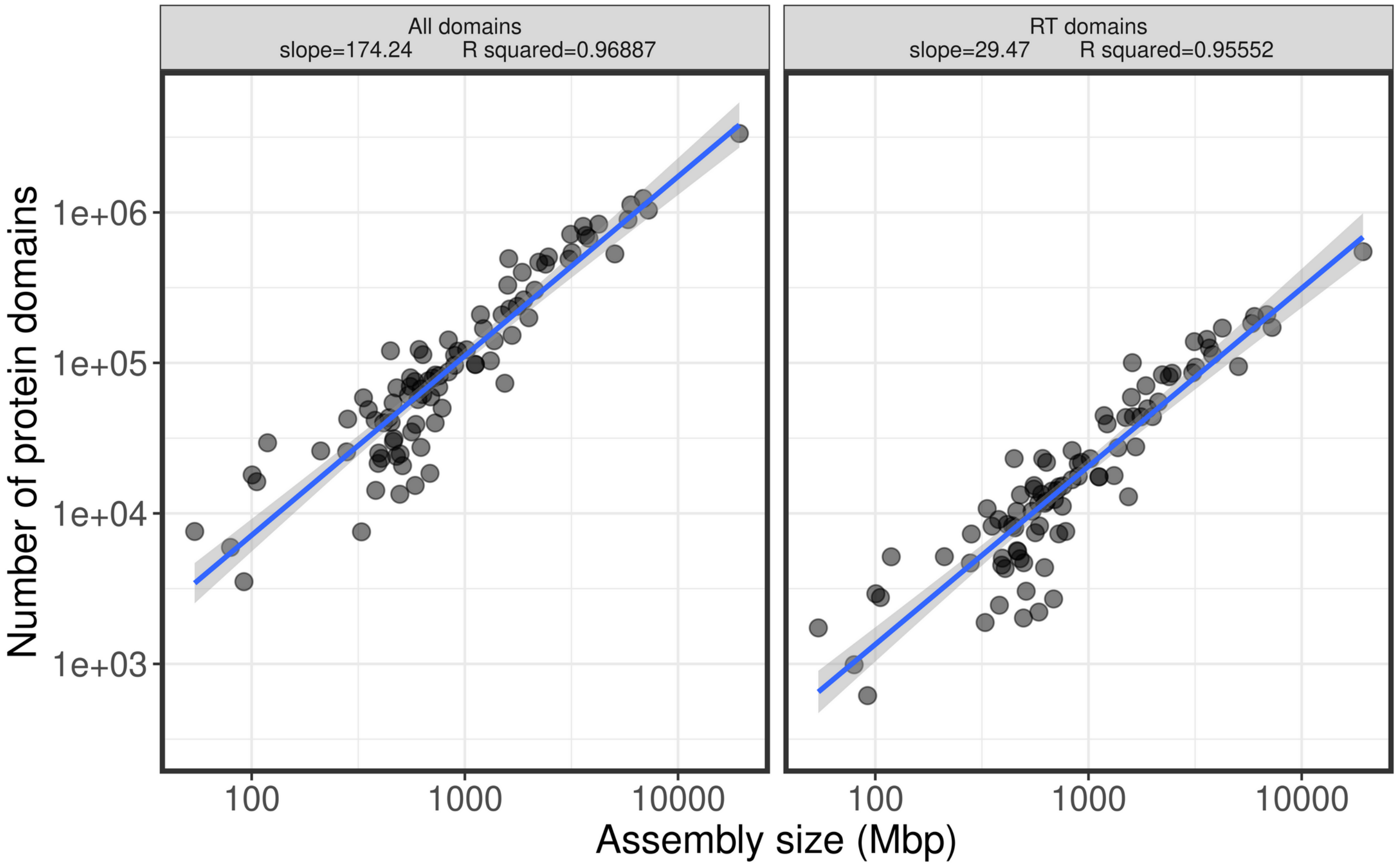


**Supplementary Figure S5.** Number of detected LTR-RT protein domains in analyzed species is proportional to analyzed assembly size.

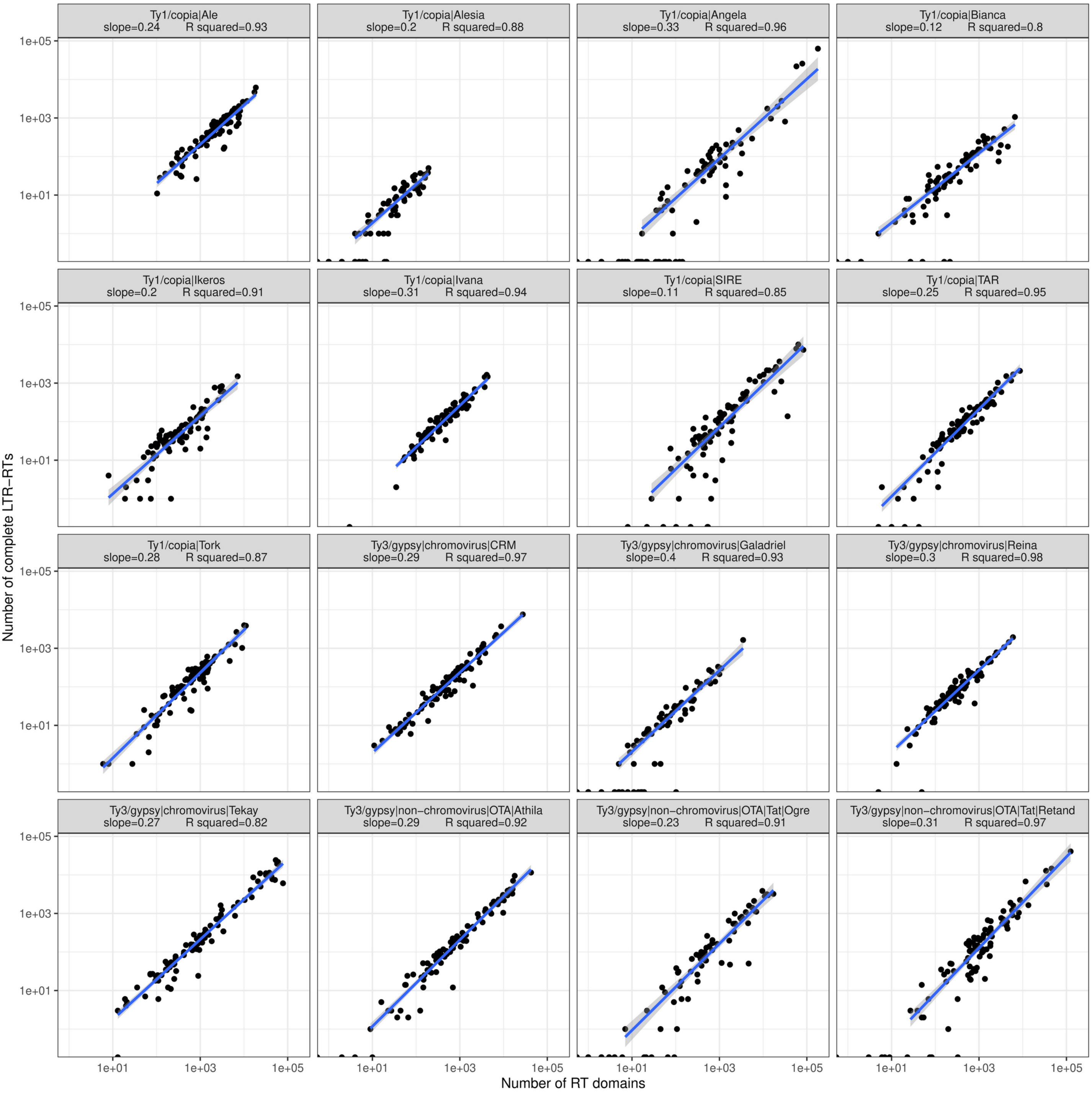


**Supplementary Figure S6.** Corelation between the numbers of RT domains detected by DANTE and numbers of elements identified by DANTE_LTR for individual lineages. Each point on the plot correspond to one species.
