## Supplementary Tables for "DANTE and DANTE_LTR: Lineage-centric annotation pipelines for long terminal repeat retrotransposons in plant genomes"

**Supplementary Table S1**. Analyzed species.

| Species | Assembly size [bp] | Number of detected elements | Order | Family | Genus |
|---|---|---|---|---|---|
| *Elaeis guineensis* | 1995563300 | 5217 | Arecales | Arecaceae | Elaeis |
| *Phoenix dactylifera* | 394279906 | 749 | Arecales | Arecaceae | Phoenix |
| *Asparagus officinalis* | 2128386688 | 11252 | Asparagales | Asparagaceae | Asparagus |
| *Apostasia shenzhenica* | 477914535 | 233 | Asparagales | Orchidaceae | Apostasia |
| *Dendrobium catenatum* | 1223162277 | 3949 | Asparagales | Orchidaceae | Dendrobium |
| *Gastrodia elata* | 1891193962 | 5633 | Asparagales | Orchidaceae | Gastrodia |
| *Phalaenopsis aphrodite* | 1494783072 | 4835 | Asparagales | Orchidaceae | Phalaenopsis |
| *Helianthus annuus* | 6005503774 | 15947 | Asterales | Asteraceae | Helianthus |
| *Lactuca sativa* | 4243749144 | 10424 | Asterales | Asteraceae | Lactuca |
| *Arabis alpina* | 623284120 | 3968 | Brassicales | Brassicaceae | Arabis |
| *Brassica nigra* | 725767990 | 2001 | Brassicales | Brassicaceae | Brassica |
| *Brassica juncea* | 1540181131 | 3366 | Brassicales | Brassicaceae | Brassica |
| *Camelina sativa* | 1120060535 | 3561 | Brassicales | Brassicaceae | Camelina |
| *Eutrema yunnanense* | 583539302 | 2751 | Brassicales | Brassicaceae | Eutrema |
| *Eutrema heterophyllum* | 543498760 | 2573 | Brassicales | Brassicaceae | Eutrema |
| *Thlaspi arvense* | 352756015 | 1926 | Brassicales | Brassicaceae | Thlaspi |
| *Carica papaya* | 479527162 | 2318 | Brassicales | Caricaceae | Carica |
| *Tarenaya hassleriana* | 379250372 | 965 | Brassicales | Cleomaceae | Tarenaya |
| *Carnegiea gigantea* | 635432796 | 4647 | Caryophyllales | Cactaceae | Carnegiea |
| *Drosera capensis* | 54040760 | 306 | Caryophyllales | Droseraceae | Drosera |
| *Lagenaria siceraria* | 562935328 | 1373 | Cucurbitales | Cucurbitaceae | Lagenaria |
| *Datisca glomerata* | 1183437095 | 6528 | Cucurbitales | Datiscaceae | Datisca |
| *Cicer reticulatum* | 602557920 | 1678 | Fabales | Fabaceae | Cicer |
| *Glycine soja* | 1761373269 | 9142 | Fabales | Fabaceae | Glycine |
| *Phaseolus vulgaris* | 118743908 | 955 | Fabales | Fabaceae | Phaseolus |
| *Vigna angularis* | 836937566 | 1824 | Fabales | Fabaceae | Vigna |
| *Quercus lobata* | 1665027898 | 5160 | Fagales | Fagaceae | Quercus |
| *Juglans microcarpa* | 927788423 | 4483 | Fagales | Juglandaceae | Juglans |
| *Juglans cathayensis* | 727243138 | 3232 | Fagales | Juglandaceae | Juglans |
| *Juglans sigillata* | 673935400 | 2812 | Fagales | Juglandaceae | Juglans |
| *Juglans nigra* | 758400756 | 3158 | Fagales | Juglandaceae | Juglans |
| *Juglans hindsii* | 895722710 | 3855 | Fagales | Juglandaceae | Juglans |
| *Pterocarya stenoptera* | 1018697693 | 4871 | Fagales | Juglandaceae | Pterocarya |
| *Morella rubra* | 589312148 | 1395 | Fagales | Myricaceae | Morella |
| *Boea hygrometrica* | 1605929661 | 13111 | Lamiales | Gesneriaceae | Dorcoceras |
| *Fraxinus excelsior* | 555779941 | 2591 | Lamiales | Oleaceae | Fraxinus |
| *Olea europaea* | 1622428655 | 12070 | Lamiales | Oleaceae | Olea |
| *Sesamum indicum* | 462295375 | 1043 | Lamiales | Pedaliaceae | Sesamum |
| *Cinnamomum kanehirae* | 1373242591 | 3107 | Laurales | Lauraceae | Cinnamomum |
| *Liriodendron chinense* | 3081011324 | 32654 | Magnoliales | Magnoliaceae | Liriodendron |
| *Hevea brasiliensis* | 2390335297 | 15787 | Malpighiales | Euphorbiaceae | Hevea |
| *Rhizophora apiculata* | 415930925 | 1400 | Malpighiales | Rhizophoraceae | Rhizophora |
| *Corchorus capsularis* | 100572175 | 615 | Malvales | Malvaceae | Corchorus |
| *Corchorus olitorius* | 105674004 | 593 | Malvales | Malvaceae | Corchorus |
| *Durio zibethinus* | 1318502838 | 5608 | Malvales | Malvaceae | Durio |
| *Gossypium barbadense* | 3181983891 | 13204 | Malvales | Malvaceae | Gossypium |
| *Gossypium hirsutum* | 3817177962 | 33731 | Malvales | Malvaceae | Gossypium |
| *Aquilaria agallocha* | 460600095 | 1693 | Malvales | Thymelaeaceae | Aquilaria |

| Species | Assembly size [bp] | Number of detected elements | Order | Family | Genus |
|---|---|---|---|---|---|
| *Punica granatum* | 496851204 | 1202 | Myrtales | Lythraceae | Punica |
| *Xerophyta viscosa* | 381974079 | 466 | Pandanales | Velloziaceae | Xerophyta |
| *Aegilops tauschii* | 6866156709 | 63171 | Poales | Poaceae | Aegilops |
| *Cenchrus americanus* | 3135920337 | 24592 | Poales | Poaceae | Cenchrus |
| *Dichanthelium oligosanthes* | 282081499 | 1310 | Poales | Poaceae | Dichanthelium |
| *Eleusine coracana* | 334717606 | 1861 | Poales | Poaceae | Eleusine |
| *Eragrostis tef* | 91996178 | 87 | Poales | Poaceae | Eragrostis |
| *Leersia perrieri* | 495447991 | 566 | Poales | Poaceae | Leersia |
| *Oryza glaberrima* | 79564300 | 203 | Poales | Poaceae | Oryza |
| *Oryza barthii* | 510062489 | 664 | Poales | Poaceae | Oryza |
| *Oryza rufipogon* | 622337164 | 933 | Poales | Poaceae | Oryza |
| *Oryza meridionalis* | 584950852 | 267 | Poales | Poaceae | Oryza |
| *Oryza glumaepatula* | 685821521 | 345 | Poales | Poaceae | Oryza |
| *Oryza punctata* | 753921303 | 2593 | Poales | Poaceae | Oryza |
| *Oryza sativa* | 782230770 | 3379 | Poales | Poaceae | Oryza |
| *Oryza sativa Japonica Group* | 374471240 | 2771 | Poales | Poaceae | Oryza |
| *Saccharum spontaneum* | 5827463848 | 31218 | Poales | Poaceae | Saccharum |
| *Triticum urartu* | 7261992288 | 56054 | Poales | Poaceae | Triticum |
| *Triticum turgidum* | 19390223261 | 166487 | Poales | Poaceae | Triticum |
| *Zea mays* | 2067864162 | 48276 | Poales | Poaceae | Zea |
| *Zizania latifolia* | 894966870 | 2775 | Poales | Poaceae | Zizania |
| *Eschscholzia californica* | 692086867 | 1808 | Ranunculales | Papaveraceae | Eschscholzia |
| *Macleaya cordata* | 451230647 | 1447 | Ranunculales | Papaveraceae | Macleaya |
| *Papaver somniferum* | 5052401771 | 18544 | Ranunculales | Papaveraceae | Papaver |
| *Cannabis sativa* | 1120060535 | 3561 | Rosales | Cannabaceae | Cannabis |
| *Humulus lupulus* | 447962224 | 3010 | Rosales | Cannabaceae | Humulus |
| *Parasponia andersonii* | 837326715 | 5331 | Rosales | Cannabaceae | Parasponia |
| *Trema orientalis* | 634523492 | 2374 | Rosales | Cannabaceae | Trema |
| *Ficus carica* | 42269757 | 67 | Rosales | Moraceae | Ficus |
| *Morus notabilis* | 440538766 | 1786 | Rosales | Moraceae | Morus |
| *Ziziphus jujuba* | 710111727 | 2532 | Rosales | Rhamnaceae | Ziziphus |
| *Prunus yedoensis* | 279012417 | 1112 | Rosales | Rosaceae | Prunus |
| *Prunus mume* | 406440083 | 772 | Rosales | Rosaceae | Prunus |
| *Rosa multiflora* | 554805873 | 2721 | Rosales | Rosaceae | Rosa |
| *Boehmeria nivea* | 465212001 | 1224 | Rosales | Urticaceae | Boehmeria |
| *Santalum album* | 327459801 | 490 | Santalales | Santalaceae | Santalum |
| *Kalanchoe fedtschenkoi* | 391878075 | 526 | Saxifragales | Crassulaceae | Kalanchoe |
| *Jaltomata sinuosa* | 2467117205 | 24559 | Solanales | Solanaceae | Jaltomata |
| *Nicotiana tabacum* | 3592401316 | 33951 | Solanales | Solanaceae | Nicotiana |
| *Nicotiana tomentosiformis* | 1859114963 | 16337 | Solanales | Solanaceae | Nicotiana |
| *Nicotiana sylvestris* | 2220944715 | 20408 | Solanales | Solanaceae | Nicotiana |
| *Nicotiana obtusifolia* | 1589554527 | 13865 | Solanales | Solanaceae | Nicotiana |
| *Nicotiana attenuata* | 3691367980 | 25093 | Solanales | Solanaceae | Nicotiana |
| *Solanum pimpinellifolium* | 609973449 | 4443 | Solanales | Solanaceae | Solanum |
| *Musa itinerans* | 210832024 | 747 | Zingiberales | Musaceae | Musa |

**Supplementary Table S2**. The number of elements identified in rice (*O. sativa)* genome by DANTE_LTR

| Lineage | Number of elements | Element mean length [bp] | LTR mean length [bp] | Number of elements with PBS & TSD | Number of elements with PBS only | Number of elements with TSD only |
|---|---|---|---|---|---|---|
| Ty1/copia\|Ale | 155 | 4790 | 173 | 88 | 38 | 6 |
| Ty1/copia\|Angela | 19 | 7671 | 1388 | 5 | 5 | 0 |
| Ty1/copia\|Bianca | 33 | 7631 | 278 | 14 | 12 | 2 |
| Ty1/copia\|Ikeros | 70 | 6552 | 490 | 48 | 9 | 0 |
| Ty1/copia\|Ivana | 119 | 5208 | 391 | 82 | 16 | 3 |
| Ty1/copia\|SIRE | 151 | 9398 | 1215 | 75 | 34 | 6 |
| Ty1/copia\|TAR | 298 | 6424 | 958 | 114 | 59 | 59 |
| Ty1/copia\|Tork | 32 | 5070 | 267 | 20 | 5 | 2 |
| Ty3/gypsy\|chromovirus\|CRM | 49 | 7766 | 908 | 28 | 9 | 3 |
| Ty3/gypsy\|chromovirus\|Reina | 135 | 5506 | 353 | 81 | 7 | 34 |
| Ty3/gypsy\|chromovirus\|Tekay | 752 | 12000 | 2989 | 342 | 169 | 19 |
| Ty3/gypsy\|non-chromovirus\|OTA\|Athila | 23 | 9714 | 1169 | 0 | 3 | 8 |
| Ty3/gypsy\|non-chromovirus\|OTA\|Tat\|Ogre | 162 | 13161 | 1190 | 10 | 16 | 58 |
| Ty3/gypsy\|non-chromovirus\|OTA\|Tat\|Retand | 773 | 11613 | 579 | 595 | 68 | 29 |

**Supplementary Table S3**. Performance metrics of annotation methods. Best results for each method and annotation type are highlighted

| | | | Metric | | | |
|---|---|---|---|---|---|---|
| **Annotation** | **Species** | **Method** | **TP** | **FN** | **FP** | **TN** |
| Complete elements | **Rice** | **DANTE_LTR** | 26872541 | 60616328 | 383797 | 286598574 |
| | | **Inpactor2** | 13321780 | 74215050 | 224438 | 286709972 |
| | | **EDTA** | 19804461 | 67744438 | 265619 | 286656722 |
| | **Maize** | **DANTE_LTR** | 456893021 | 1060485719 | 30349077 | 520136345 |
| | | **Inpactor2** | 289978568 | 1226746184 | 28769546 | 522369864 |
| | | **EDTA** | 217168457 | 1310655957 | 17887420 | 522152328 |
| Library based | **Rice** | **DANTE_LTR** | 65625866 | 21865533 | 3803586 | 283176255 |
| | | **Inpactor2** | 76890588 | 7429910 | 15590751 | 274559991 |
| | | **EDTA** | 80131281 | 2883204 | 23554064 | 267902691 |
| | **Maize** | **DANTE_LTR** | 1212898818 | 229765047 | 143975459 | 481224838 |
| | | **Inpactor2** | 1267952352 | 77283441 | 350200119 | 372428250 |
| | | **EDTA** | 1234901908 | 212942428 | 214092198 | 405927628 |

| | | | Metric | | | | | |
|---|---|---|---|---|---|---|---|---|
| **Annotation** | **Species** | **Method** | **precision** | **sensitivity** | **specificity** | **FDR** | **F1** | **accuracy** |
| Complete elements | **Rice** | **DANTE_LTR** | 0.9859 | **0.3072** | 0.9987 | 0.0141 | **0.4684** | **0.8371** |
| | | **Inpactor2** | 0.9834 | 0.1522 | **0.9992** | 0.0166 | 0.2636 | 0.8012 |
| | | **EDTA** | **0.9868** | 0.2262 | 0.9991 | **0.0132** | 0.3680 | 0.8184 |
| | **Maize** | **DANTE_LTR** | **0.9377** | **0.3011** | 0.9449 | **0.0623** | **0.4558** | **0.4725** |
| | | **Inpactor2** | 0.9097 | 0.1912 | 0.9478 | 0.0903 | 0.3160 | 0.3928 |
| | | **EDTA** | 0.9239 | 0.1421 | **0.9669** | 0.0761 | 0.2464 | 0.3575 |
| Library based | **Rice** | **DANTE_LTR** | **0.9452** | 0.7501 | **0.9867** | **0.0548** | 0.8364 | 0.9315 |
| | | **Inpactor2** | 0.8314 | 0.9119 | 0.9463 | 0.1686 | **0.8698** | **0.9385** |
| | | **EDTA** | 0.7728 | **0.9653** | 0.9192 | 0.2272 | 0.8584 | 0.9294 |
| | **Maize** | **DANTE_LTR** | **0.8939** | 0.8407 | **0.7697** | **0.1061** | **0.8665** | **0.8193** |
| | | **Inpactor2** | 0.7836 | **0.9426** | 0.5154 | 0.2164 | 0.8557 | 0.7933 |
| | | **EDTA** | 0.8522 | 0.8529 | 0.6547 | 0.1478 | 0.8526 | 0.7935 |
