## Supplementary File S1 for "DANTE and DANTE_LTR: Lineage-centric annotation pipelines for long terminal repeat retrotransposons in plant genomes": summary.html

DANTE LTR Summary


DANTE\_LTR Summary

Number of elements

TE structure summary and LTR identity

Ty1/copia|Ale

Ty1/copia|Angela

Ty1/copia|Bianca

Ty1/copia|Ikeros

Ty1/copia|Ivana

Ty1/copia|SIRE

Ty1/copia|TAR

Ty1/copia|Tork

Ty3/gypsy|chromovirus|CRM

Ty3/gypsy|chromovirus|Reina

Ty3/gypsy|chromovirus|Tekay

Ty3/gypsy|non-chromovirus|OTA|Athila

Ty3/gypsy|non-chromovirus|OTA|Tat|Ogre

Ty3/gypsy|non-chromovirus|OTA|Tat|Retand

### DANTE LTR Summary

| Lineage | Number of elements | Element mean length   [bp] | LTR mean length   [bp] | Number of elements  with PBS & TSD | Number of elements  with PBS only | Number of elements  with TSD only |
| --- | --- | --- | --- | --- | --- | --- |
| Ty1/copia\|Ale | 675 | 5363 | 141 | 307 | 184 | 61 |
| Ty1/copia\|Angela | 382 | 7804 | 1274 | 216 | 79 | 5 |
| Ty1/copia\|Bianca | 118 | 7777 | 198 | 82 | 12 | 5 |
| Ty1/copia\|Ikeros | 23 | 7199 | 559 | 15 | 3 | 1 |
| Ty1/copia\|Ivana | 314 | 5829 | 393 | 193 | 55 | 9 |
| Ty1/copia\|SIRE | 19283 | 9198 | 1233 | 9822 | 4079 | 355 |
| Ty1/copia\|TAR | 776 | 6380 | 928 | 107 | 112 | 249 |
| Ty1/copia\|Tork | 5 | 4734 | 296 | 2 | 2 | 0 |
| Ty3/gypsy\|chromovirus\|CRM | 536 | 6858 | 569 | 129 | 255 | 18 |
| Ty3/gypsy\|chromovirus\|Reina | 765 | 5302 | 379 | 397 | 70 | 187 |
| Ty3/gypsy\|chromovirus\|Tekay | 9121 | 10331 | 1955 | 1843 | 3493 | 231 |
| Ty3/gypsy\|non-chromovirus\|OTA\|Athila | 1 | 11313 | 1441 | 0 | 0 | 0 |
| Ty3/gypsy\|non-chromovirus\|OTA\|Tat\|Ogre | 373 | 9269 | 687 | 30 | 15 | 204 |
| Ty3/gypsy\|non-chromovirus\|OTA\|Tat\|Retand | 15904 | 11992 | 632 | 9860 | 2160 | 402 |

#### Number of elements

#### TE structure summary and LTR identity

#### Ty1/copia|Ale summary

#### Ale : Structure of individual elements

#### Ty1/copia|Angela summary

#### Angela : Structure of individual elements

#### Ty1/copia|Bianca summary

#### Bianca : Structure of individual elements

#### Ty1/copia|Ikeros summary

#### Ikeros : Structure of individual elements

#### Ty1/copia|Ivana summary

#### Ivana : Structure of individual elements

#### Ty1/copia|SIRE summary

#### SIRE : Structure of individual elements

#### Ty1/copia|TAR summary

#### TAR : Structure of individual elements

#### Ty1/copia|Tork summary

#### Tork : Structure of individual elements

#### Ty3/gypsy|chromovirus|CRM summary

#### CRM : Structure of individual elements

#### Ty3/gypsy|chromovirus|Reina summary

#### Reina : Structure of individual elements

#### Ty3/gypsy|chromovirus|Tekay summary

#### Tekay : Structure of individual elements

#### Ty3/gypsy|non-chromovirus|OTA|Athila summary

#### Athila : Structure of individual elements

#### Ty3/gypsy|non-chromovirus|OTA|Tat|Ogre summary

#### Ogre : Structure of individual elements

#### Ty3/gypsy|non-chromovirus|OTA|Tat|Retand summary

#### Retand : Structure of individual elements
