## Supplementary figures and images for "DANTE and DANTE_LTR: Lineage-centric annotation pipelines for long terminal repeat retrotransposons in plant genomes"

### all_elements.png

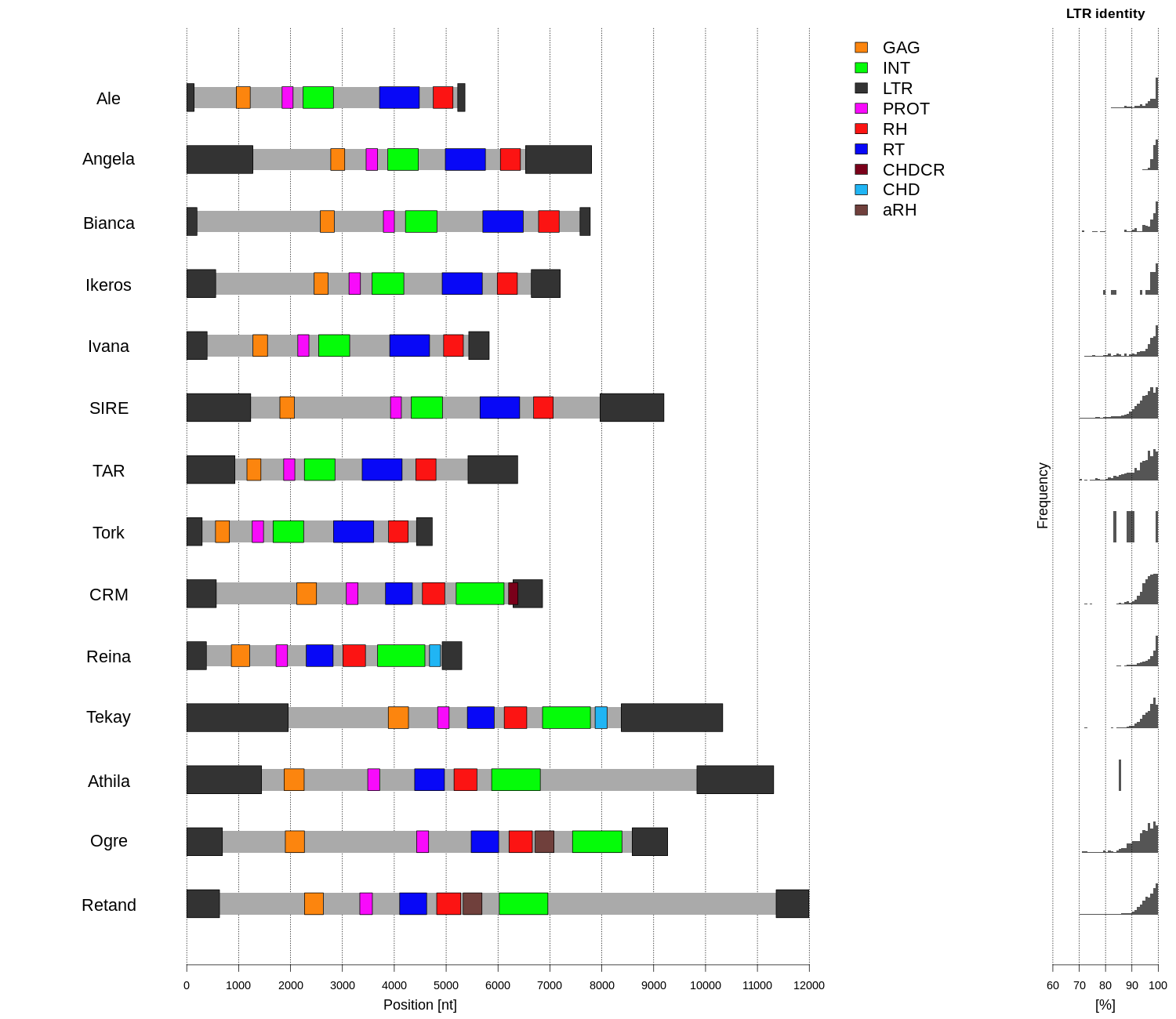

### all_elements.png

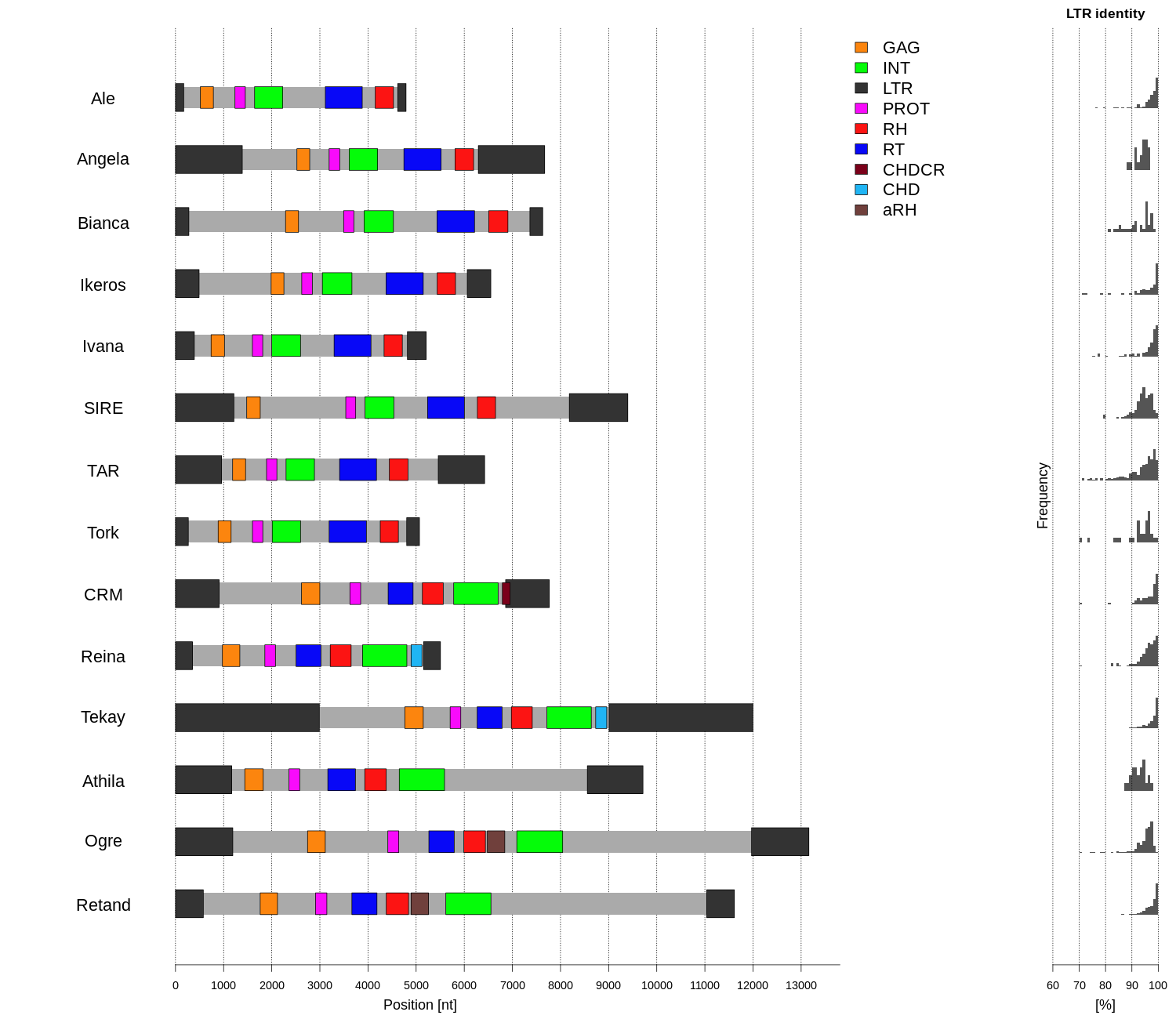

### Class_I_LTR_Ty1_copia_Ale_structure.png

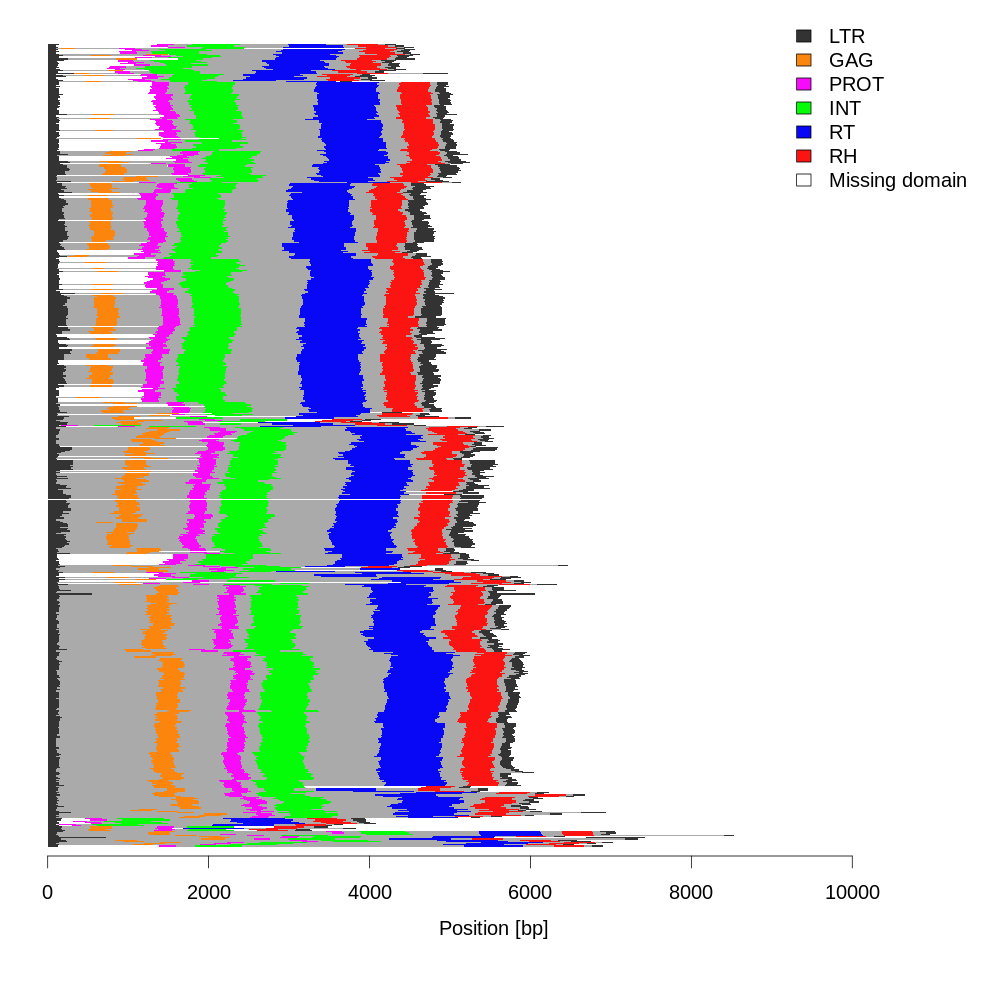

### Class_I_LTR_Ty1_copia_Ale_structure.png

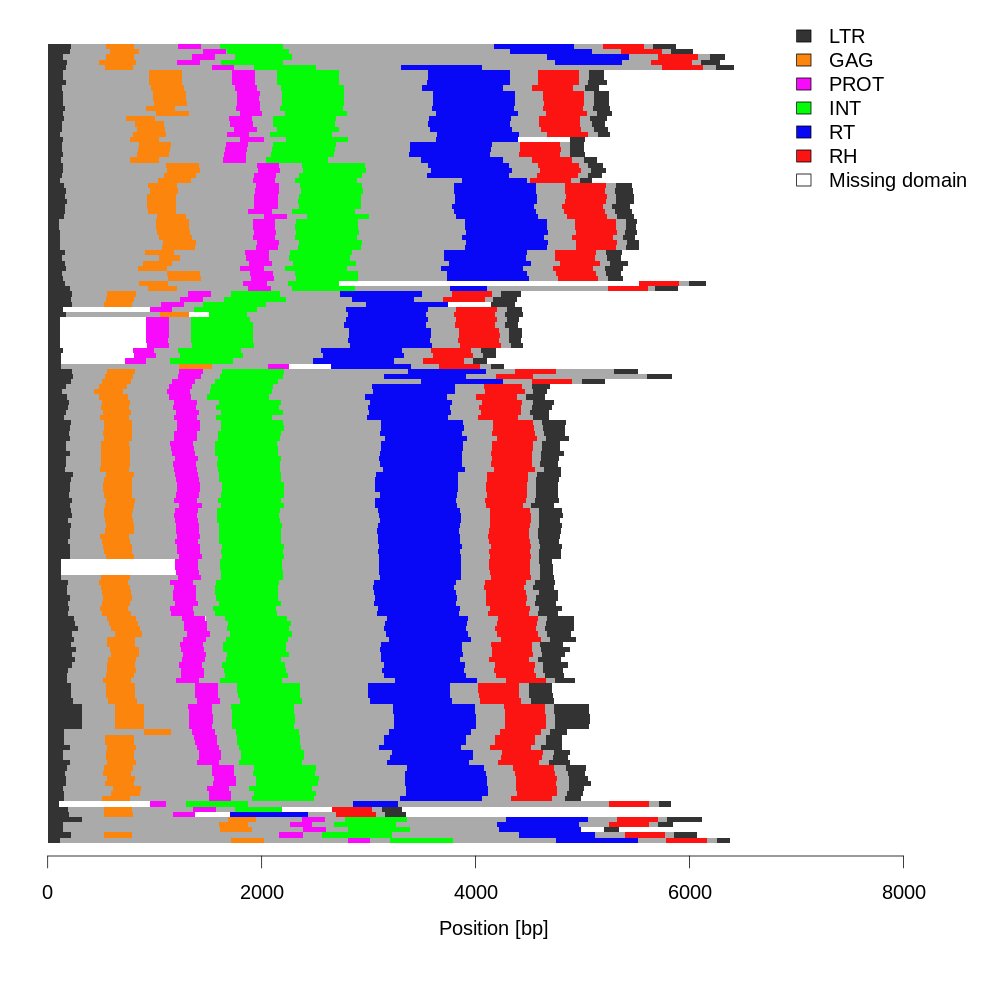

### Class_I_LTR_Ty1_copia_Ale_summary.png

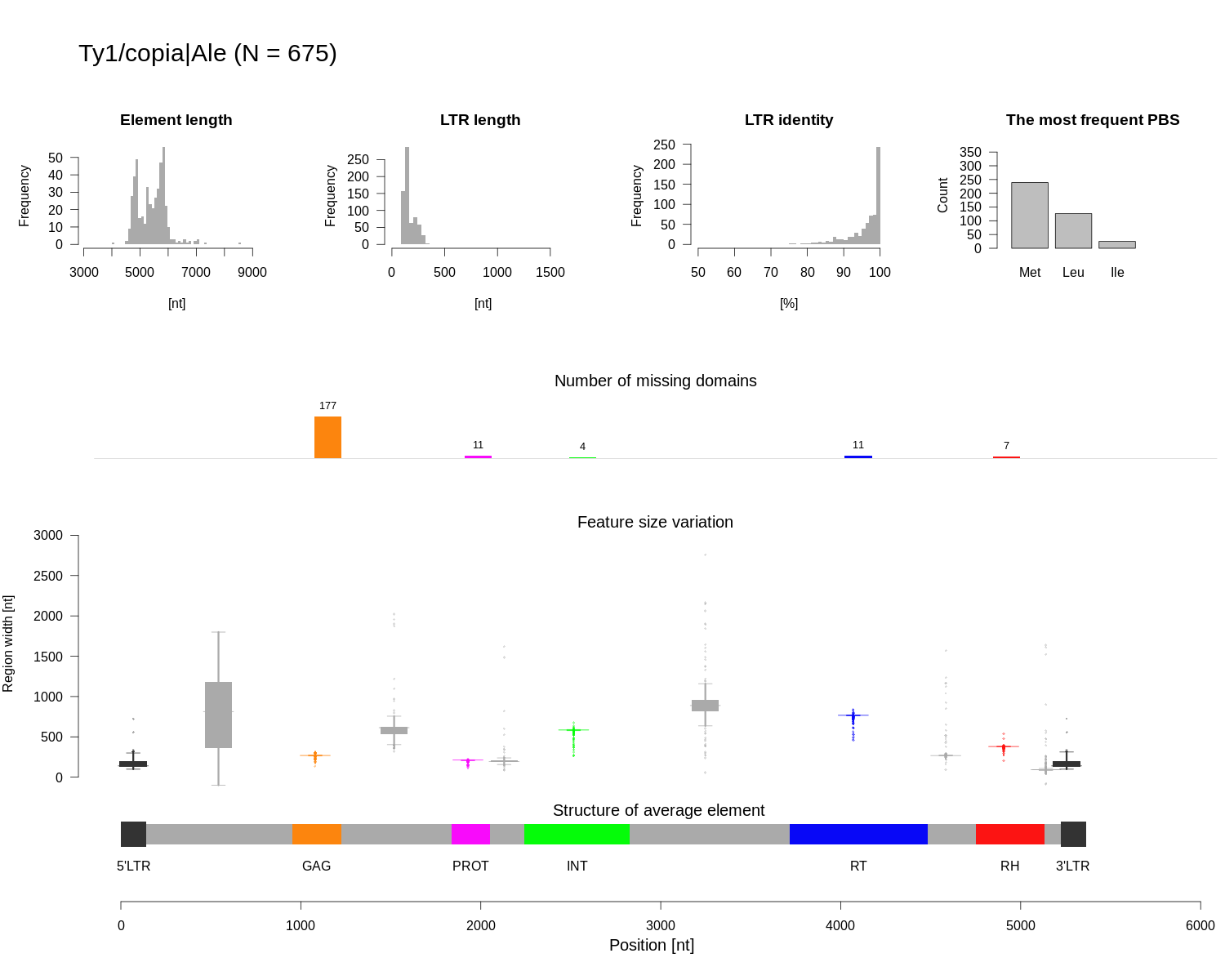

### Class_I_LTR_Ty1_copia_Ale_summary.png

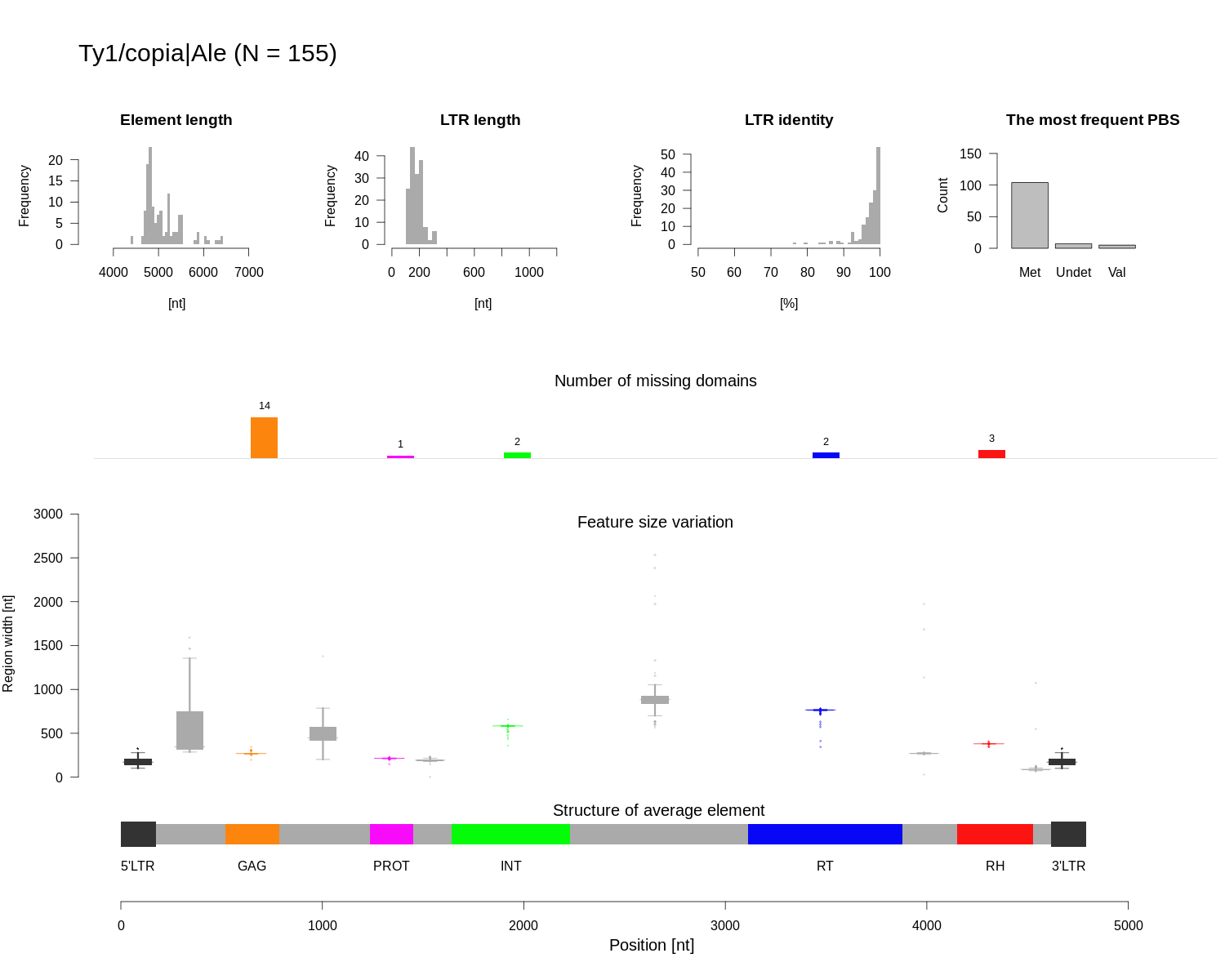

### Class_I_LTR_Ty1_copia_Angela_structure.png

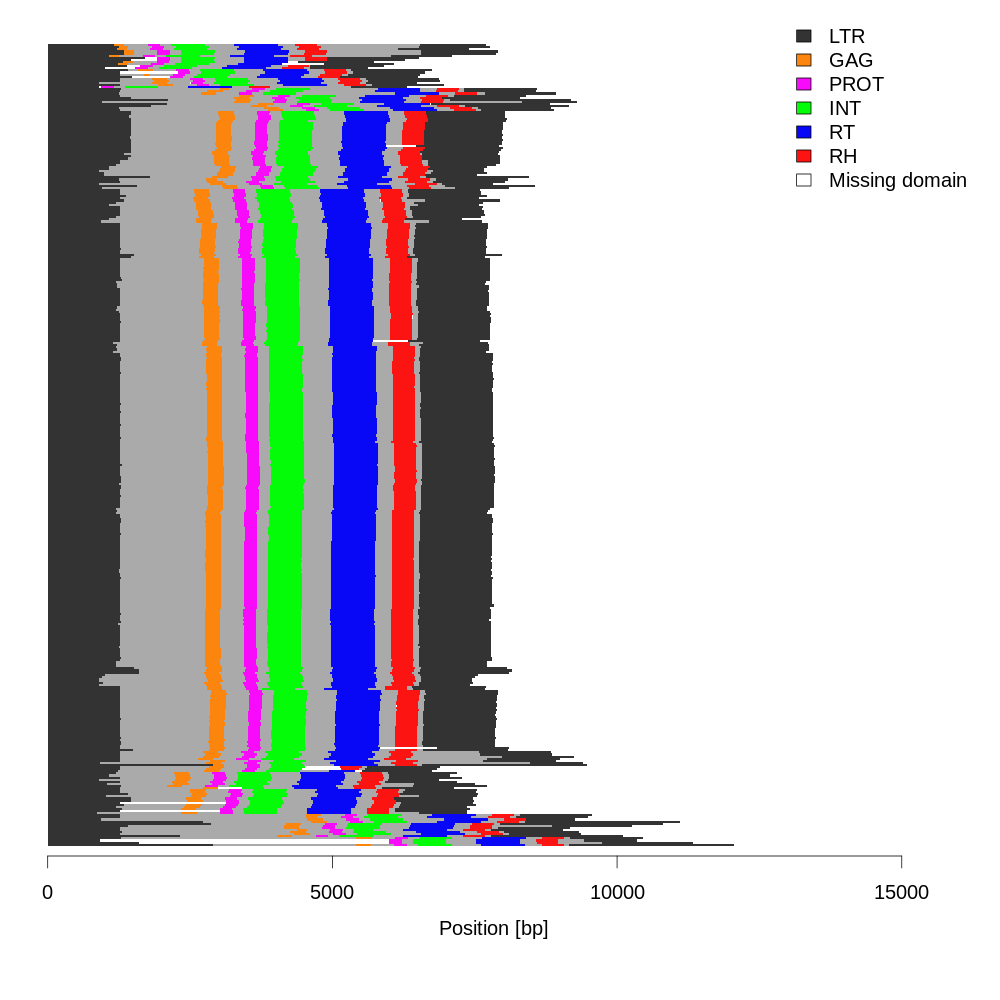

### Class_I_LTR_Ty1_copia_Angela_structure.png

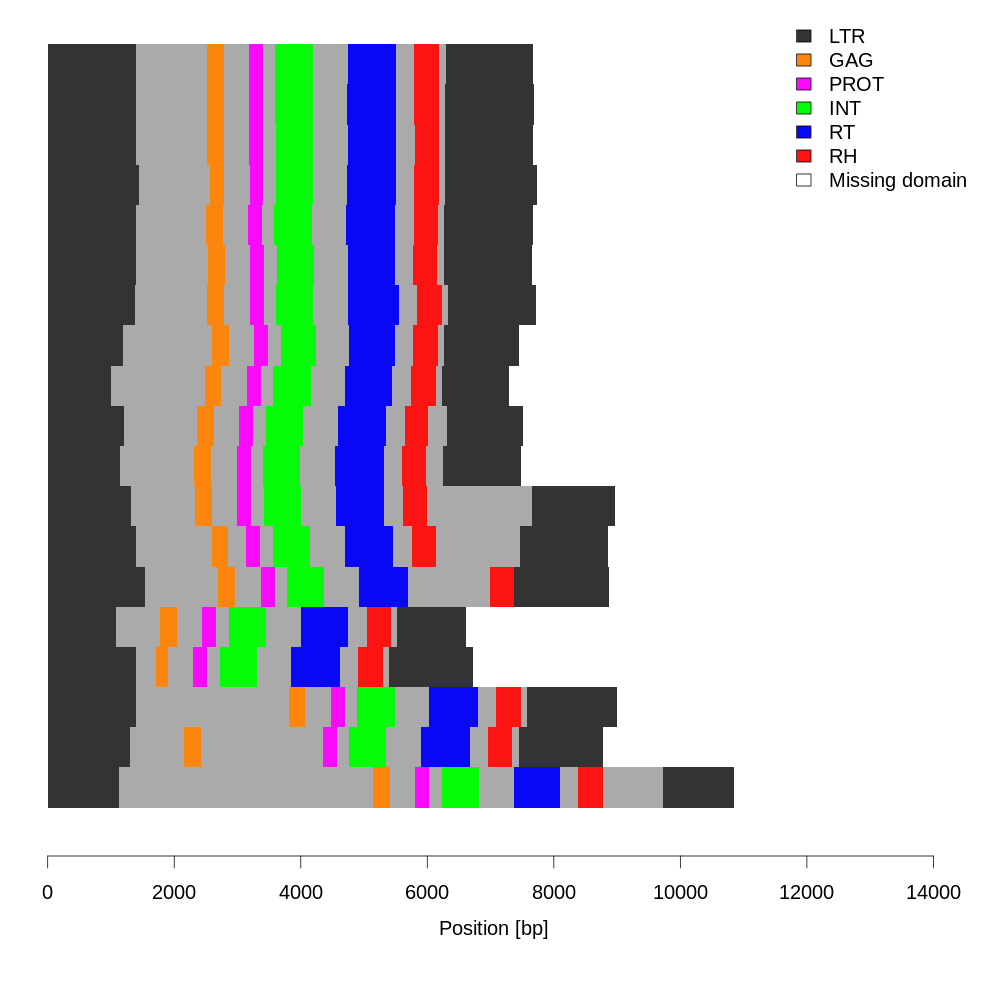

### Class_I_LTR_Ty1_copia_Angela_summary.png

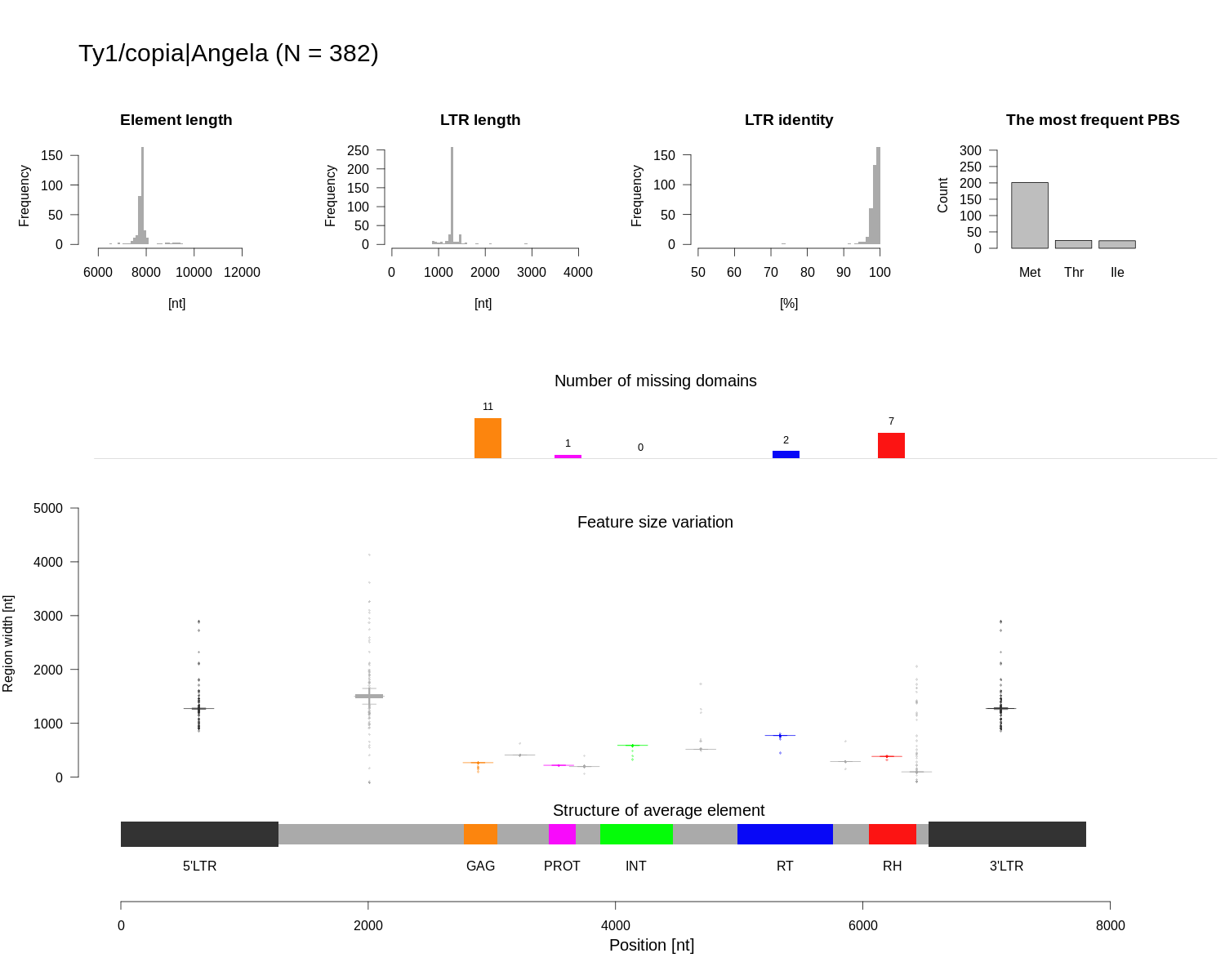

### Class_I_LTR_Ty1_copia_Angela_summary.png

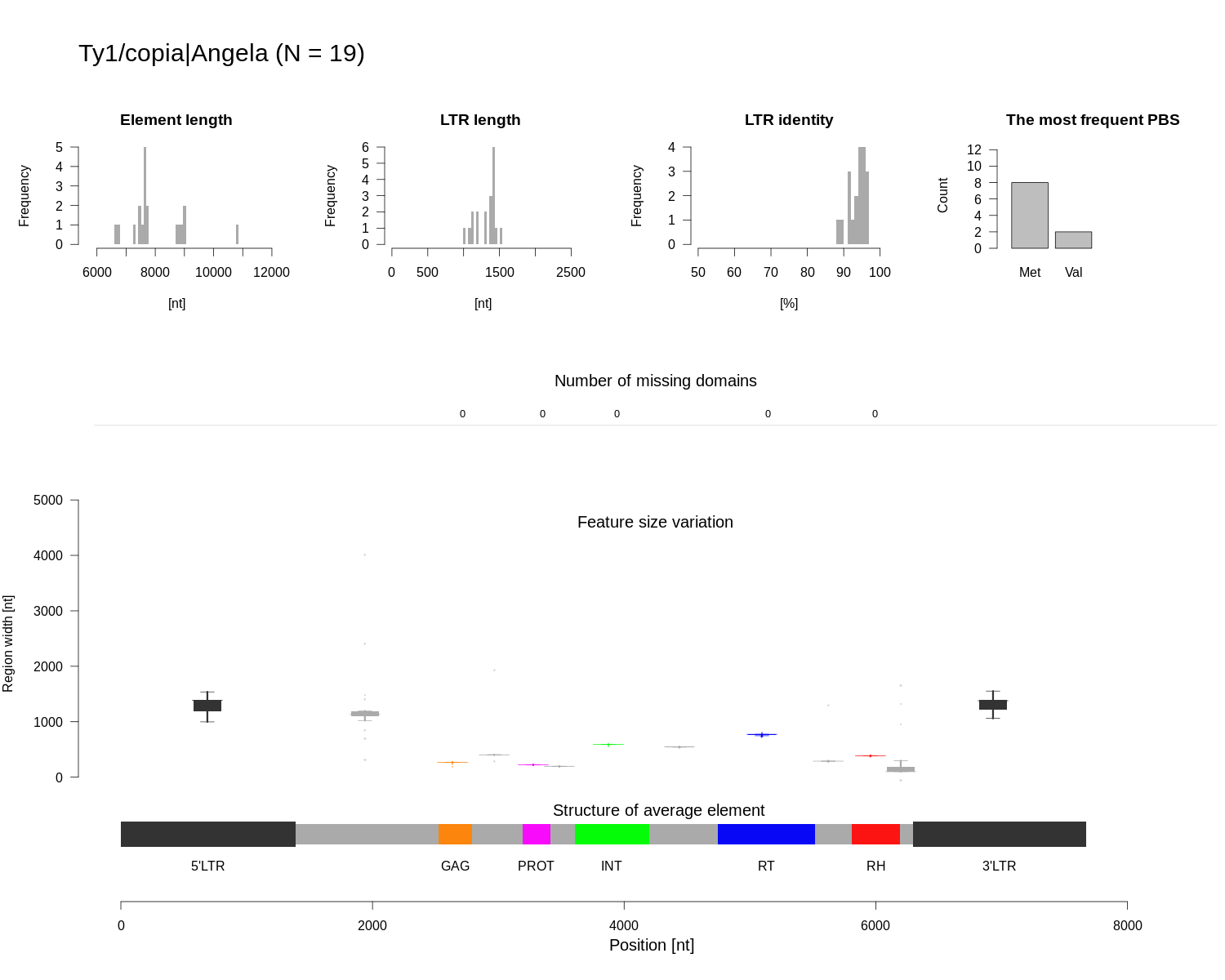

### Class_I_LTR_Ty1_copia_Bianca_structure.png

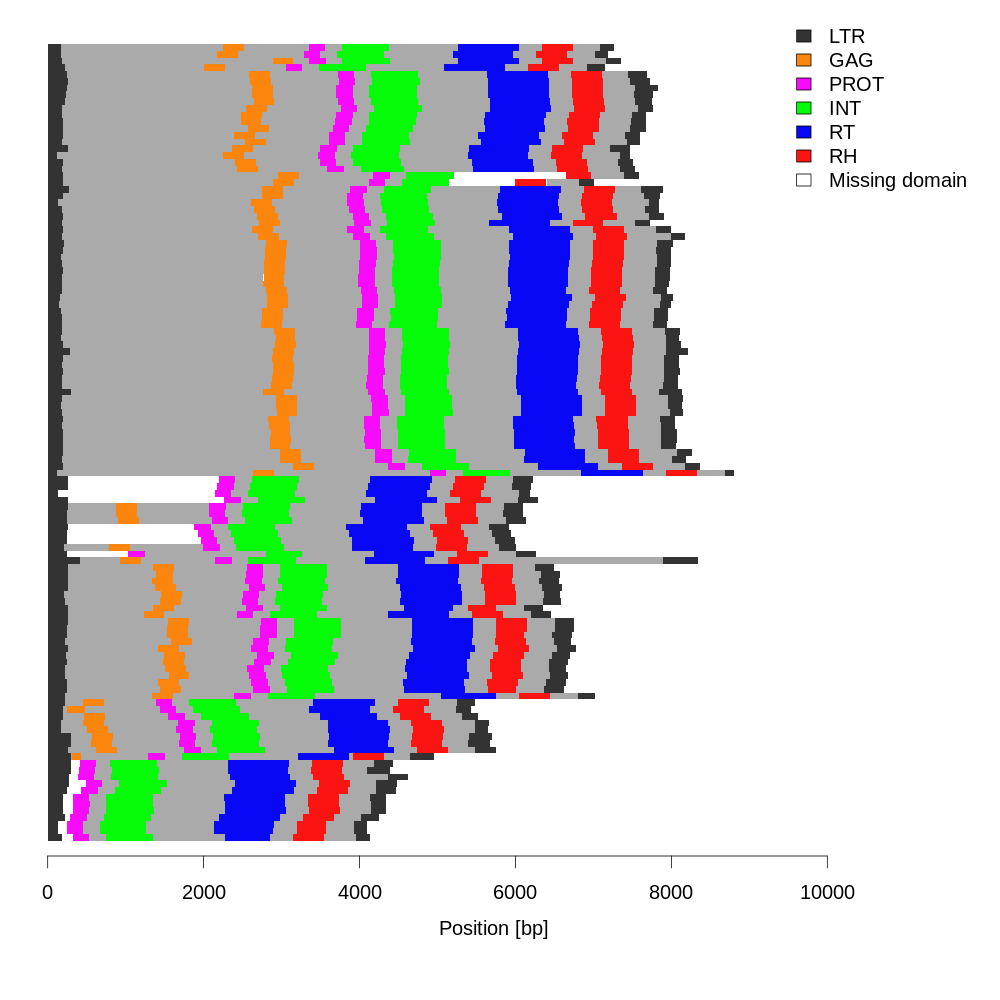

### Class_I_LTR_Ty1_copia_Bianca_structure.png

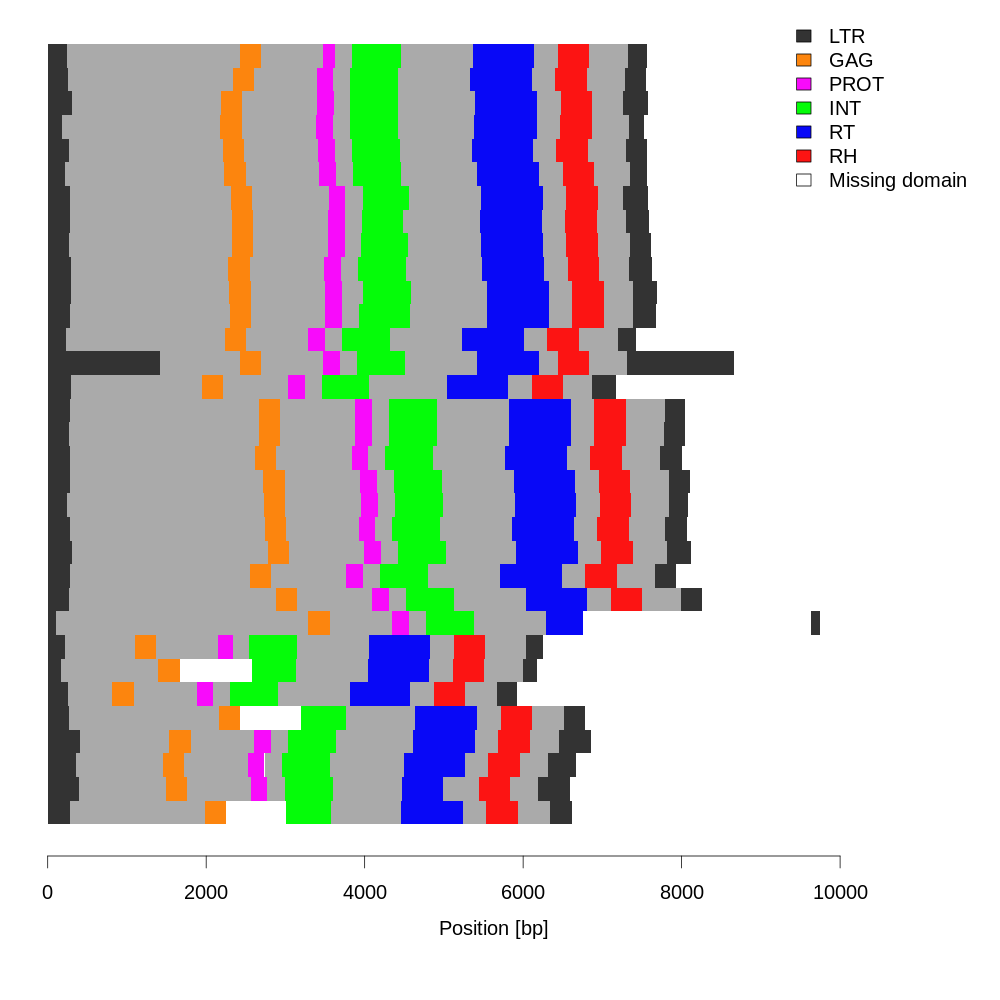

### Class_I_LTR_Ty1_copia_Bianca_summary.png

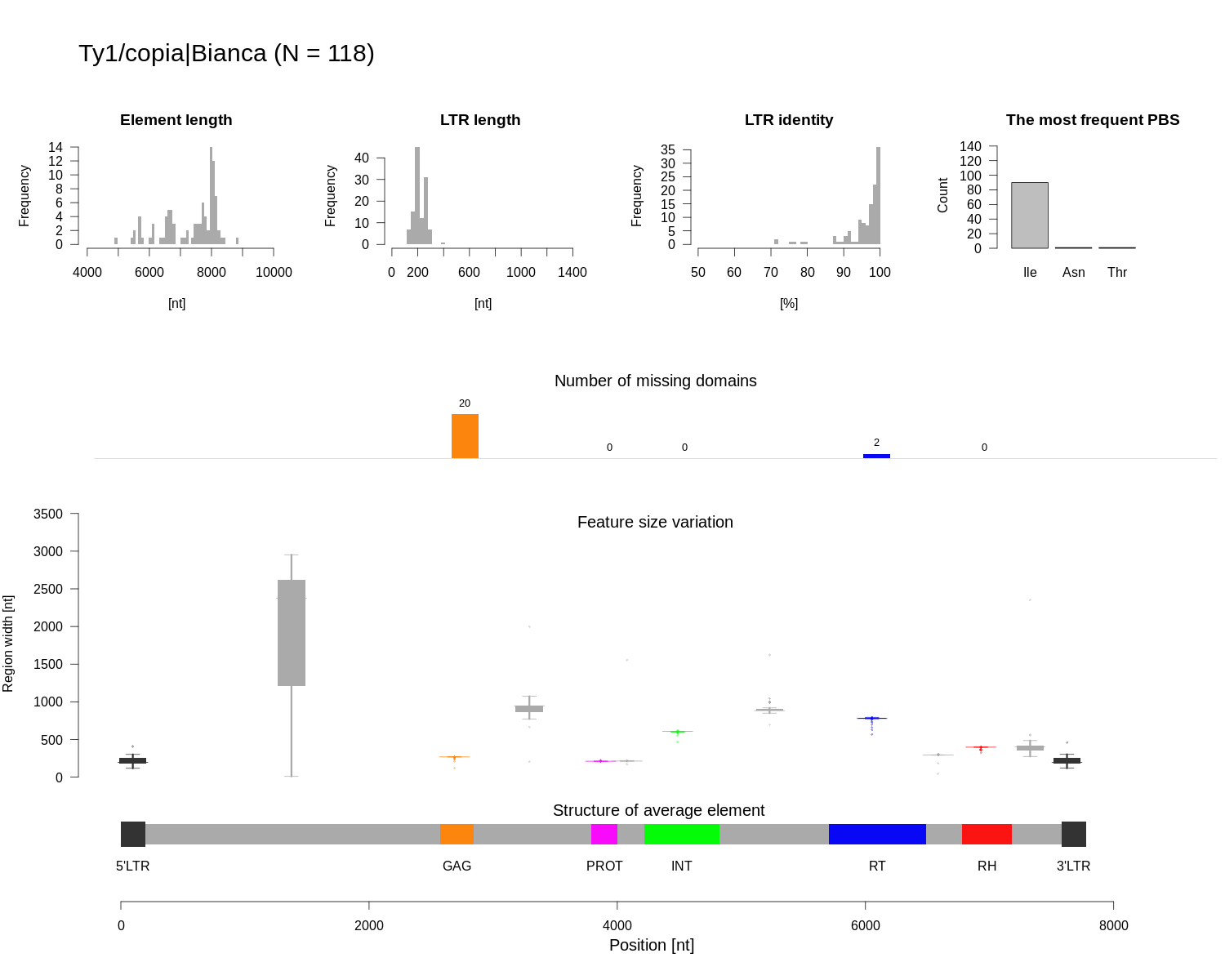

### Class_I_LTR_Ty1_copia_Bianca_summary.png

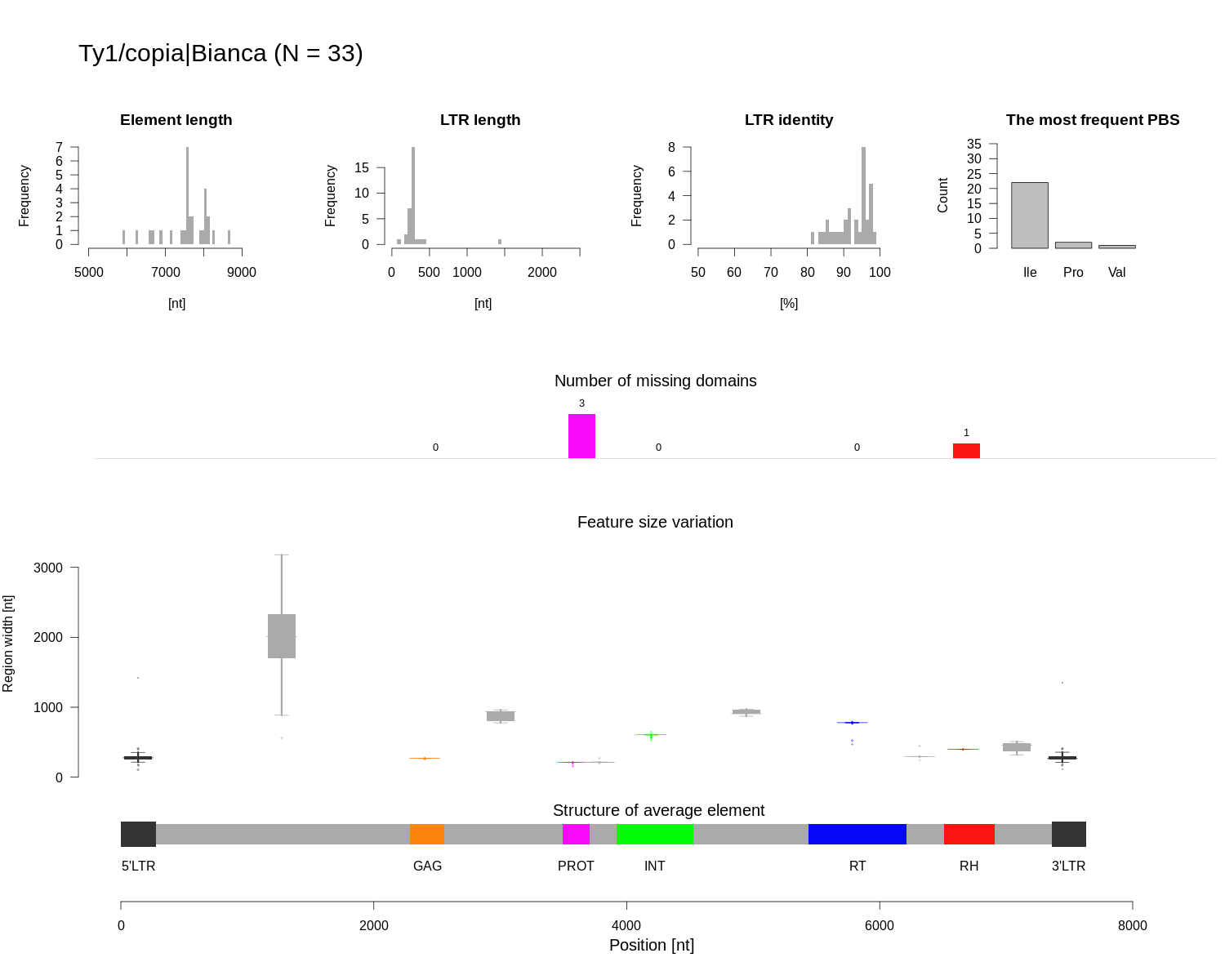

### Class_I_LTR_Ty1_copia_Ikeros_structure.png

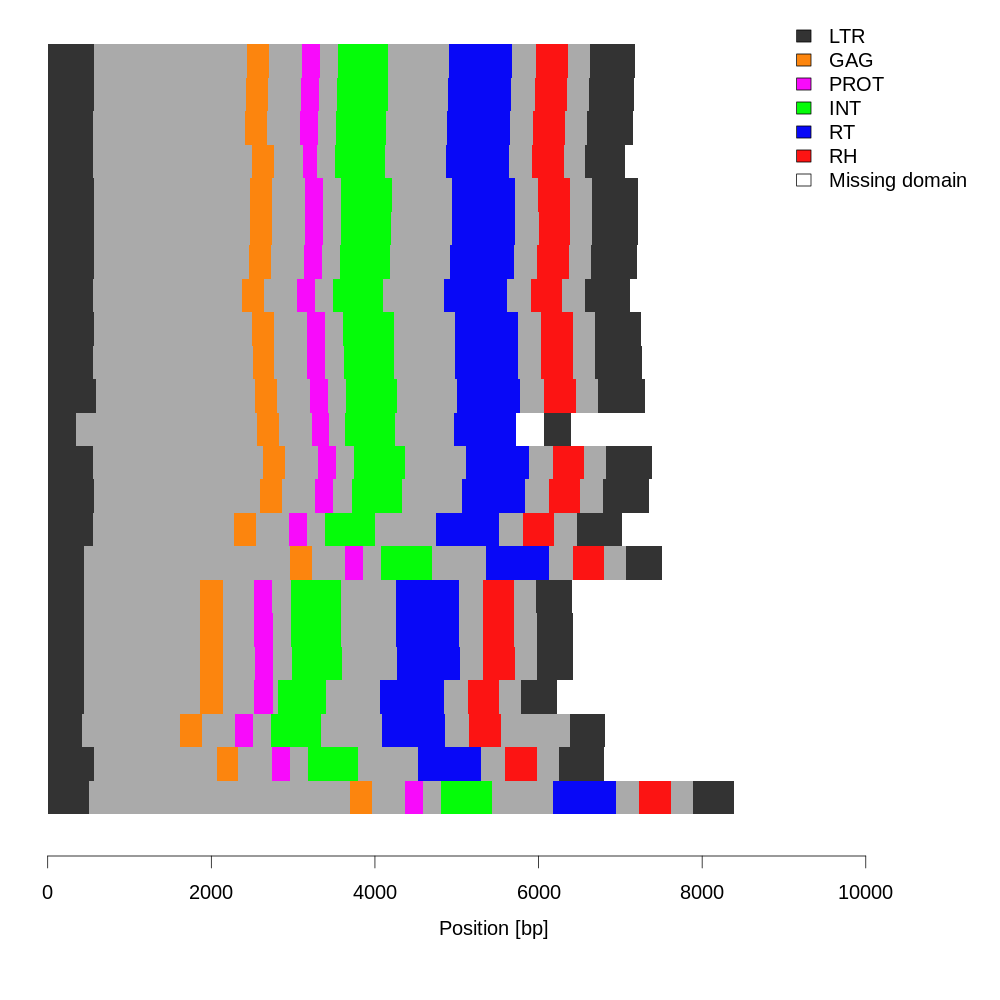

### Class_I_LTR_Ty1_copia_Ikeros_structure.png

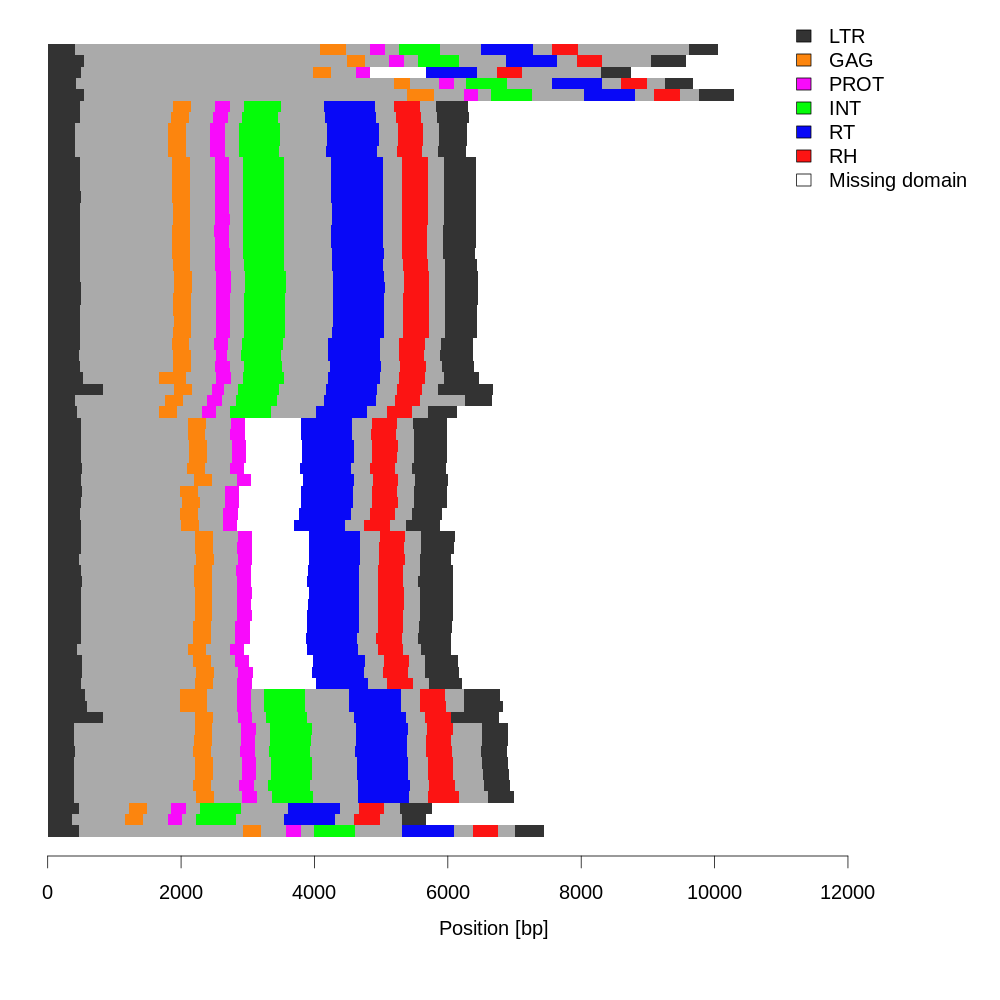

### Class_I_LTR_Ty1_copia_Ikeros_summary.png

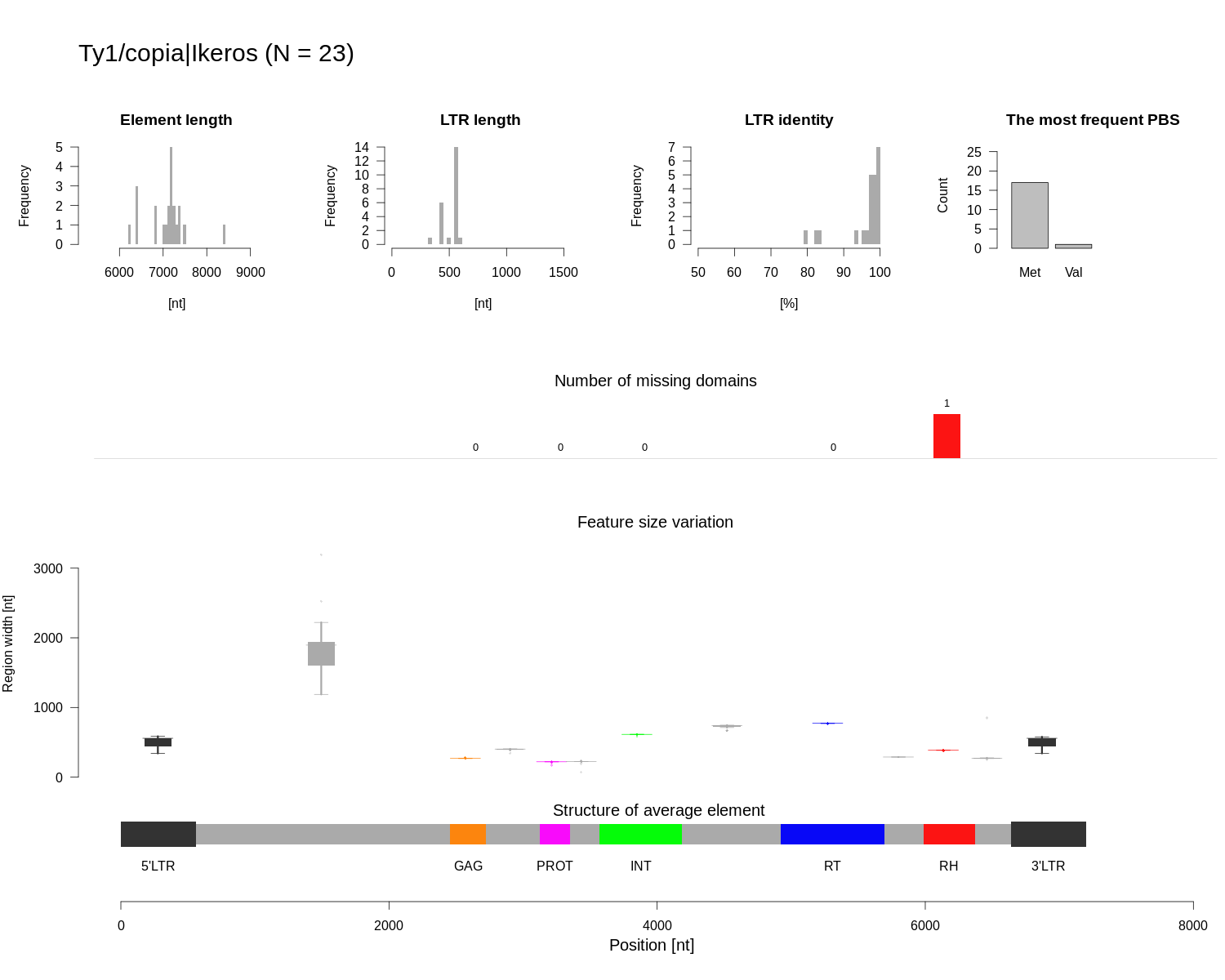

### Class_I_LTR_Ty1_copia_Ikeros_summary.png

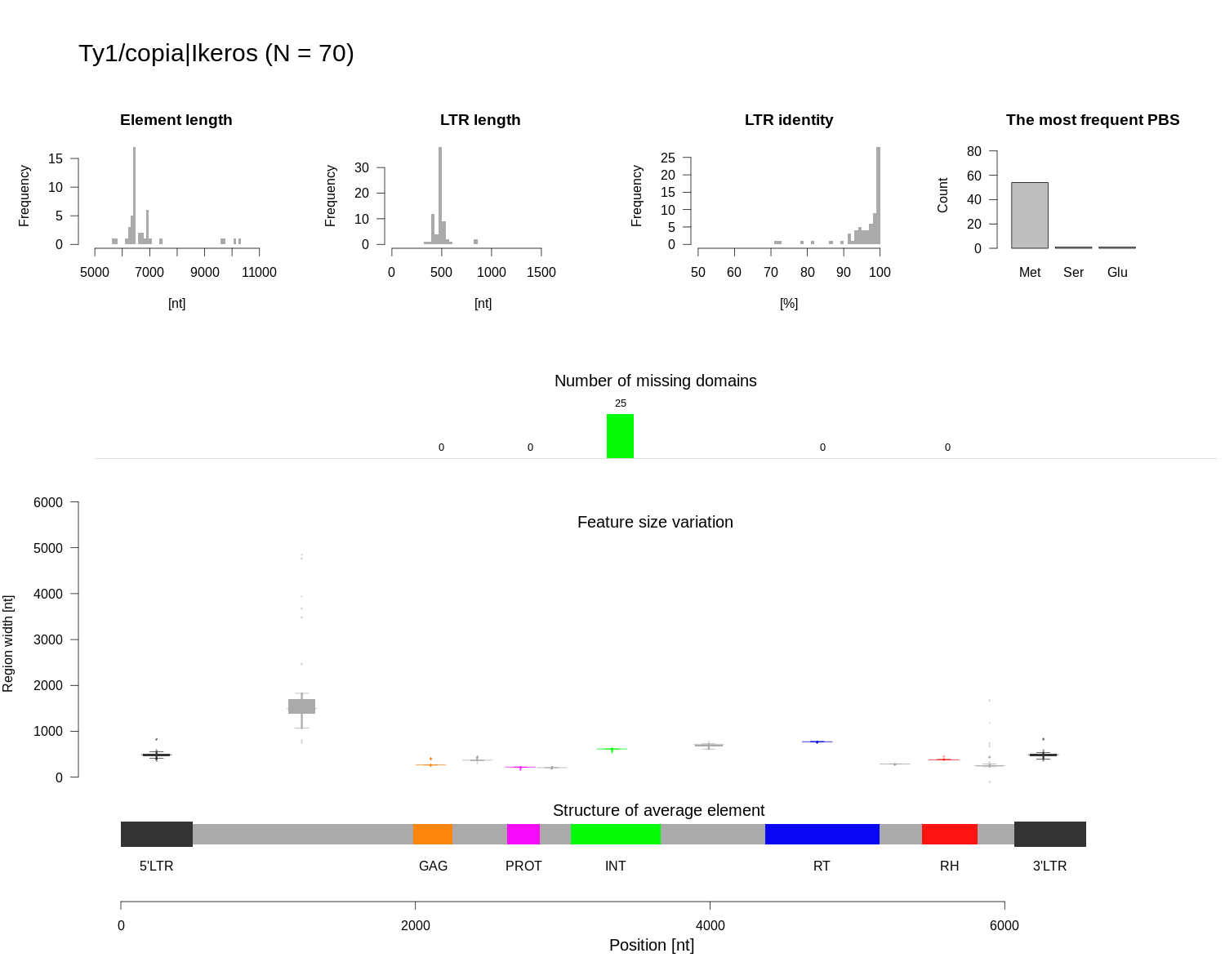

### Class_I_LTR_Ty1_copia_Ivana_structure.png

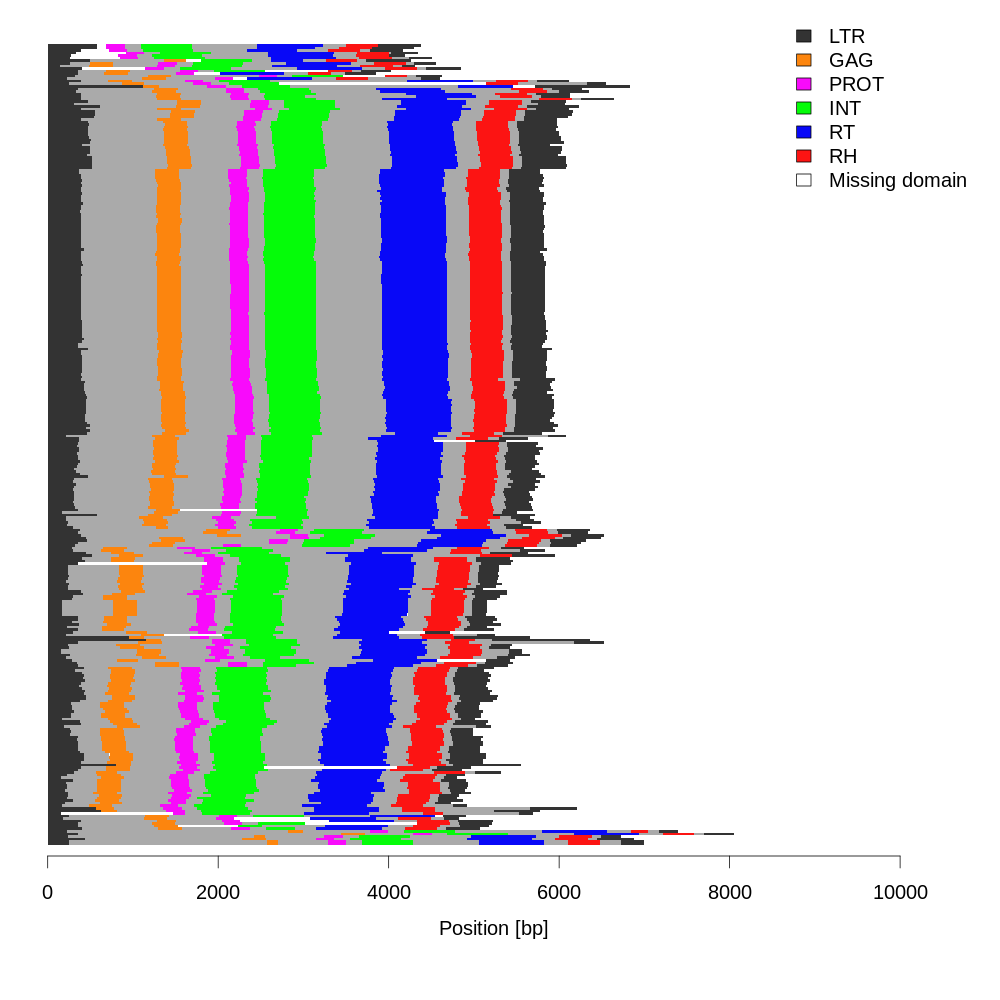

### Class_I_LTR_Ty1_copia_Ivana_structure.png

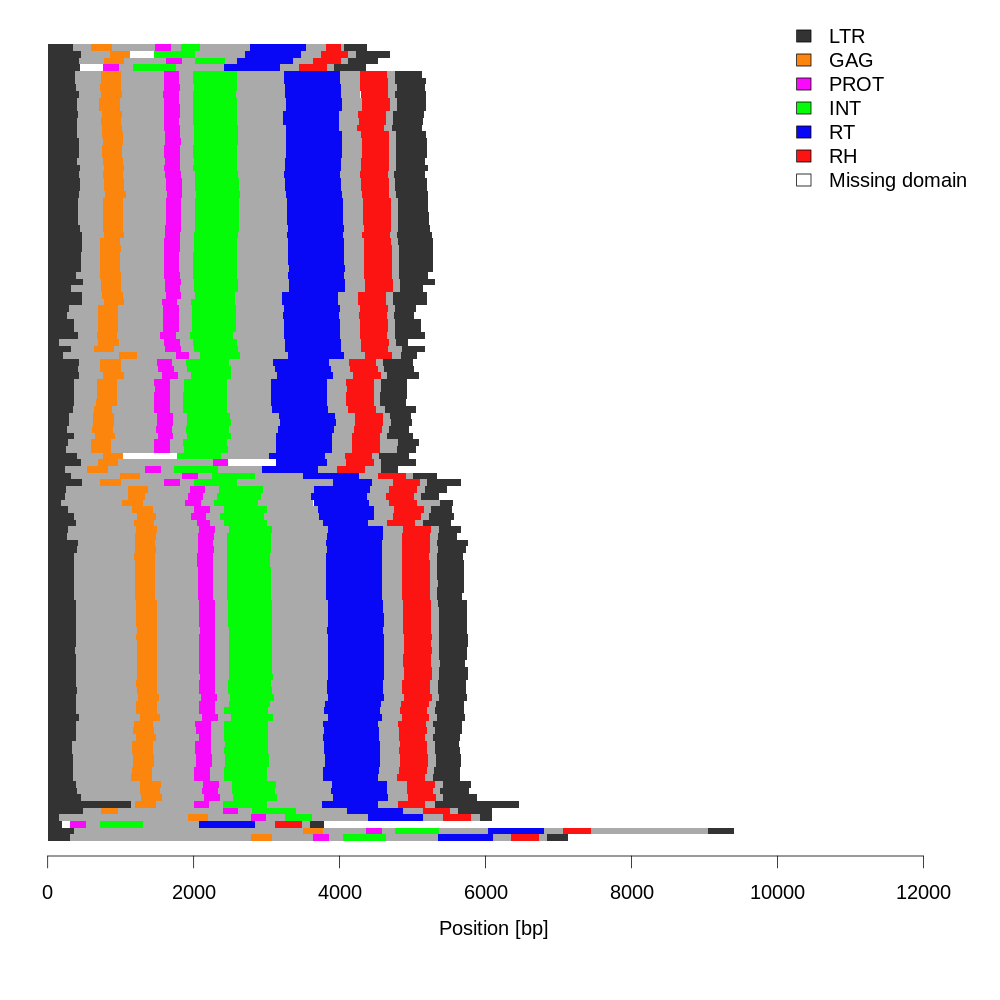

### Class_I_LTR_Ty1_copia_Ivana_summary.png

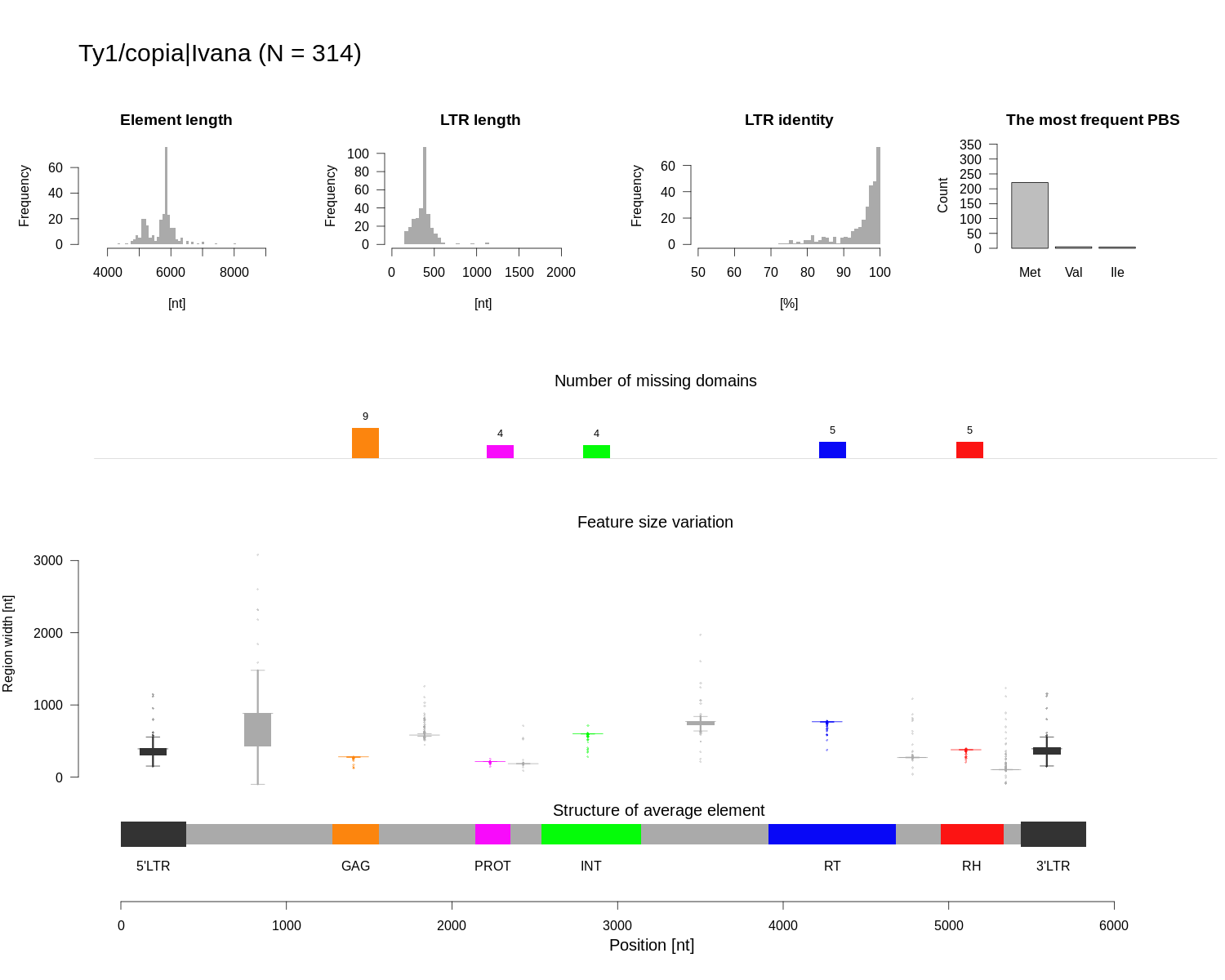

### Class_I_LTR_Ty1_copia_Ivana_summary.png

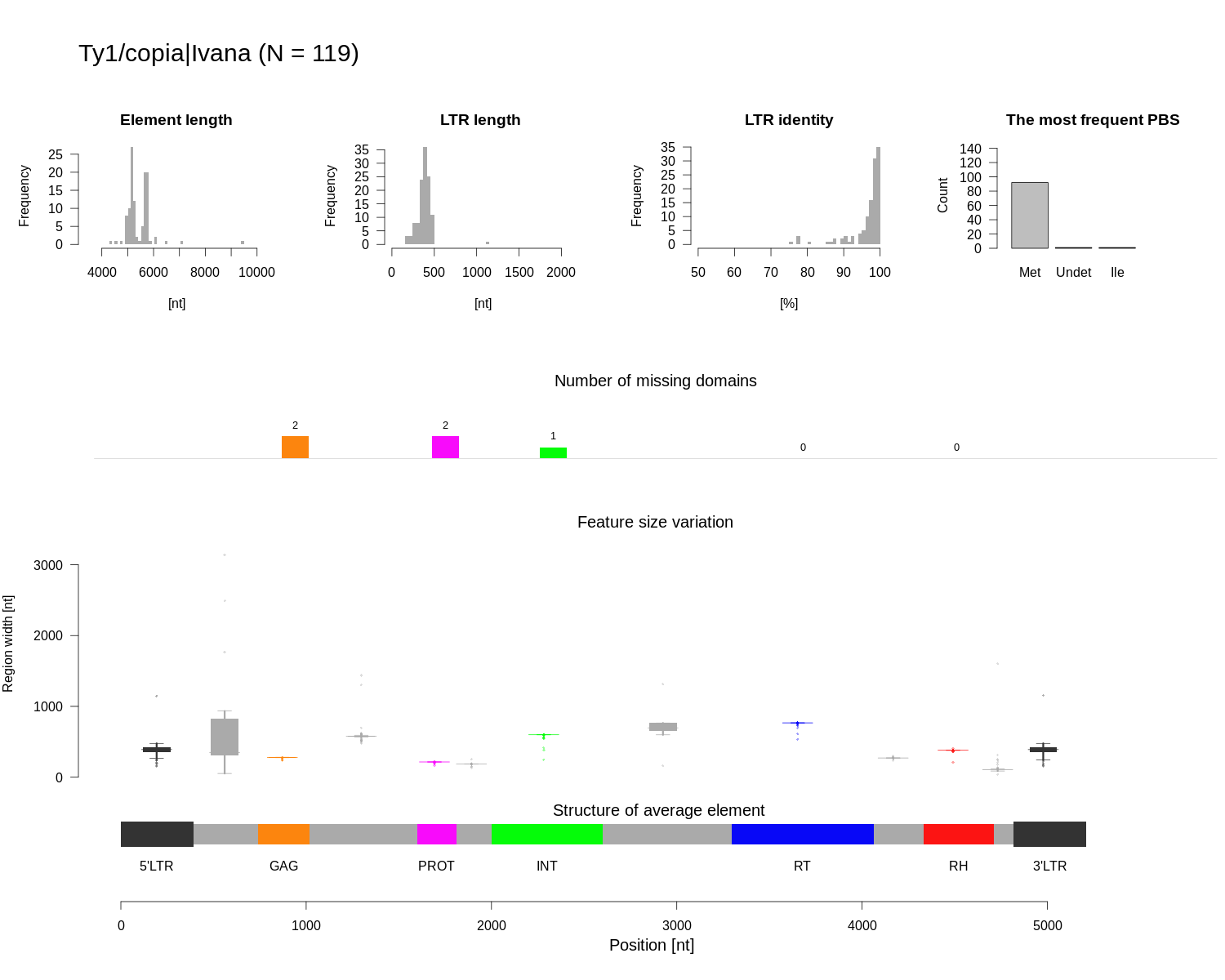

### Class_I_LTR_Ty1_copia_SIRE_structure.png

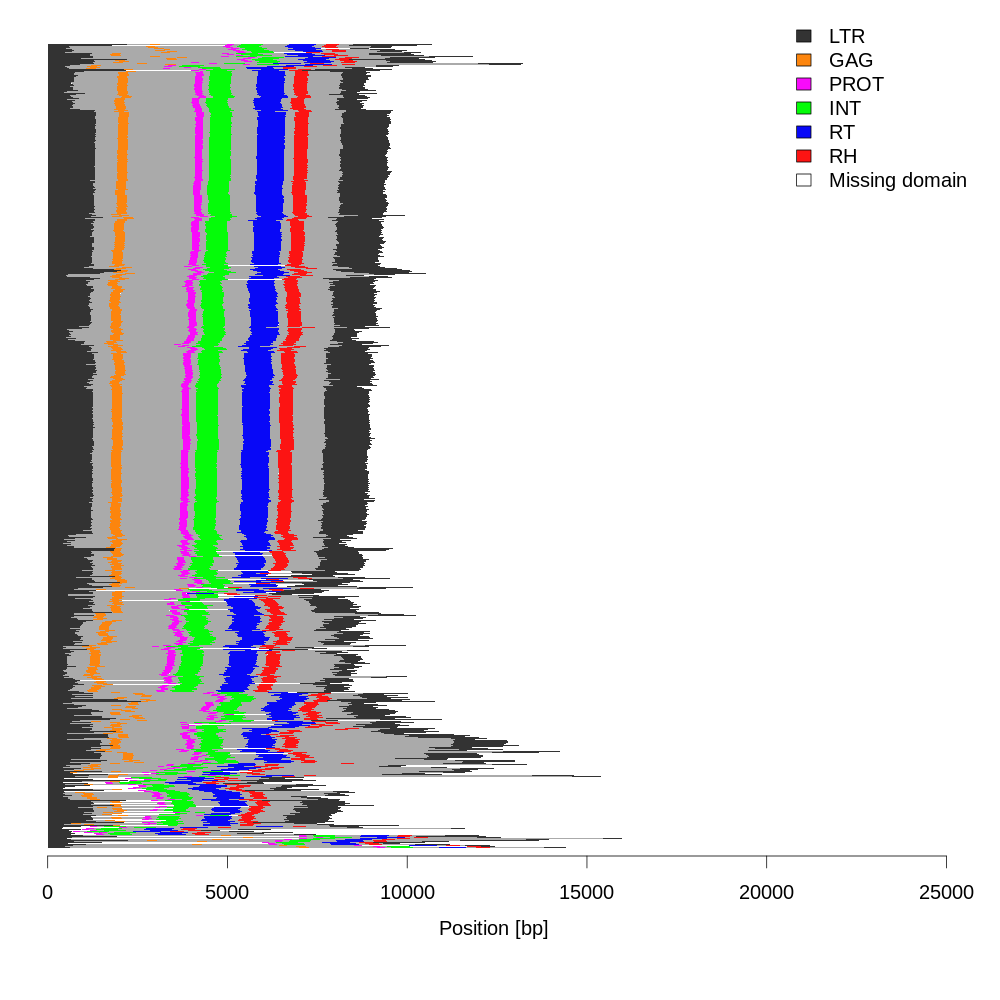

### Class_I_LTR_Ty1_copia_SIRE_structure.png

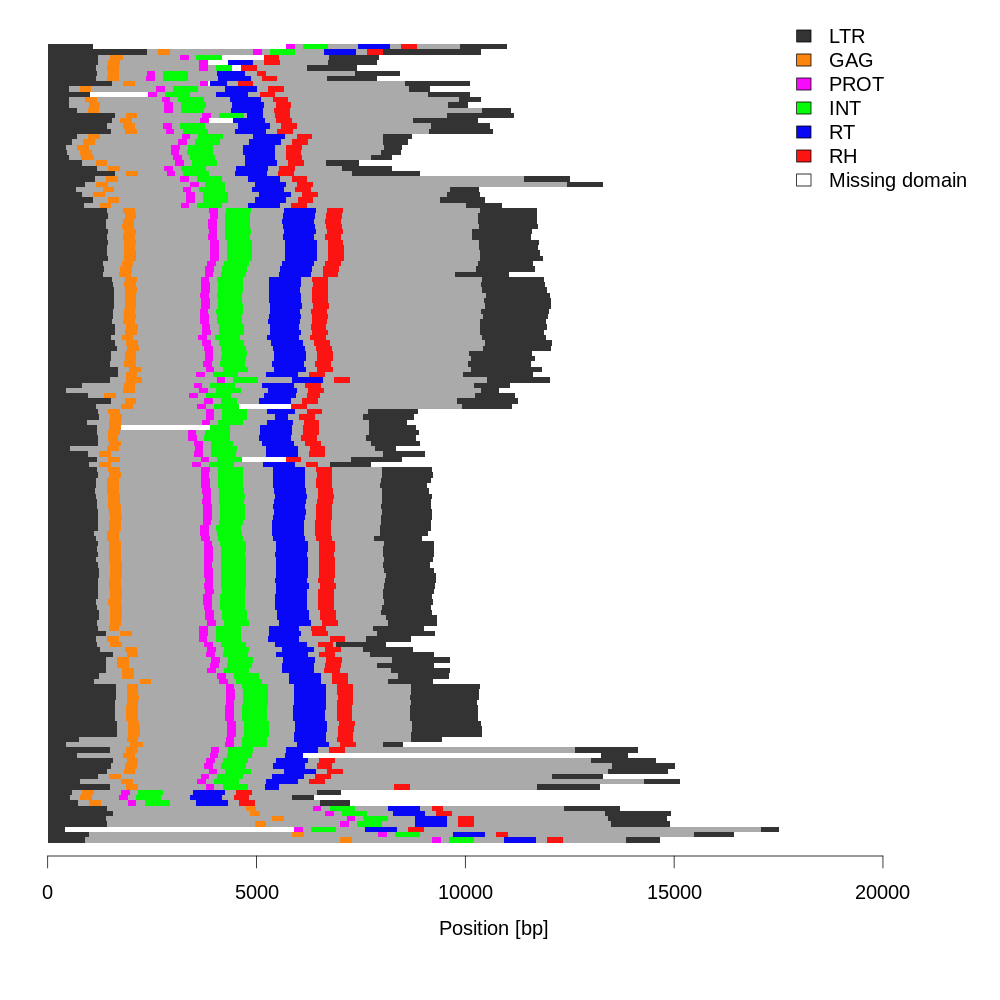

### Class_I_LTR_Ty1_copia_SIRE_summary.png

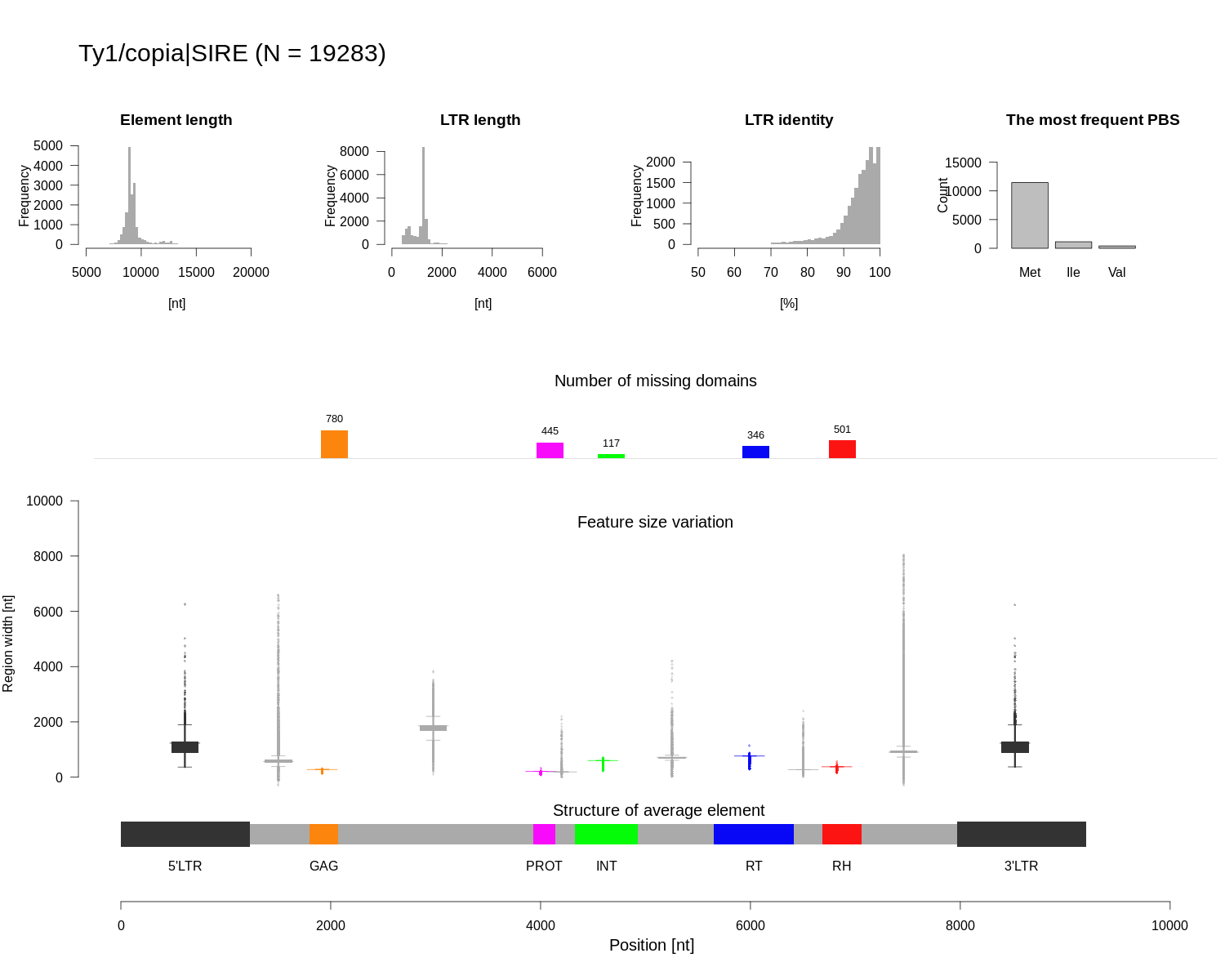

### Class_I_LTR_Ty1_copia_SIRE_summary.png

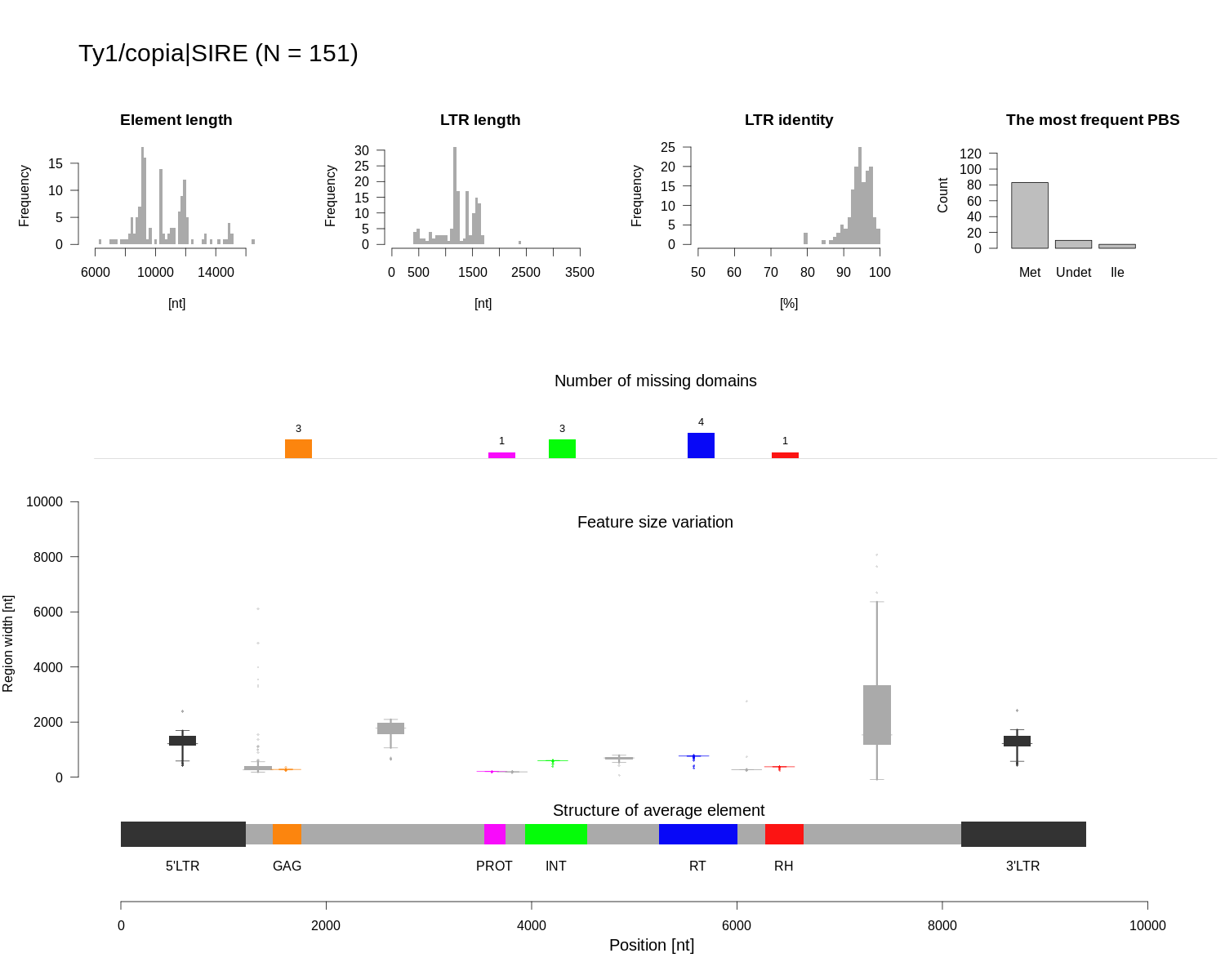

### Class_I_LTR_Ty1_copia_TAR_structure.png

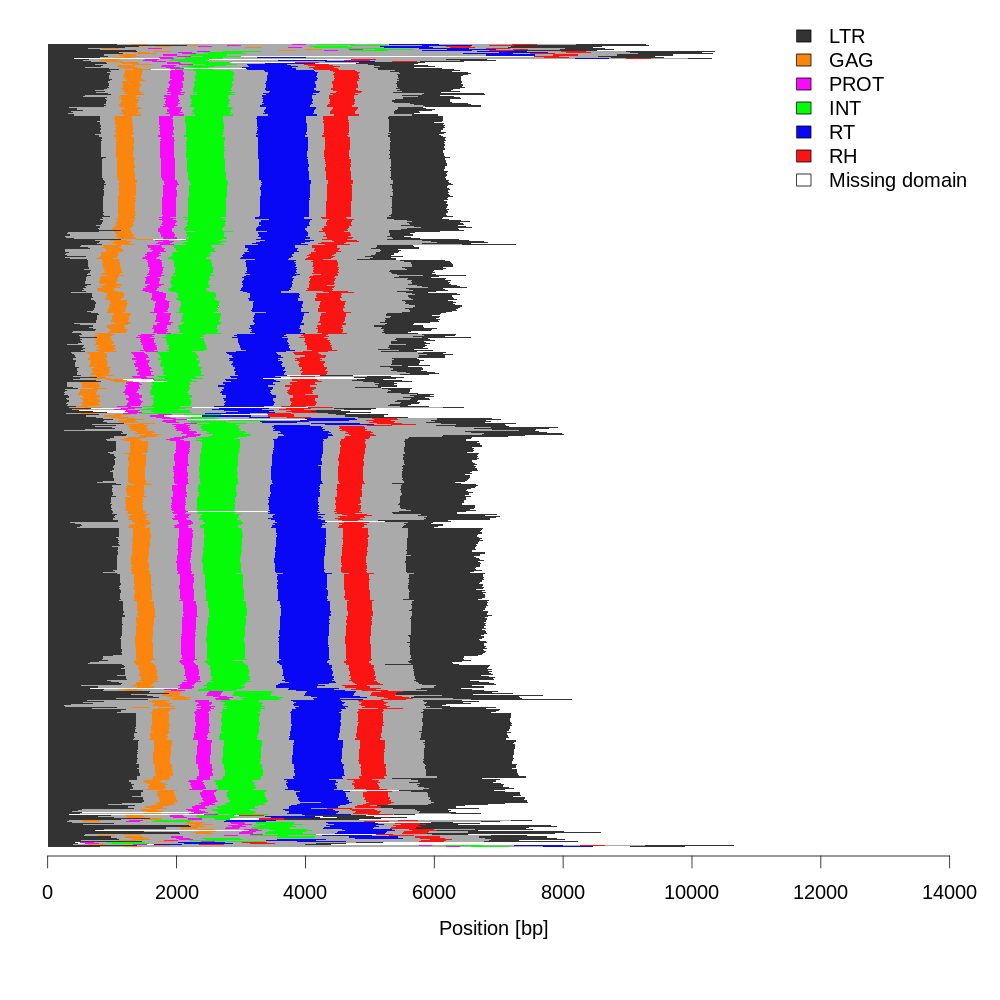

### Class_I_LTR_Ty1_copia_TAR_structure.png

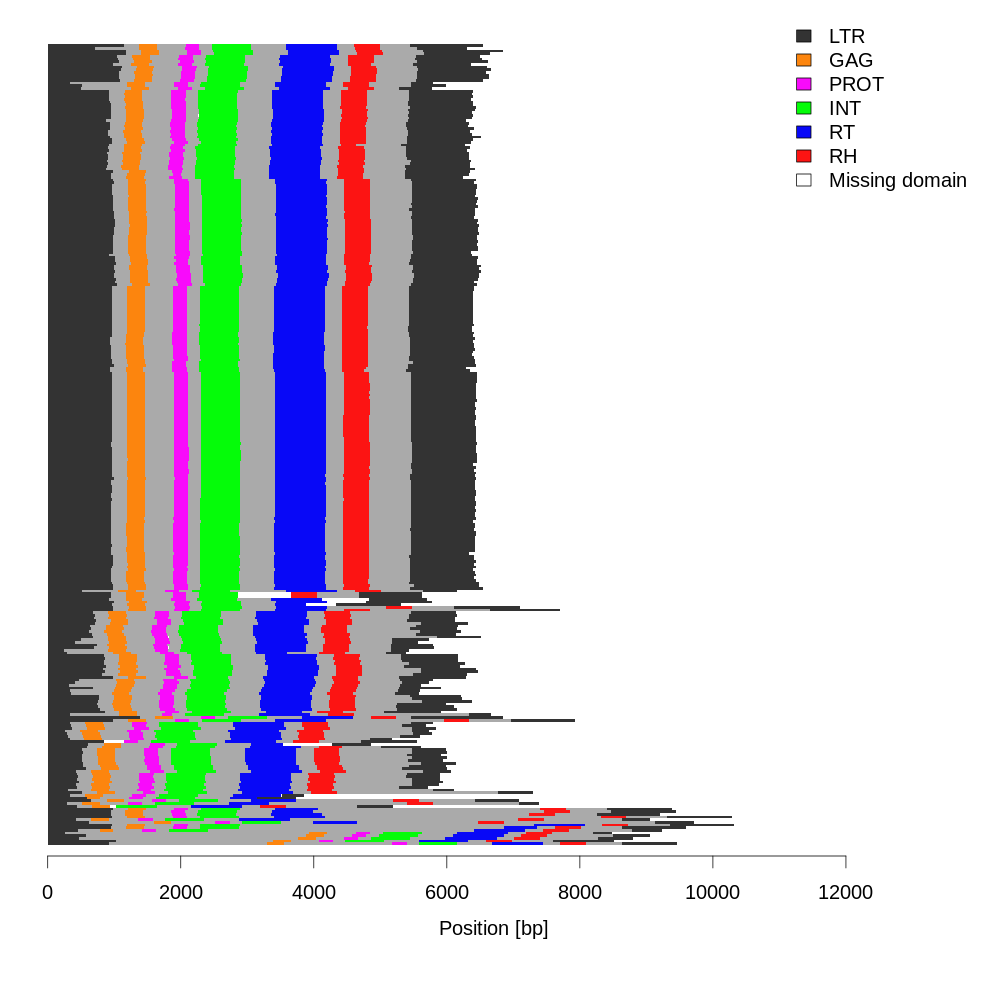

### Class_I_LTR_Ty1_copia_TAR_summary.png

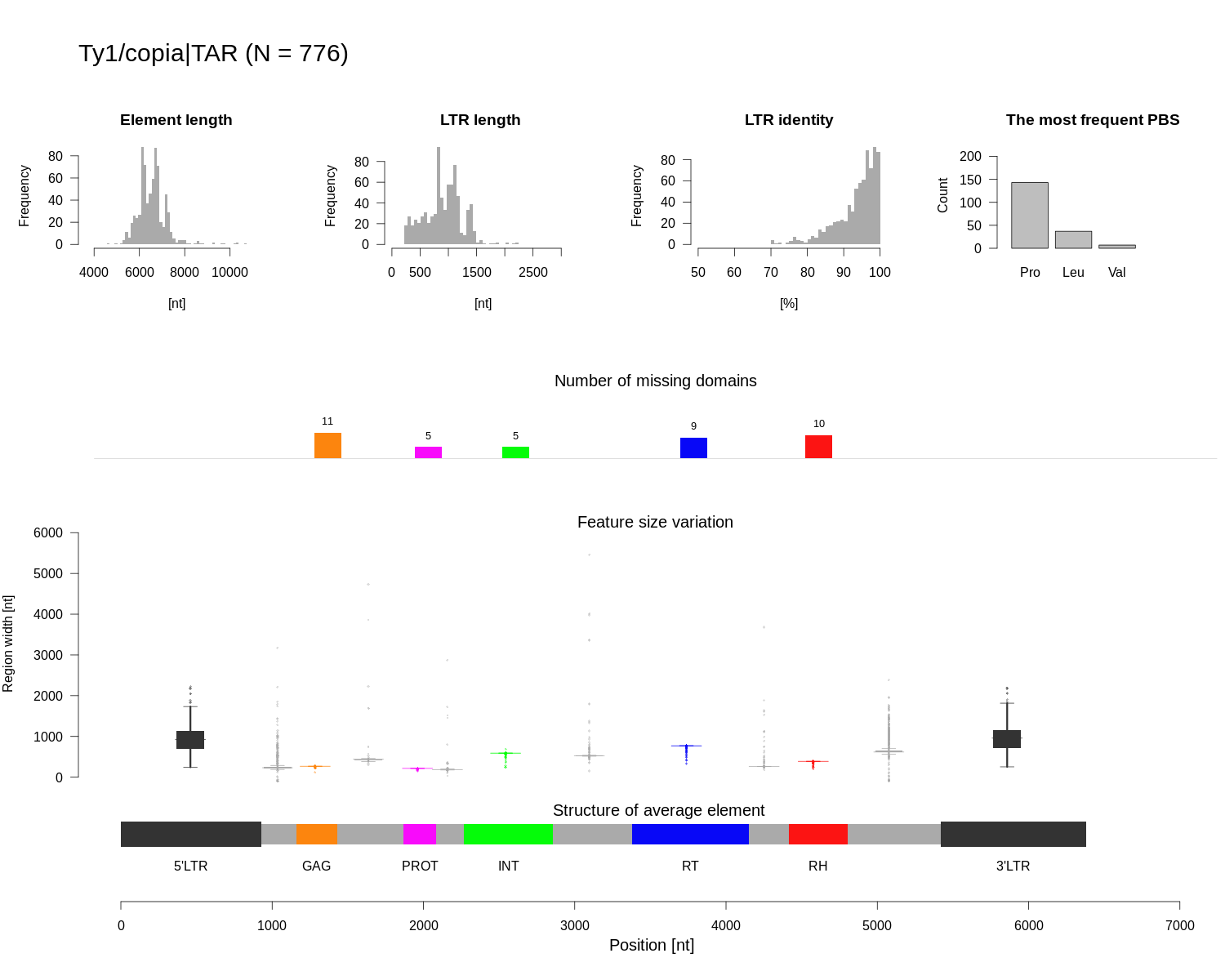

### Class_I_LTR_Ty1_copia_TAR_summary.png

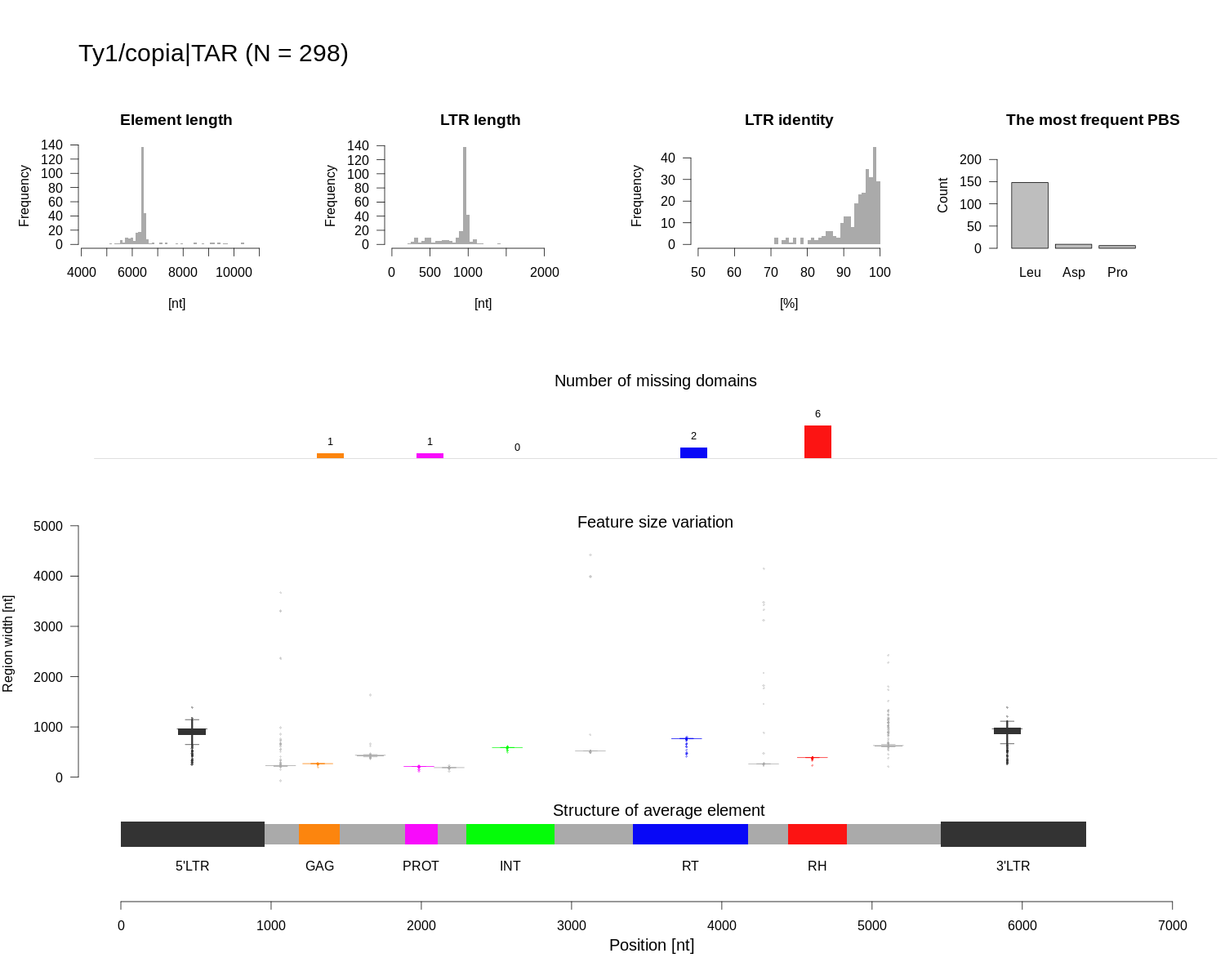

### Class_I_LTR_Ty1_copia_Tork_structure.png

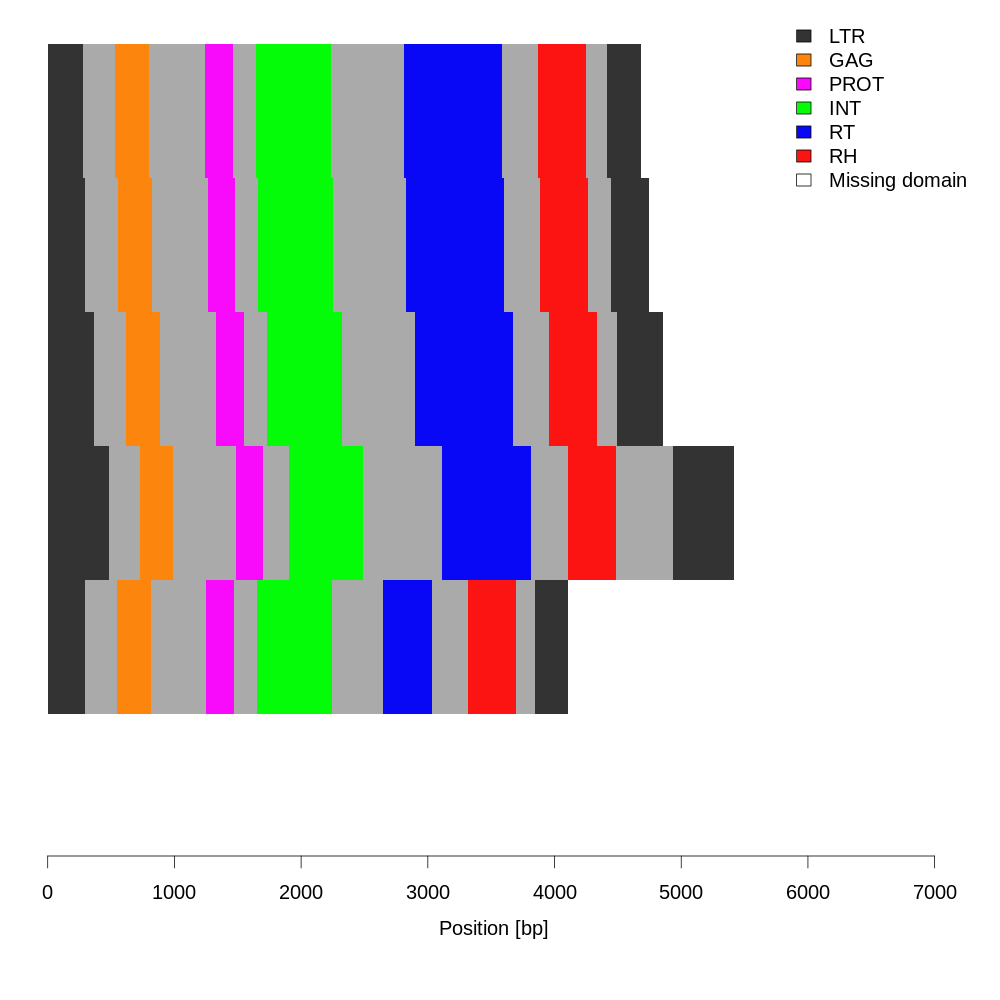

### Class_I_LTR_Ty1_copia_Tork_structure.png

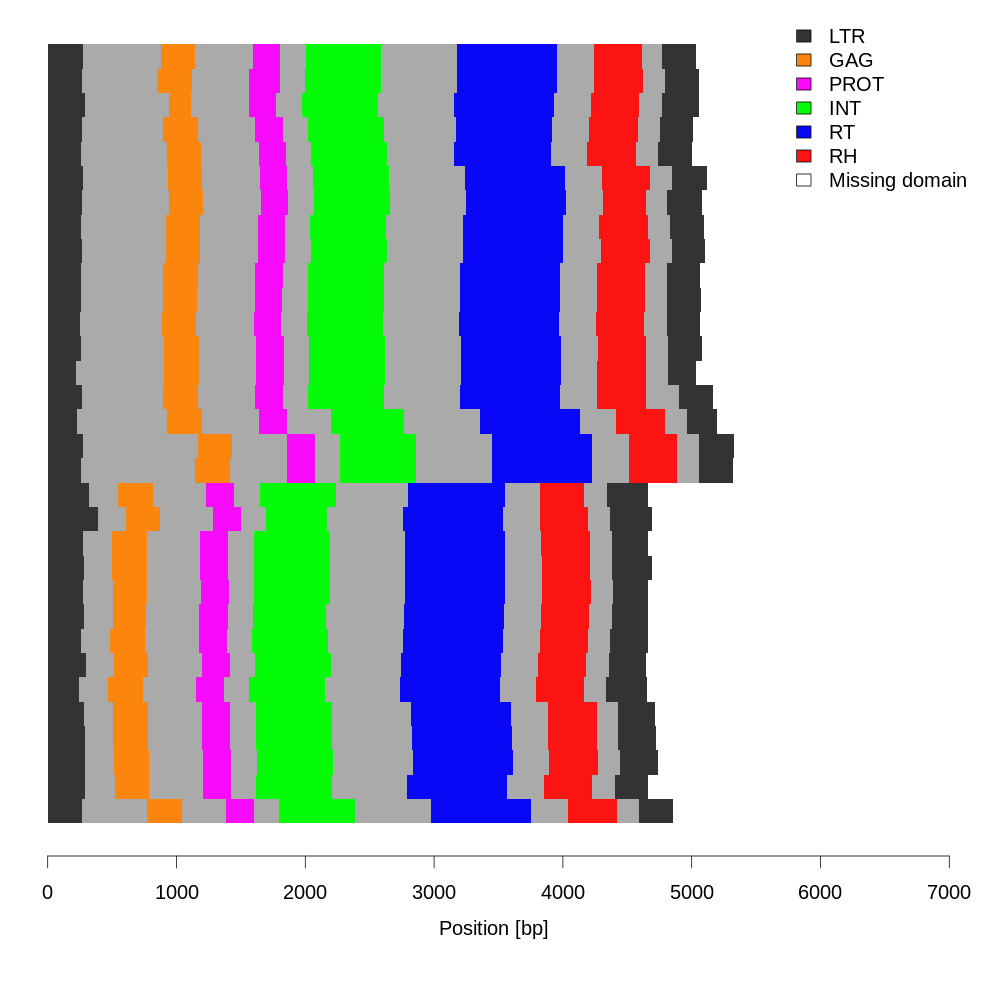

### Class_I_LTR_Ty1_copia_Tork_summary.png

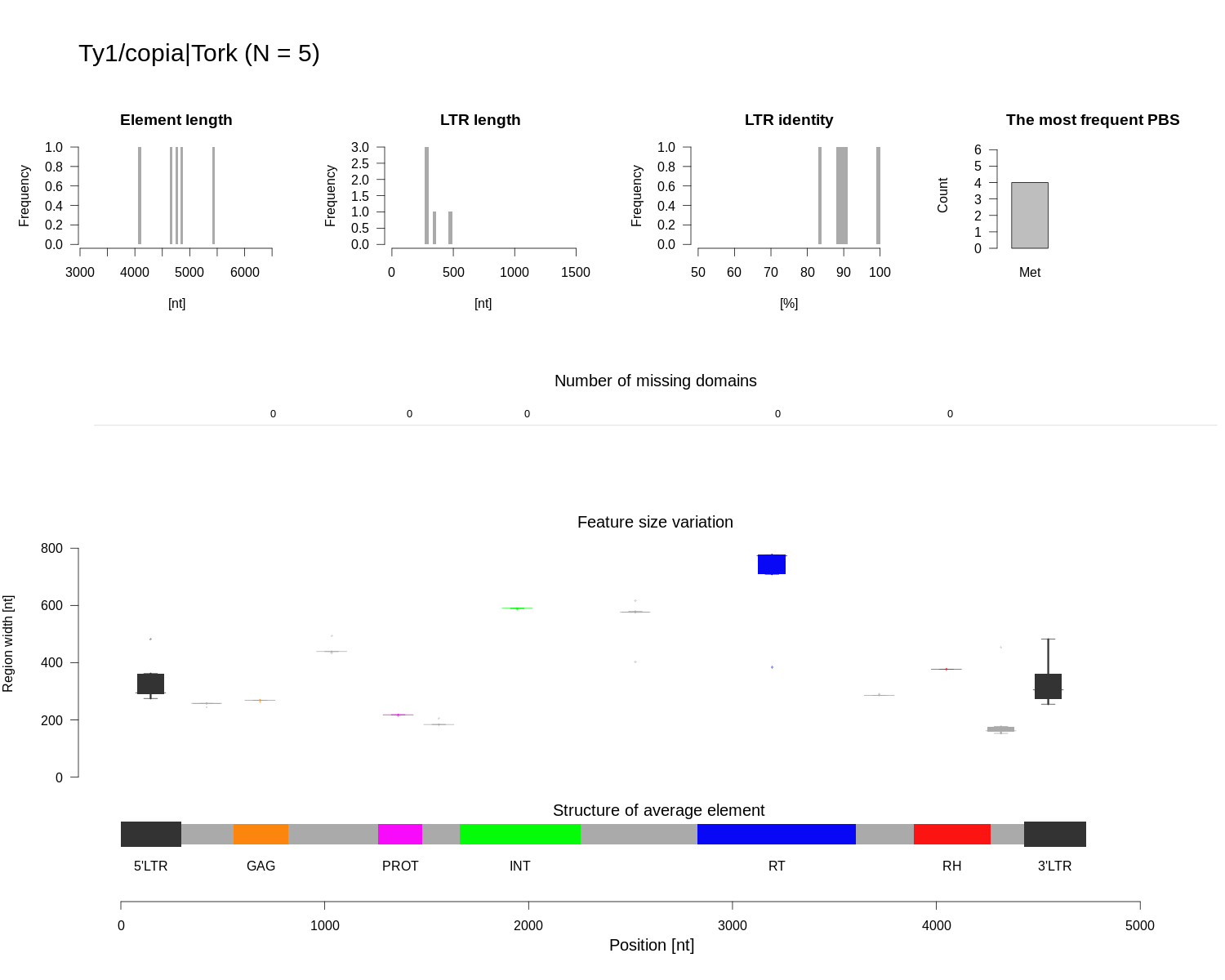

### Class_I_LTR_Ty1_copia_Tork_summary.png

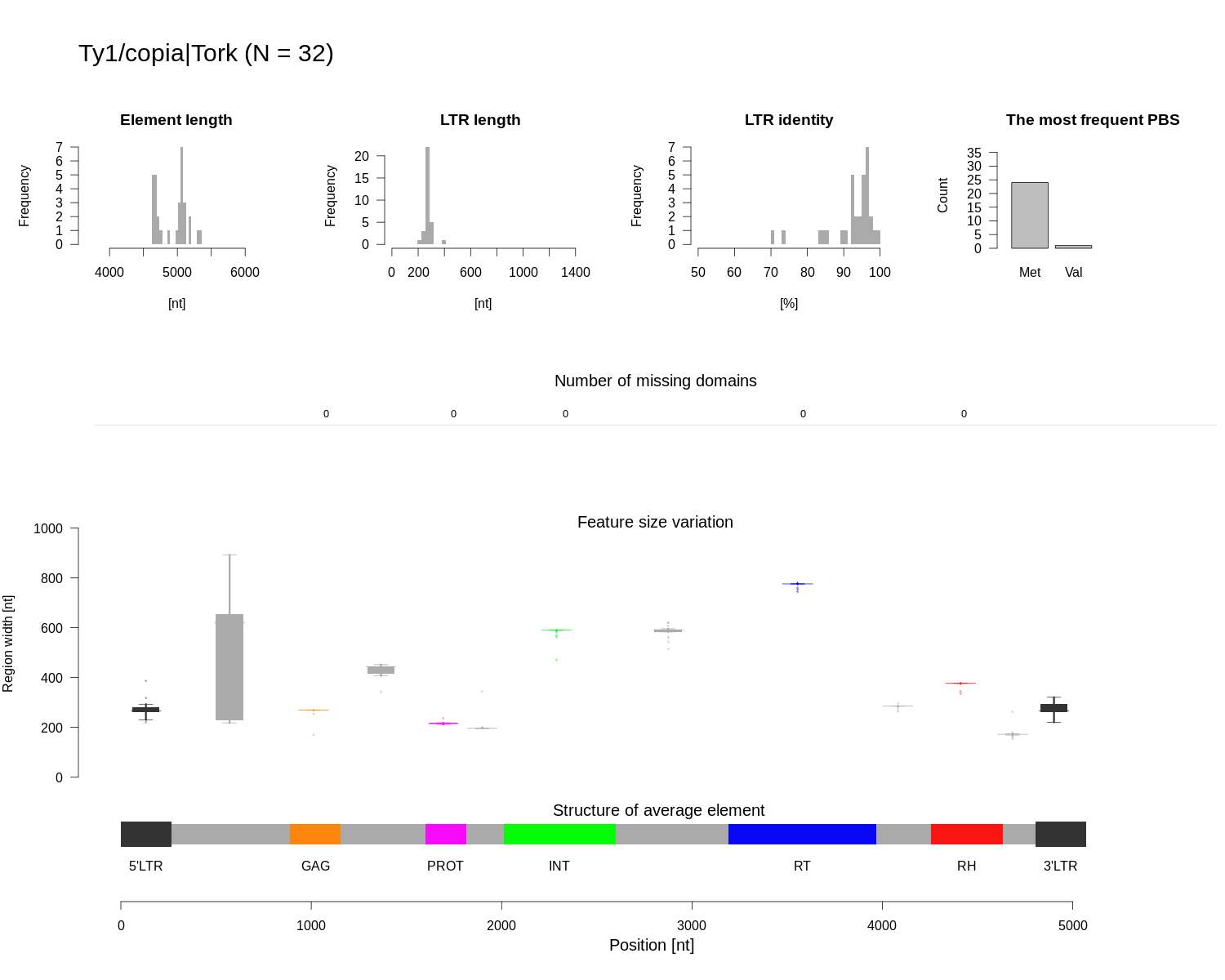

### Class_I_LTR_Ty3_gypsy_chromovirus_CRM_structure.png

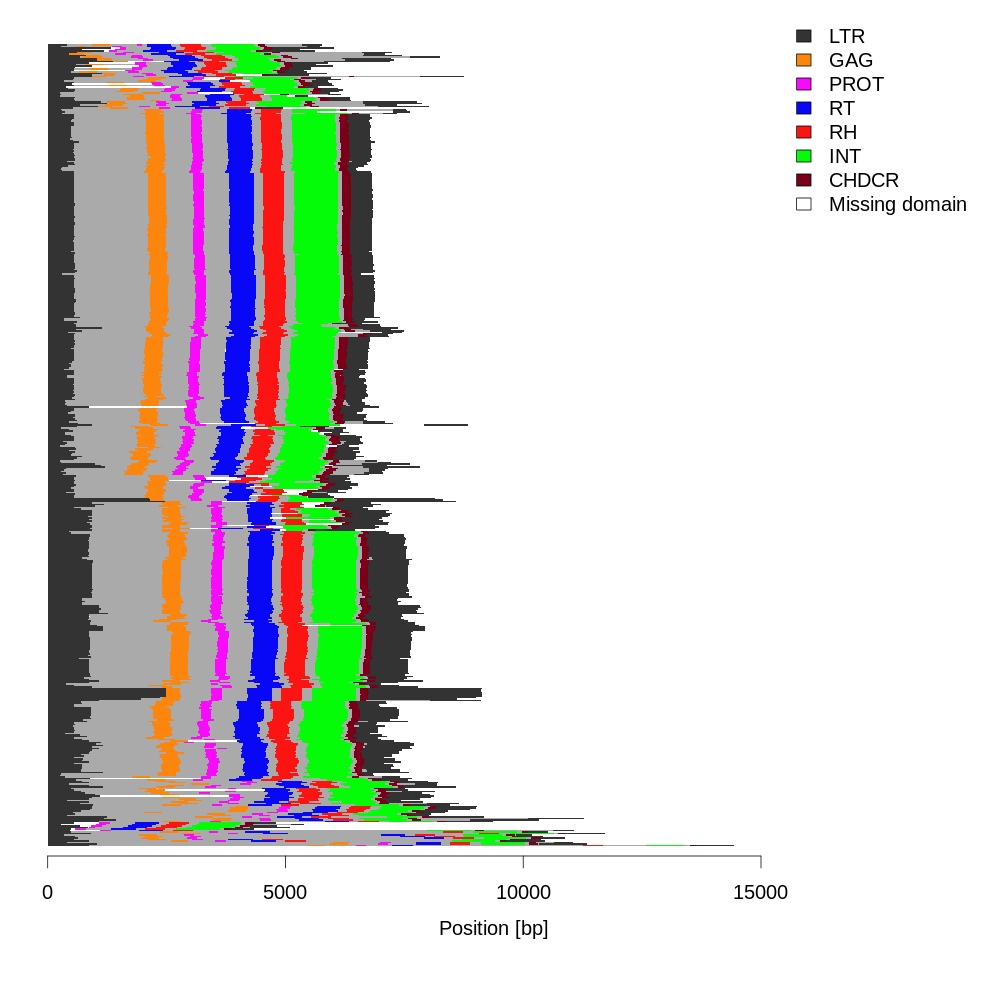

### Class_I_LTR_Ty3_gypsy_chromovirus_CRM_structure.png

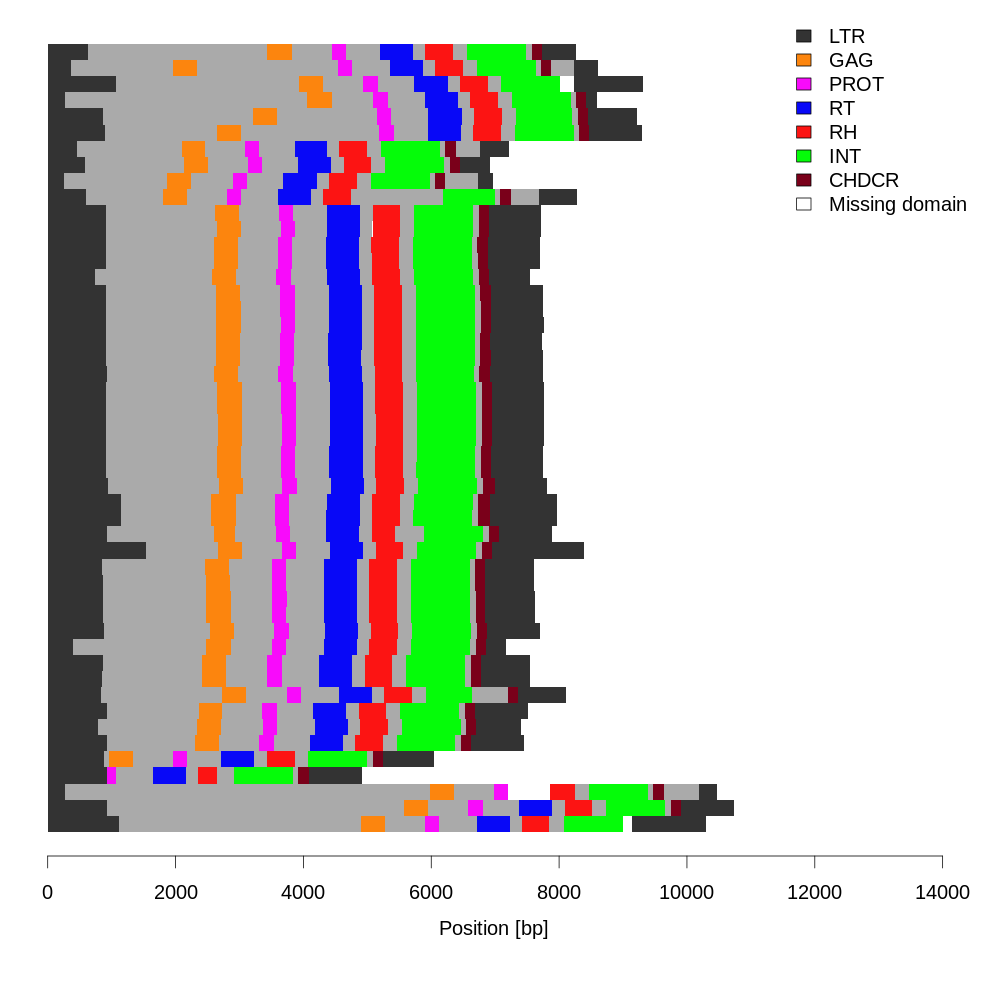

### Class_I_LTR_Ty3_gypsy_chromovirus_CRM_summary.png

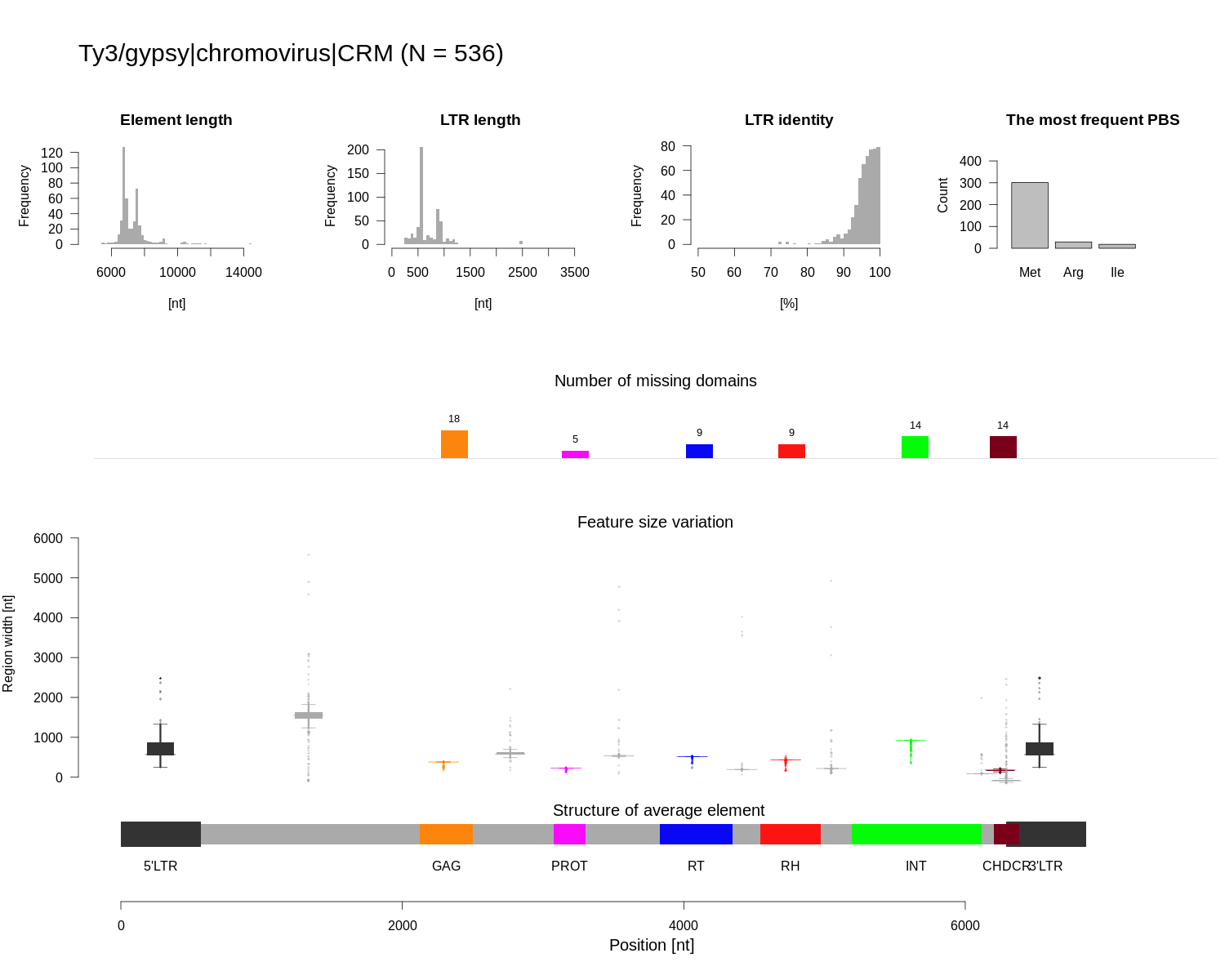

### Class_I_LTR_Ty3_gypsy_chromovirus_CRM_summary.png

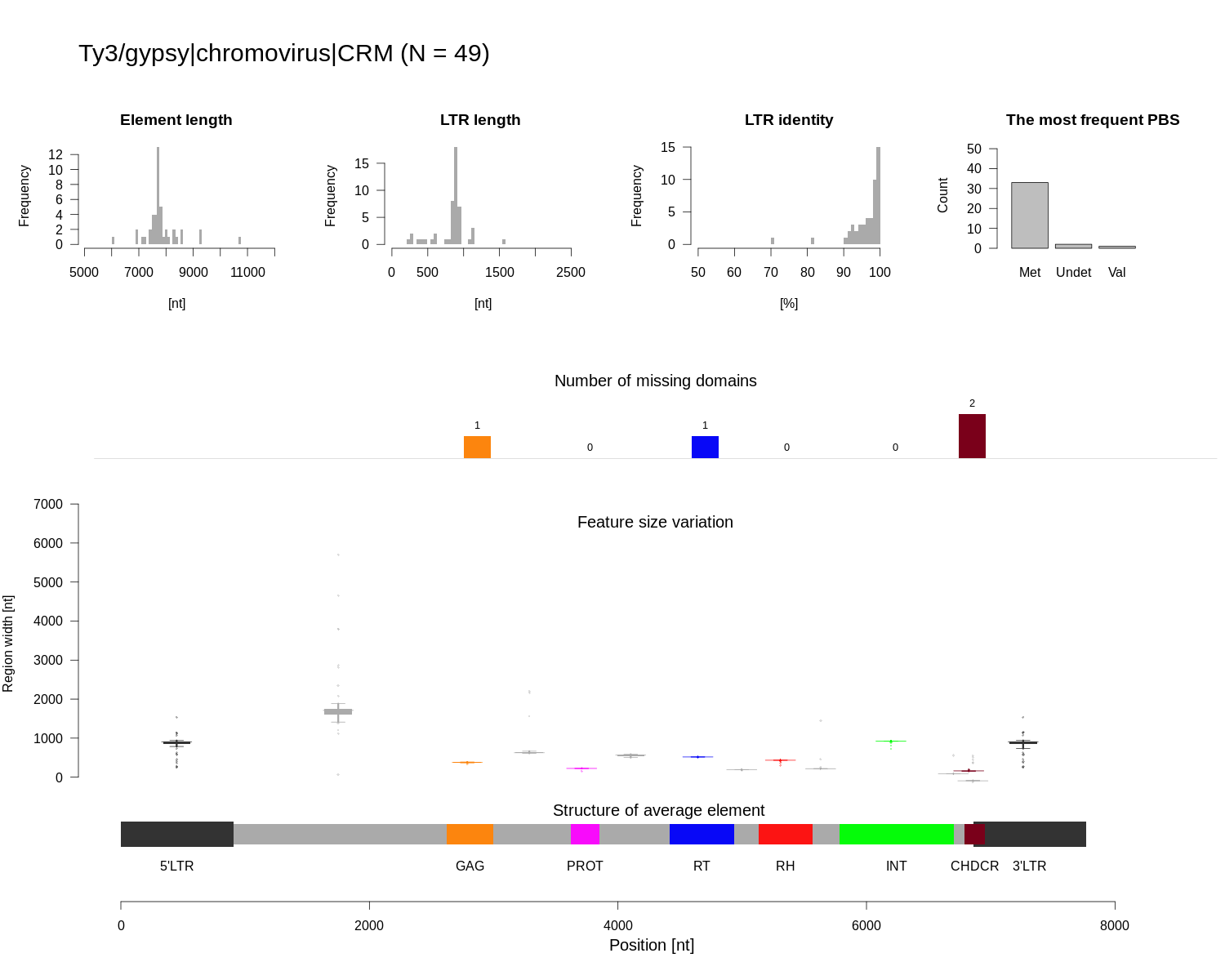

### Class_I_LTR_Ty3_gypsy_chromovirus_Reina_structure.png

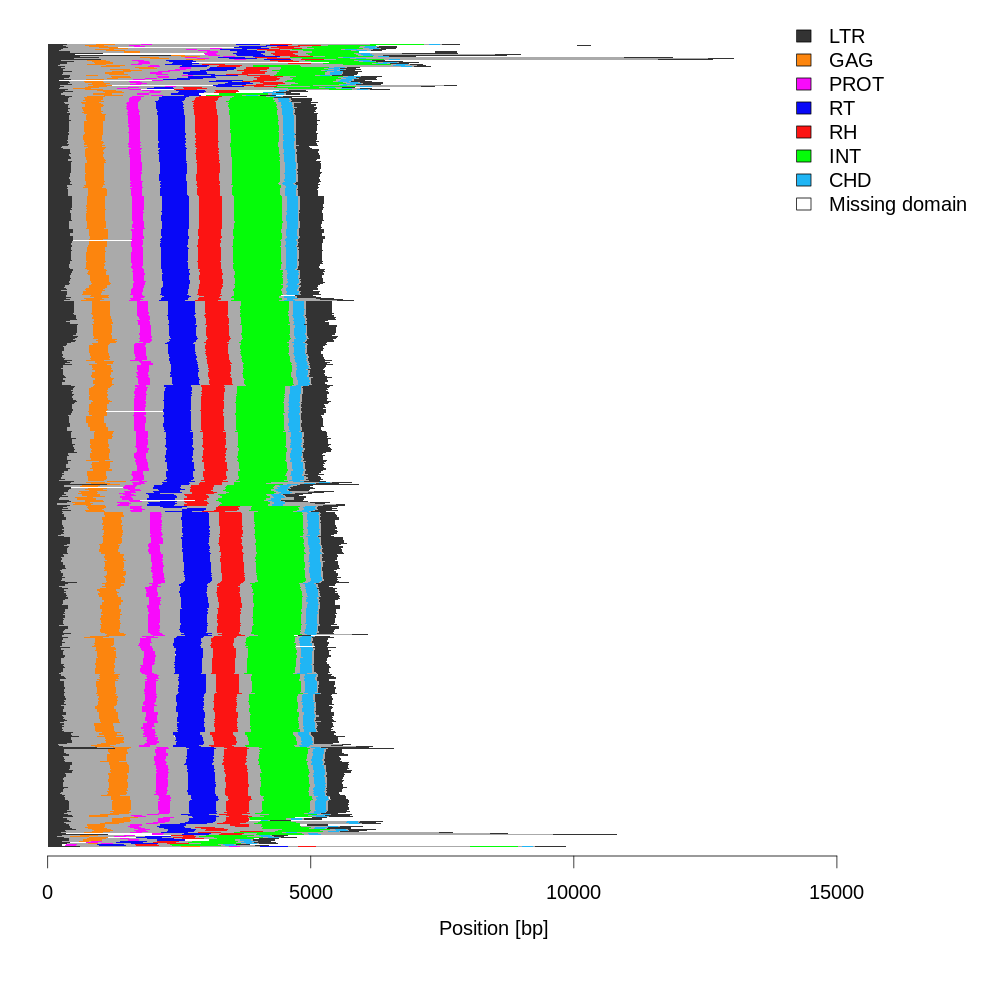

### Class_I_LTR_Ty3_gypsy_chromovirus_Reina_structure.png

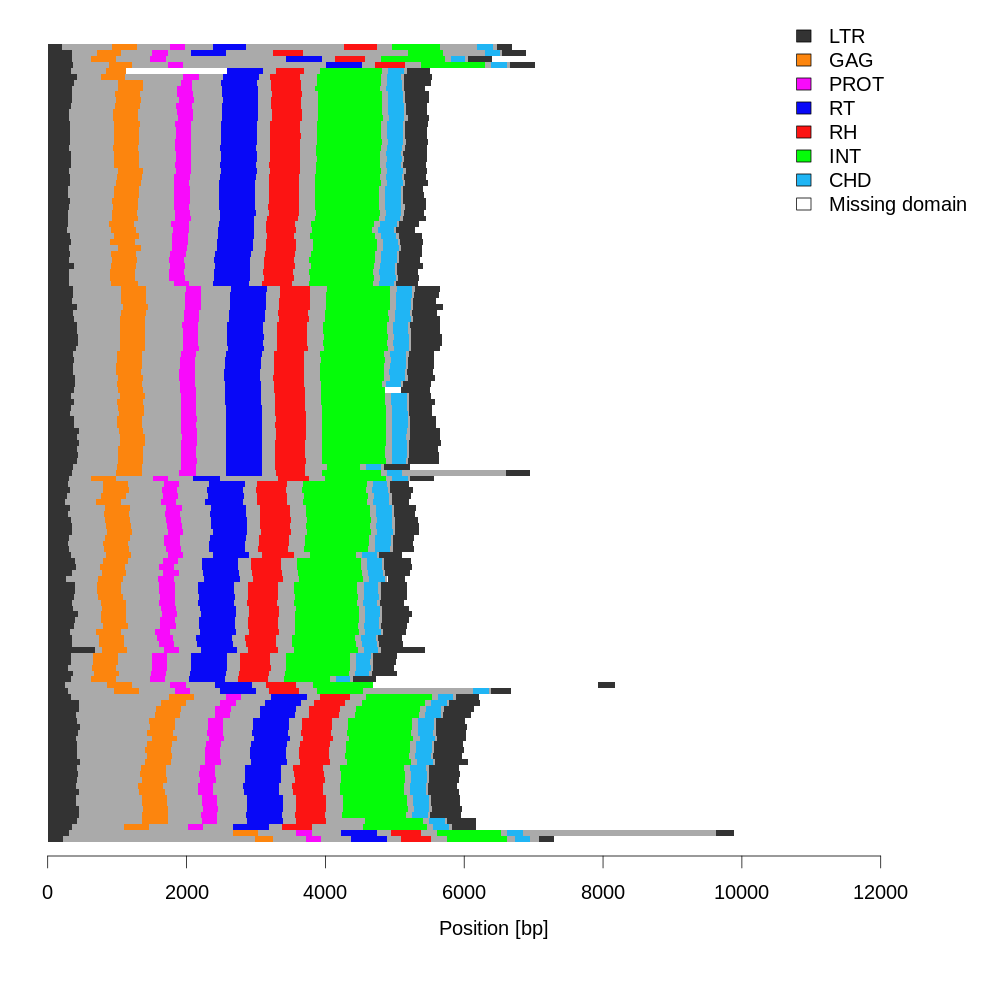

### Class_I_LTR_Ty3_gypsy_chromovirus_Reina_summary.png

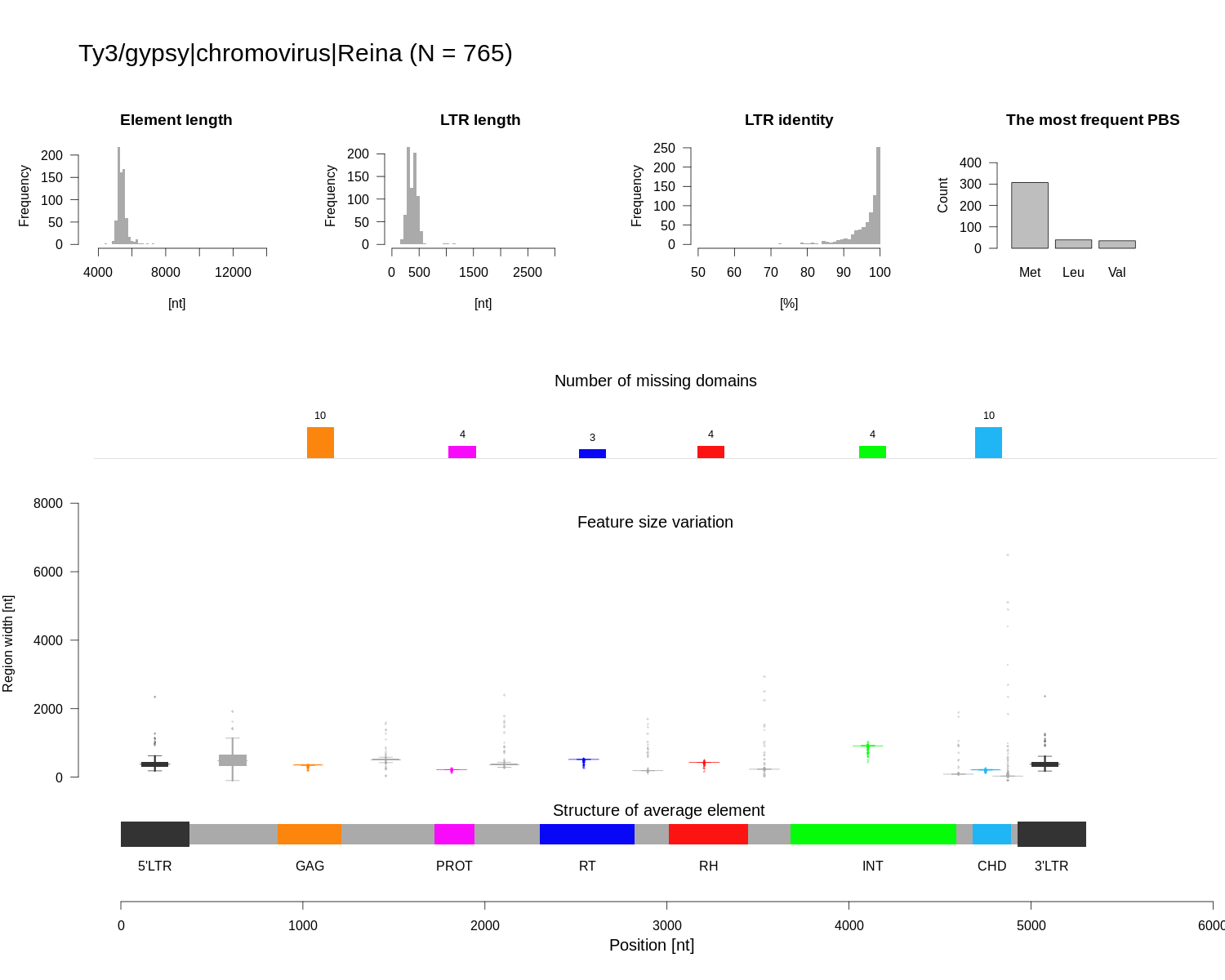

### Class_I_LTR_Ty3_gypsy_chromovirus_Reina_summary.png

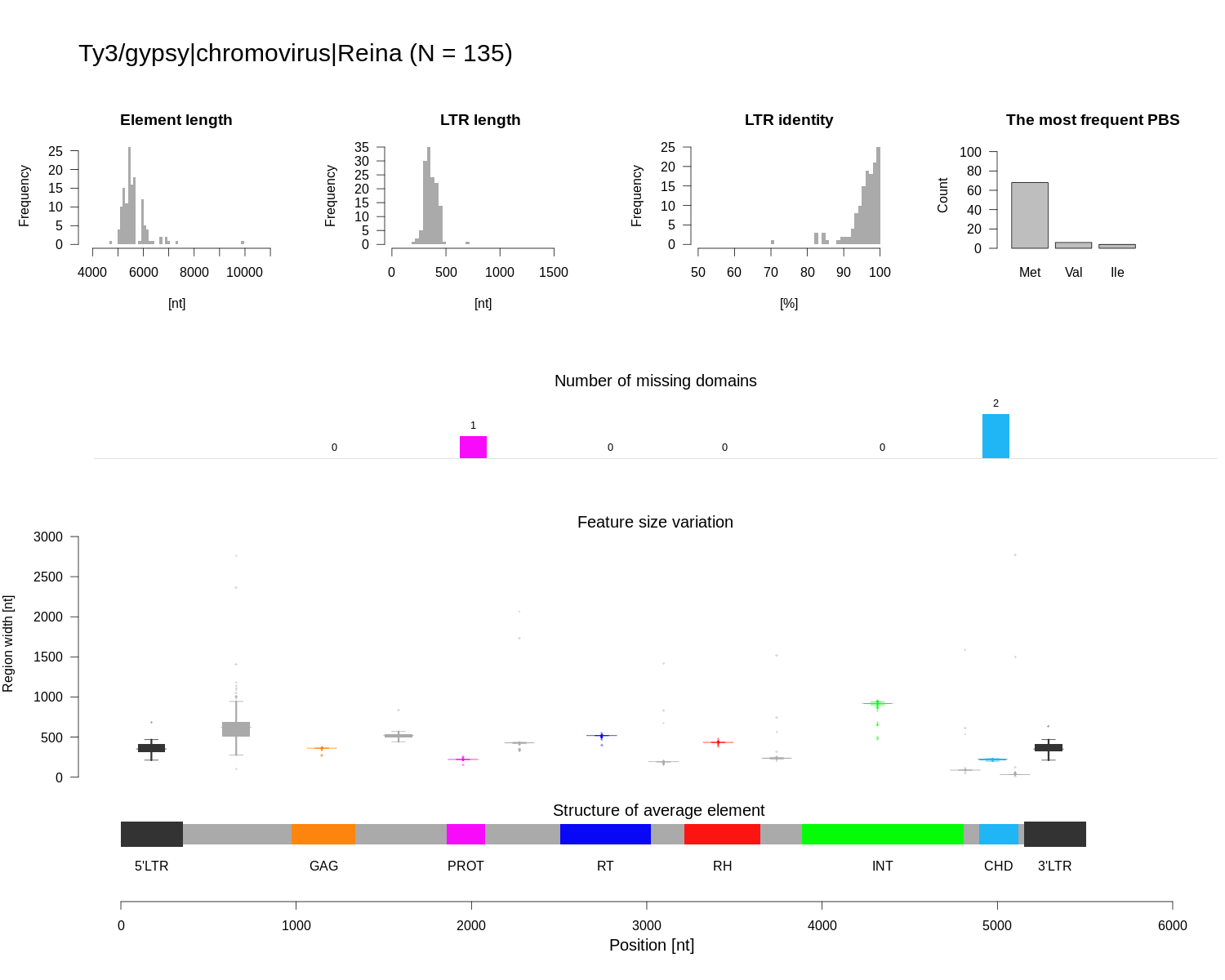

### Class_I_LTR_Ty3_gypsy_chromovirus_Tekay_structure.png

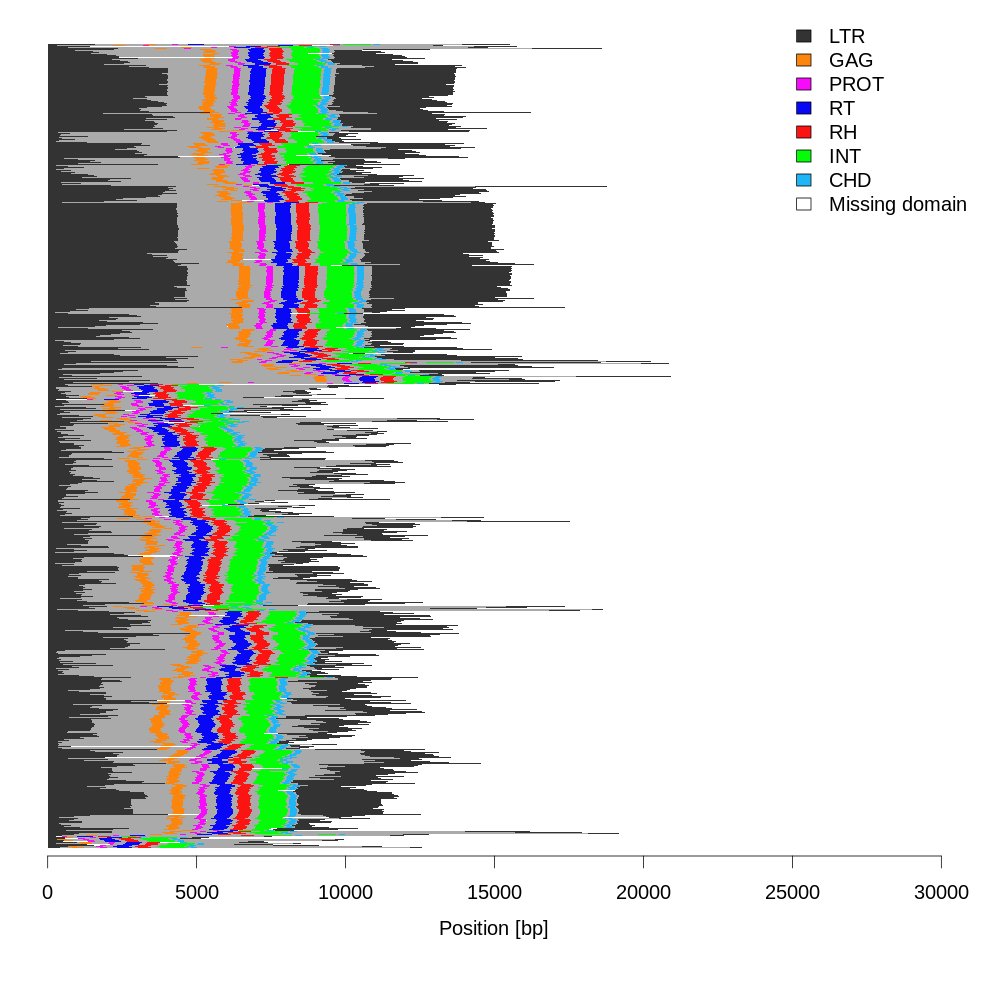

### Class_I_LTR_Ty3_gypsy_chromovirus_Tekay_structure.png

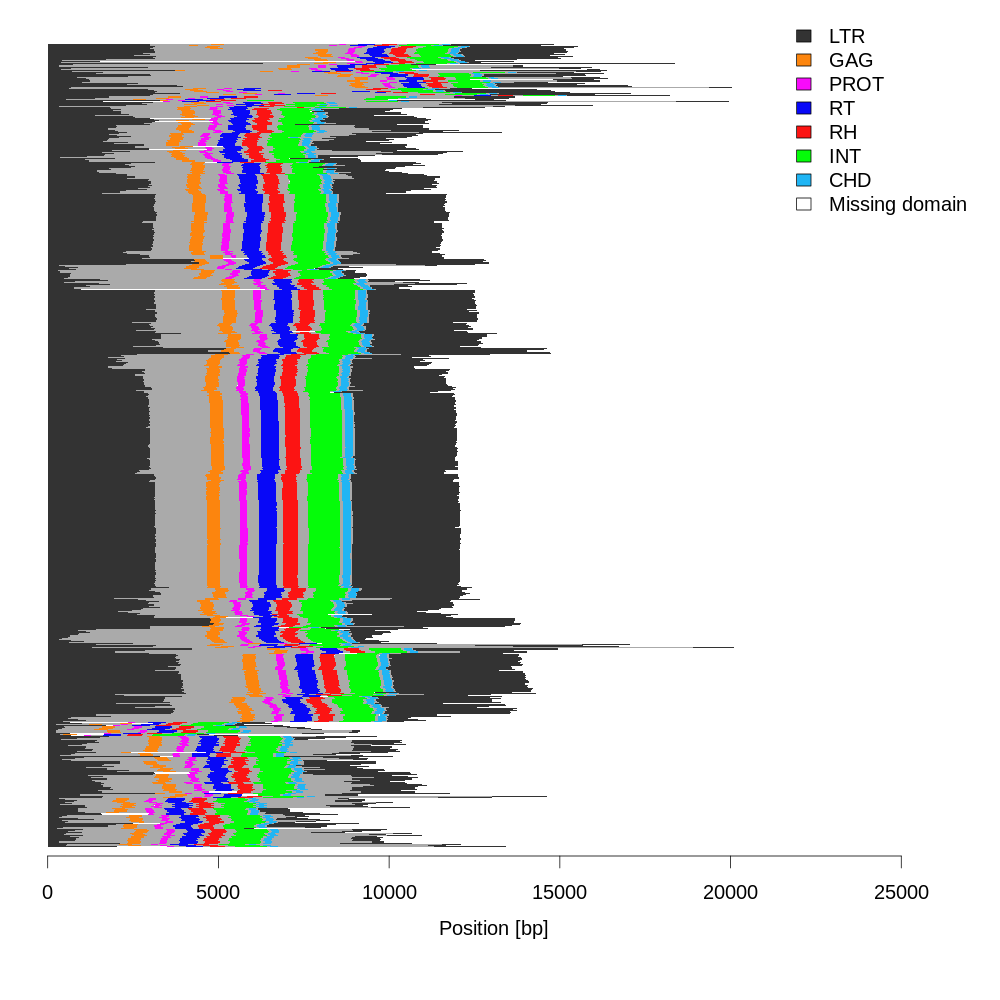

### Class_I_LTR_Ty3_gypsy_chromovirus_Tekay_summary.png

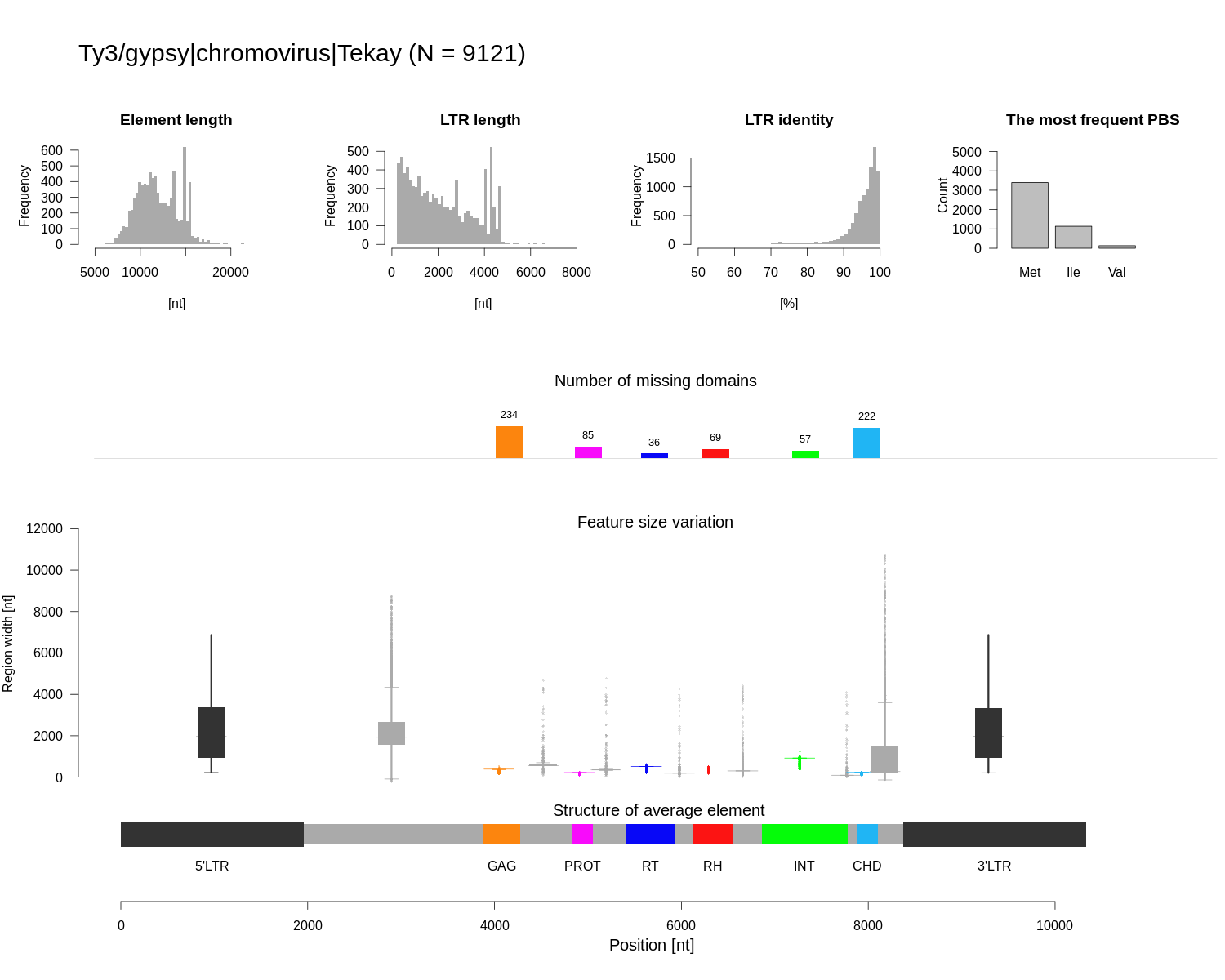

### Class_I_LTR_Ty3_gypsy_chromovirus_Tekay_summary.png

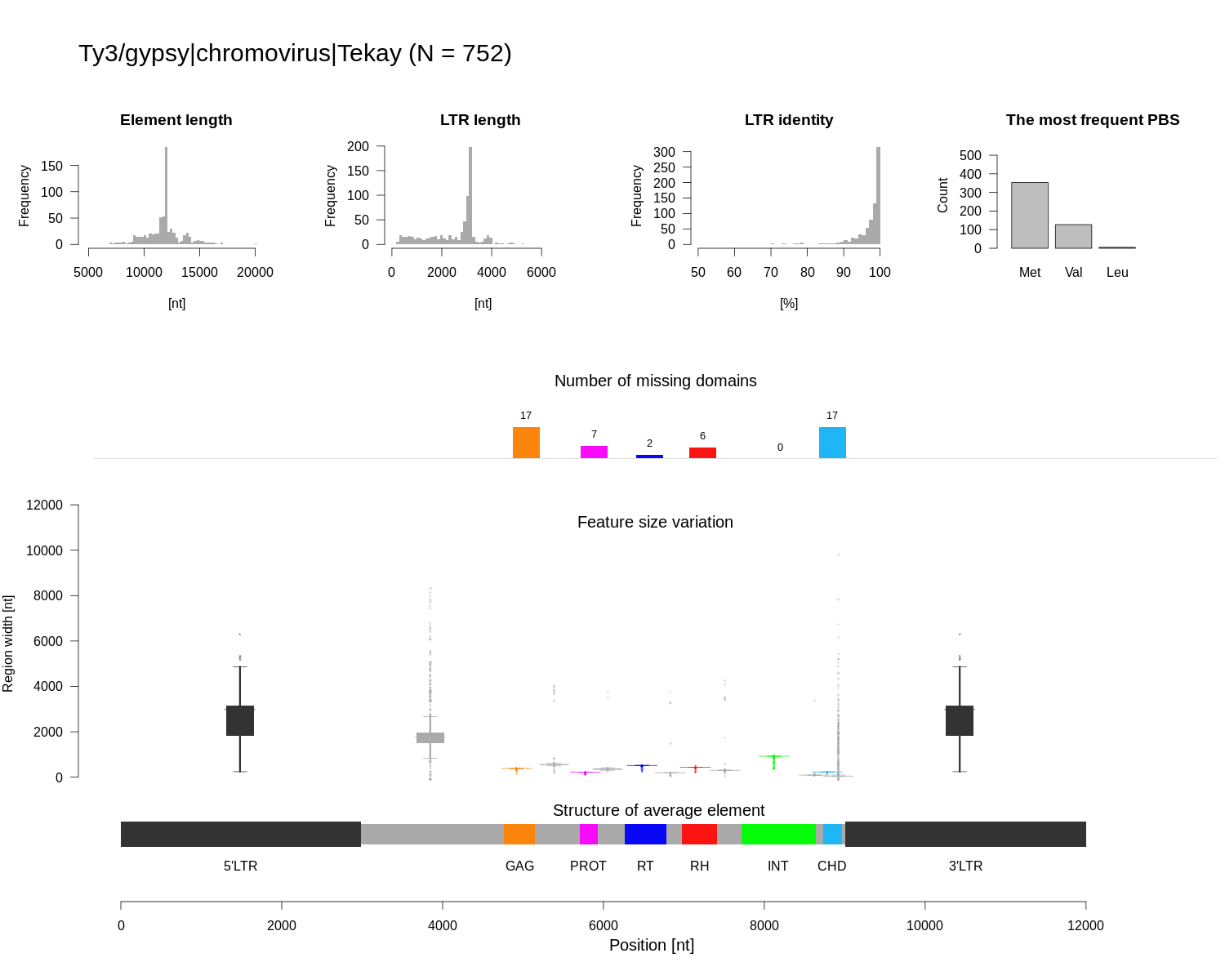

### Class_I_LTR_Ty3_gypsy_non_chromovirus_OTA_Athila_structure.png

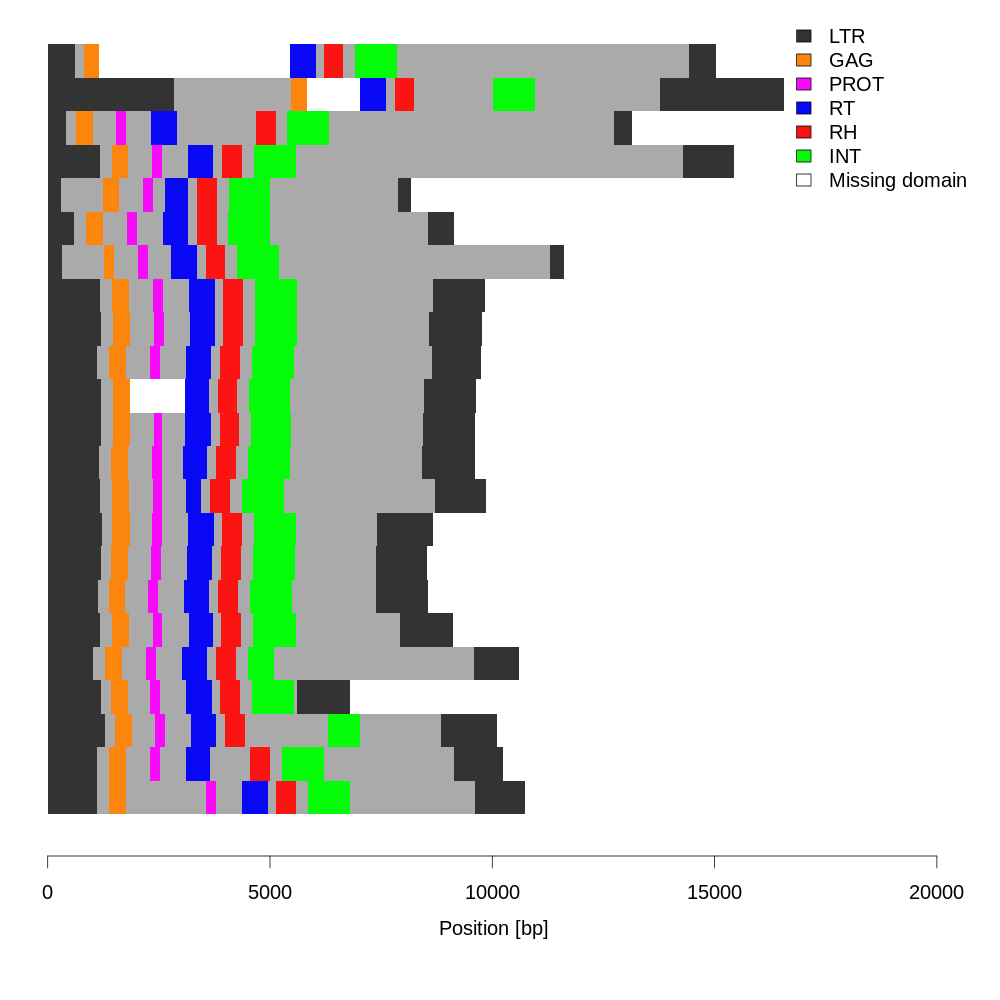

### Class_I_LTR_Ty3_gypsy_non_chromovirus_OTA_Athila_summary.png

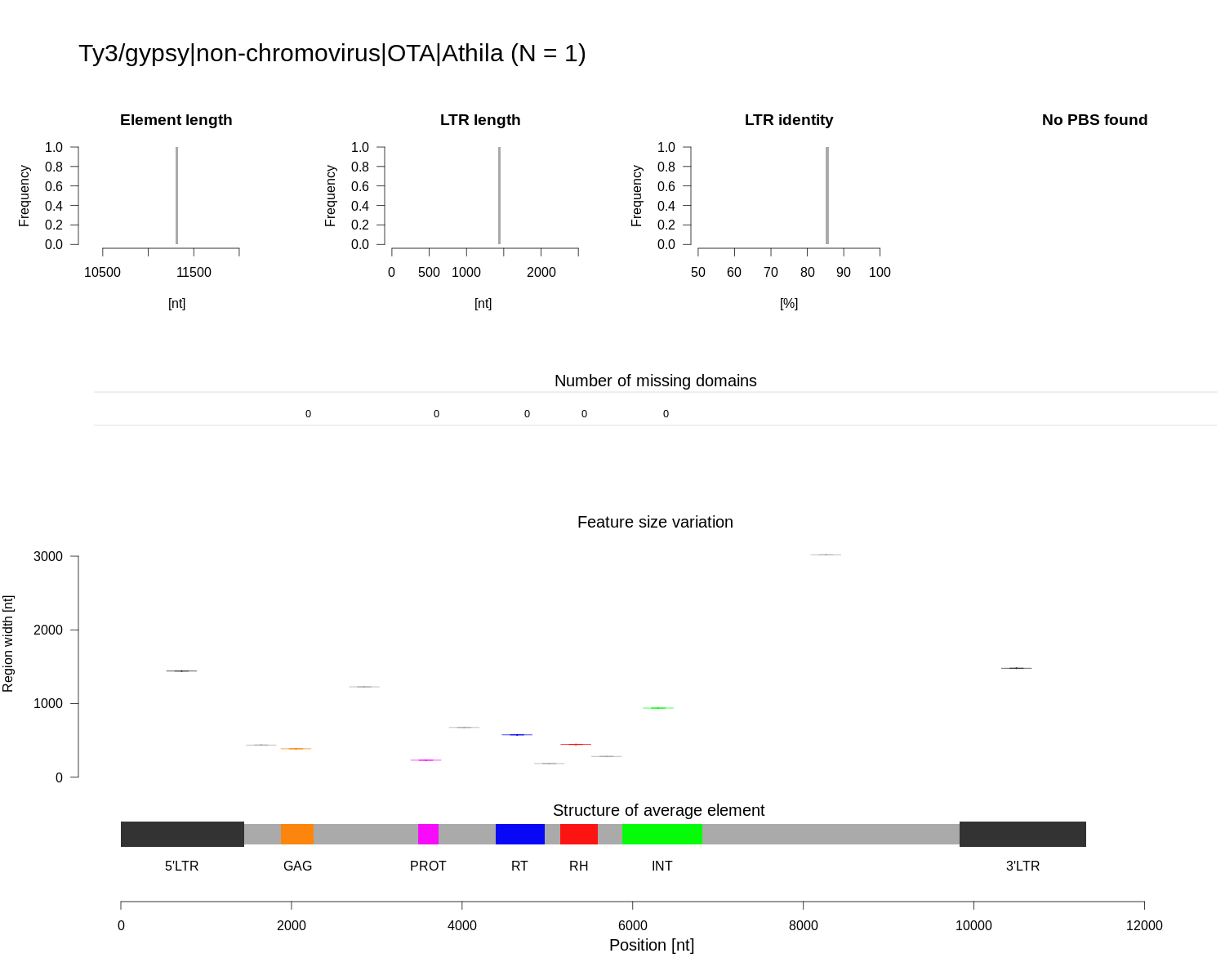

### Class_I_LTR_Ty3_gypsy_non_chromovirus_OTA_Athila_summary.png

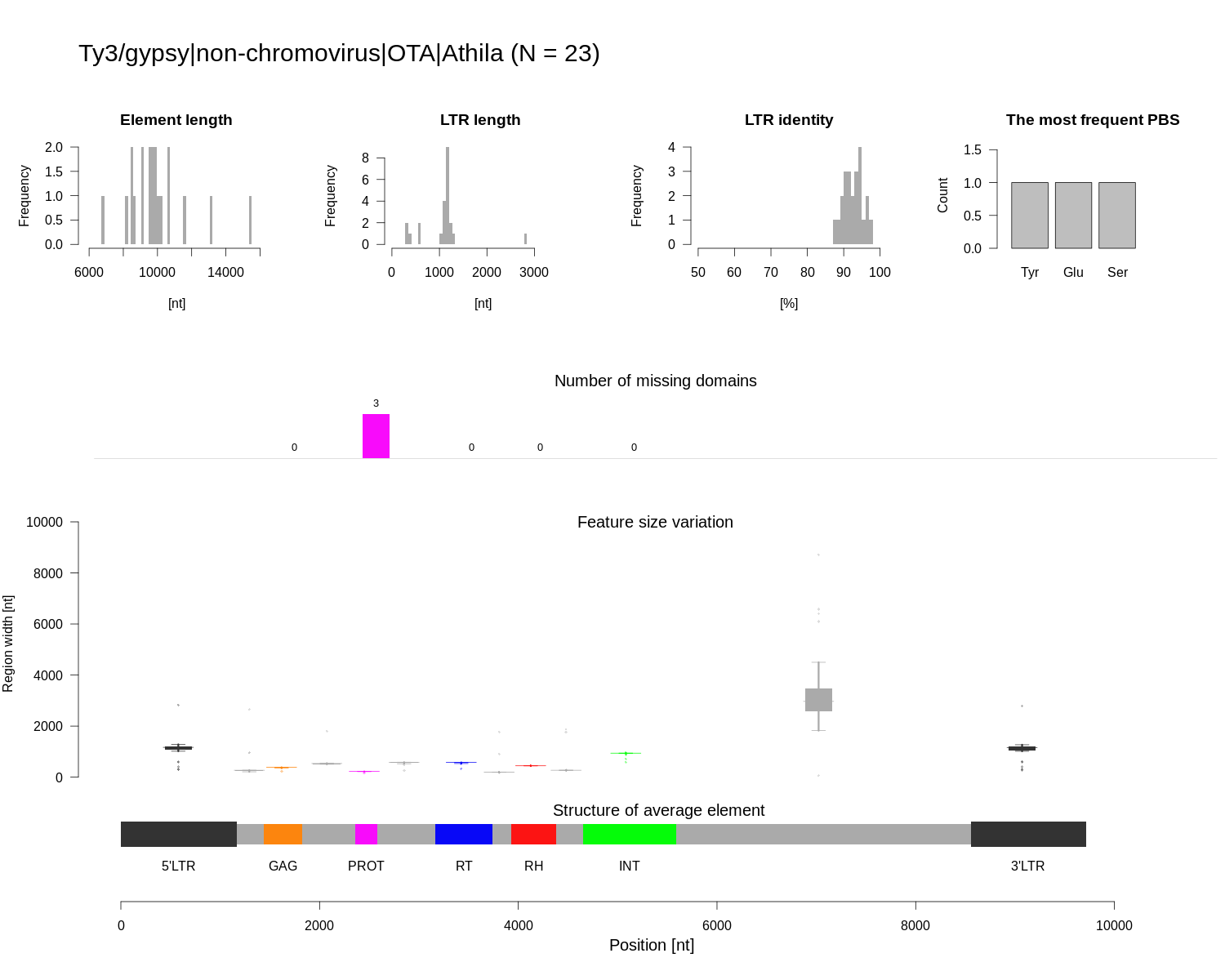

### Class_I_LTR_Ty3_gypsy_non_chromovirus_OTA_Tat_Ogre_structure.png

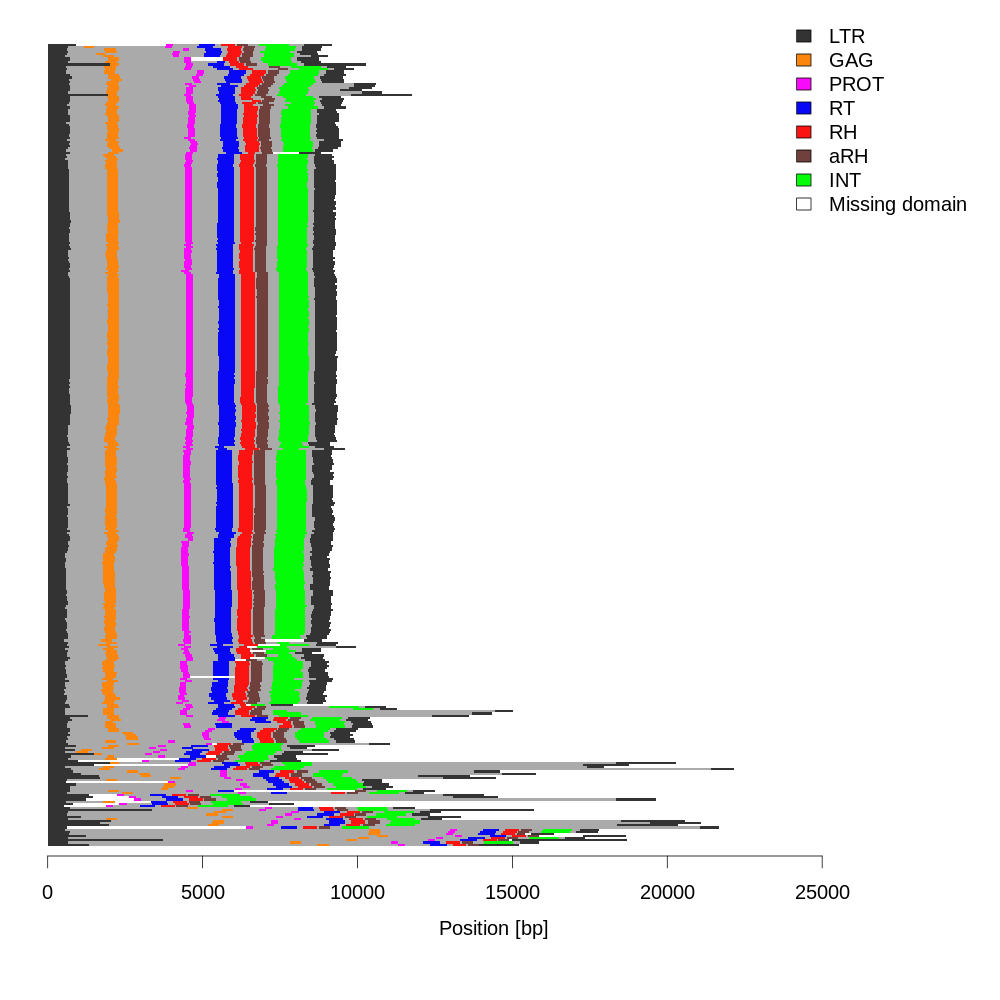

### Class_I_LTR_Ty3_gypsy_non_chromovirus_OTA_Tat_Ogre_structure.png

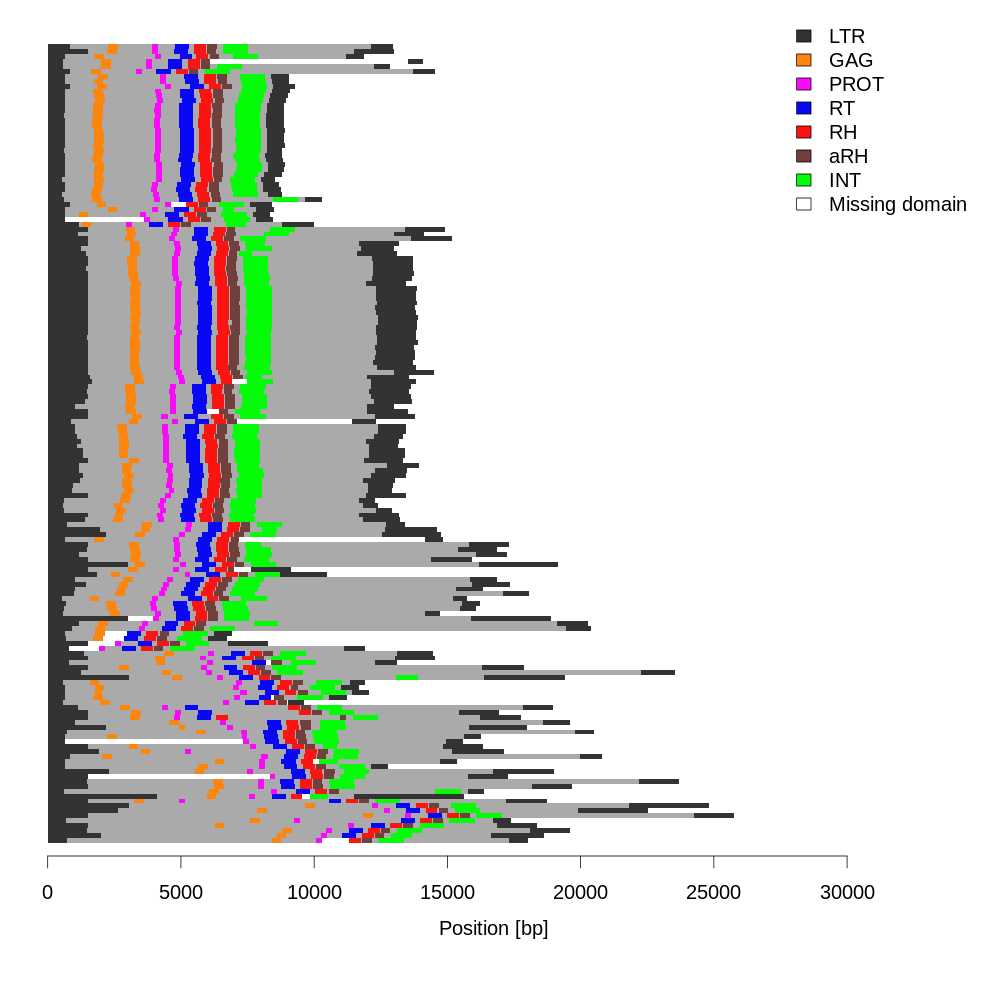

### Class_I_LTR_Ty3_gypsy_non_chromovirus_OTA_Tat_Ogre_summary.png

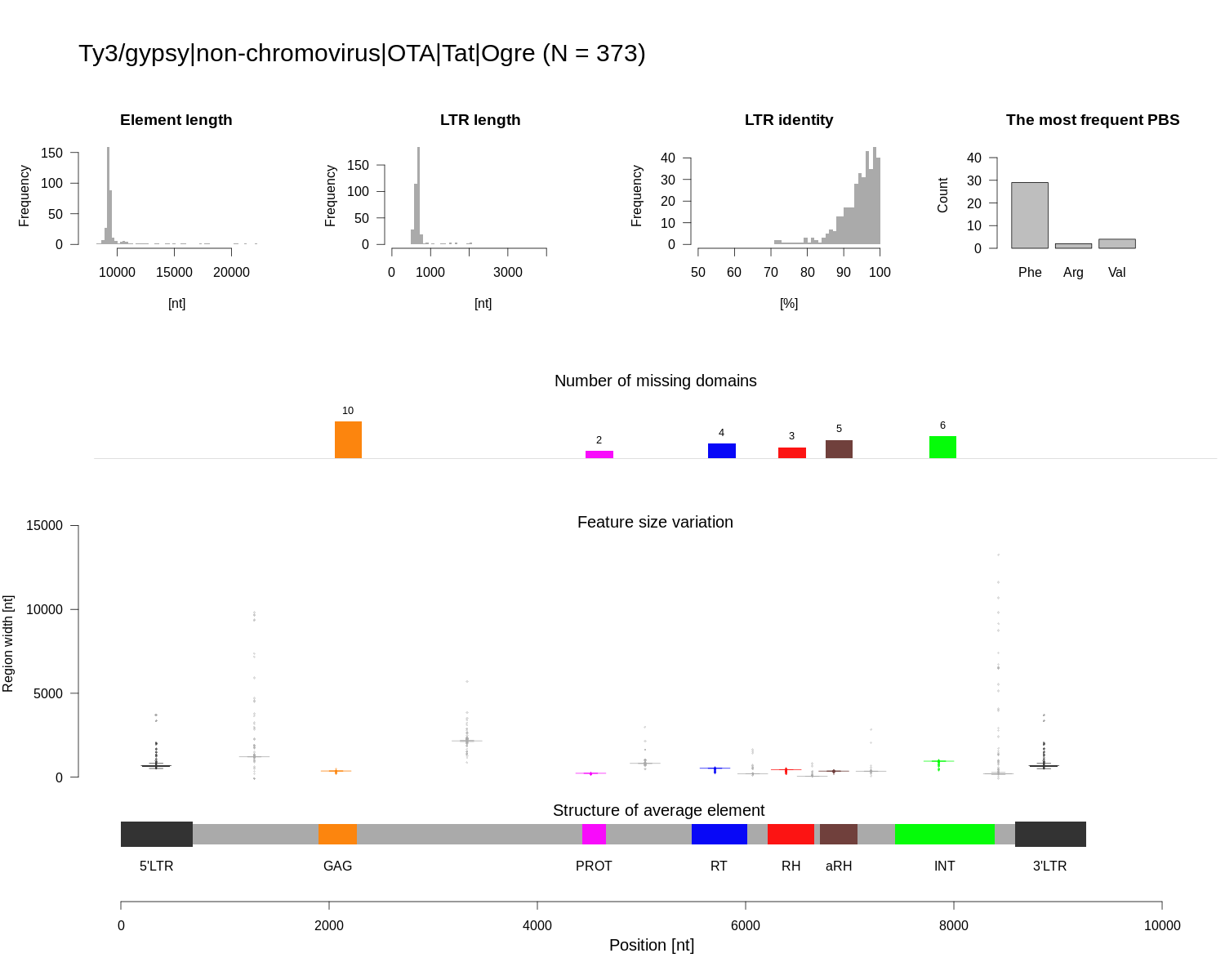

### Class_I_LTR_Ty3_gypsy_non_chromovirus_OTA_Tat_Ogre_summary.png

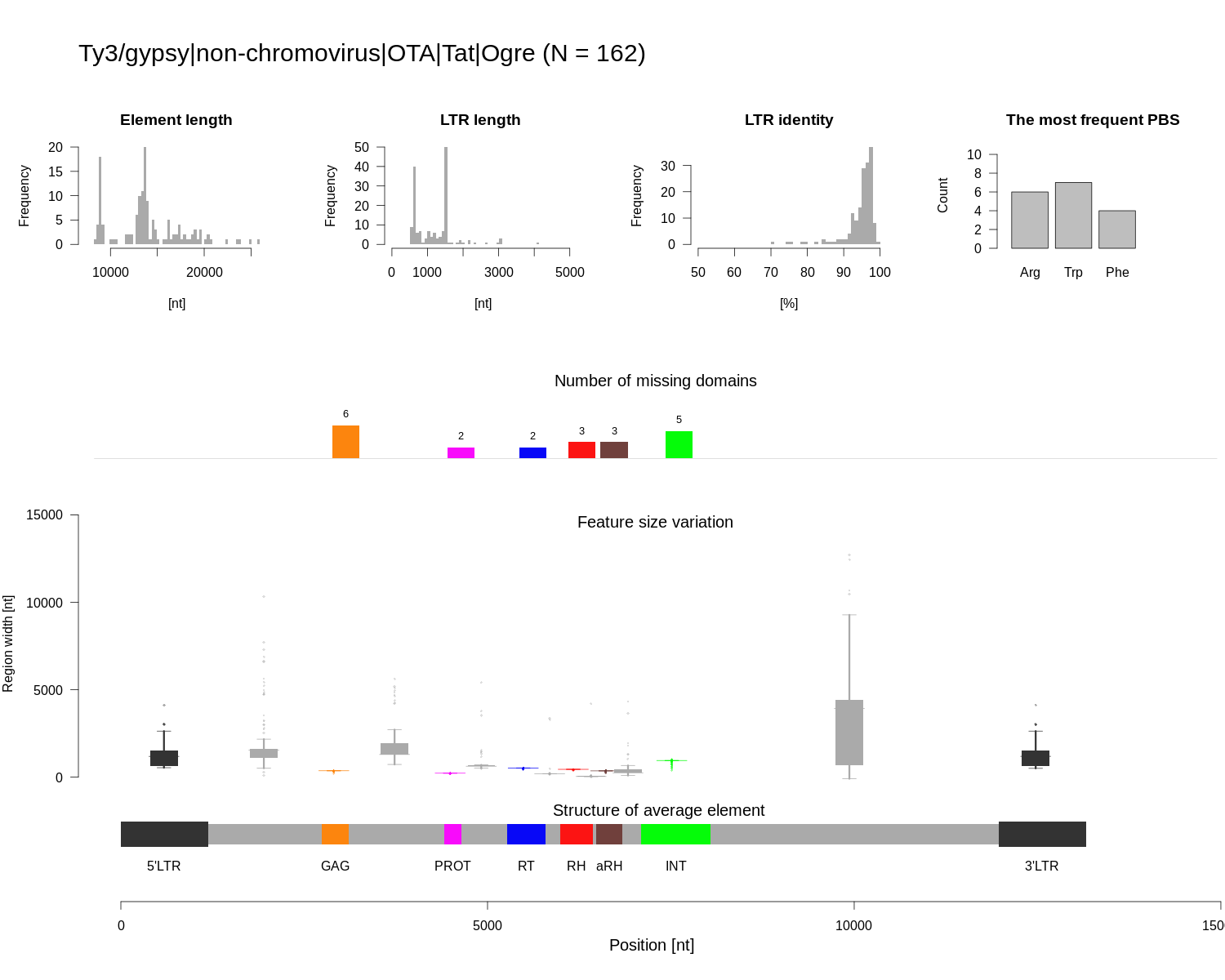

### Class_I_LTR_Ty3_gypsy_non_chromovirus_OTA_Tat_Retand_structure.png

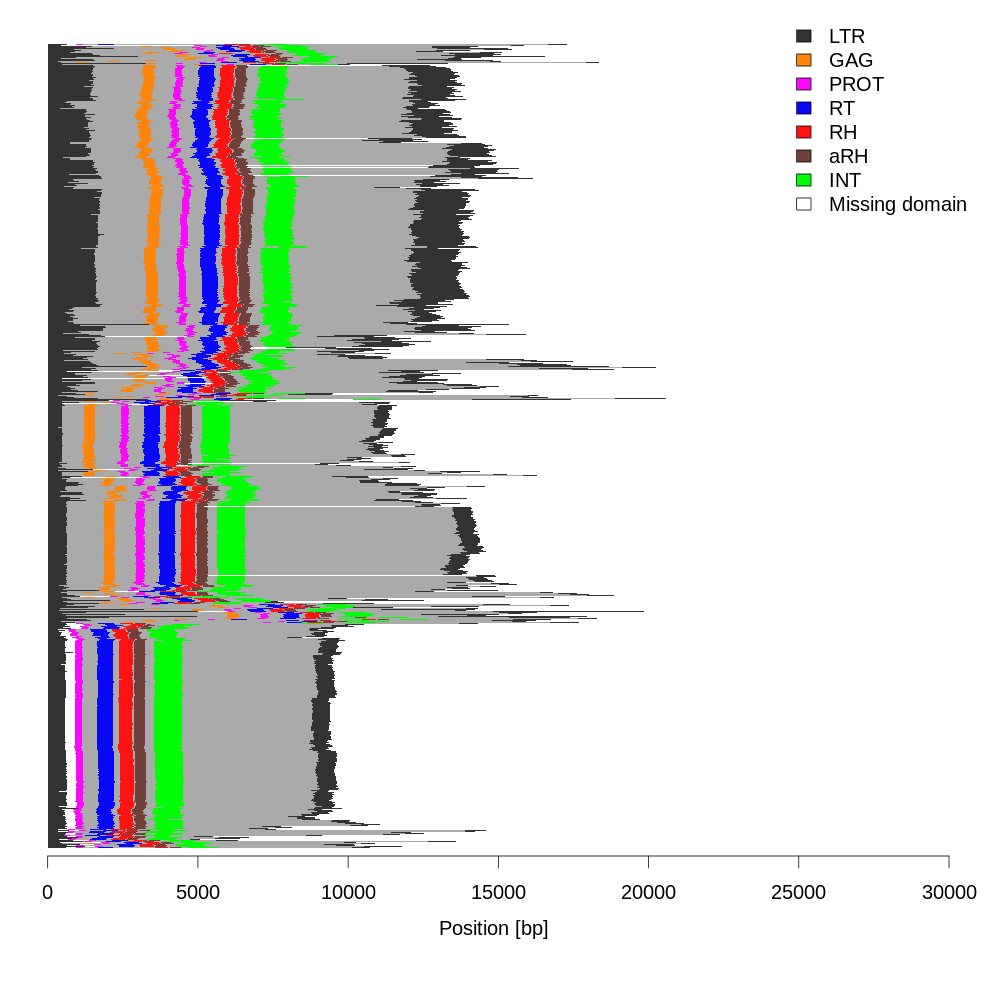

### Class_I_LTR_Ty3_gypsy_non_chromovirus_OTA_Tat_Retand_structure.png

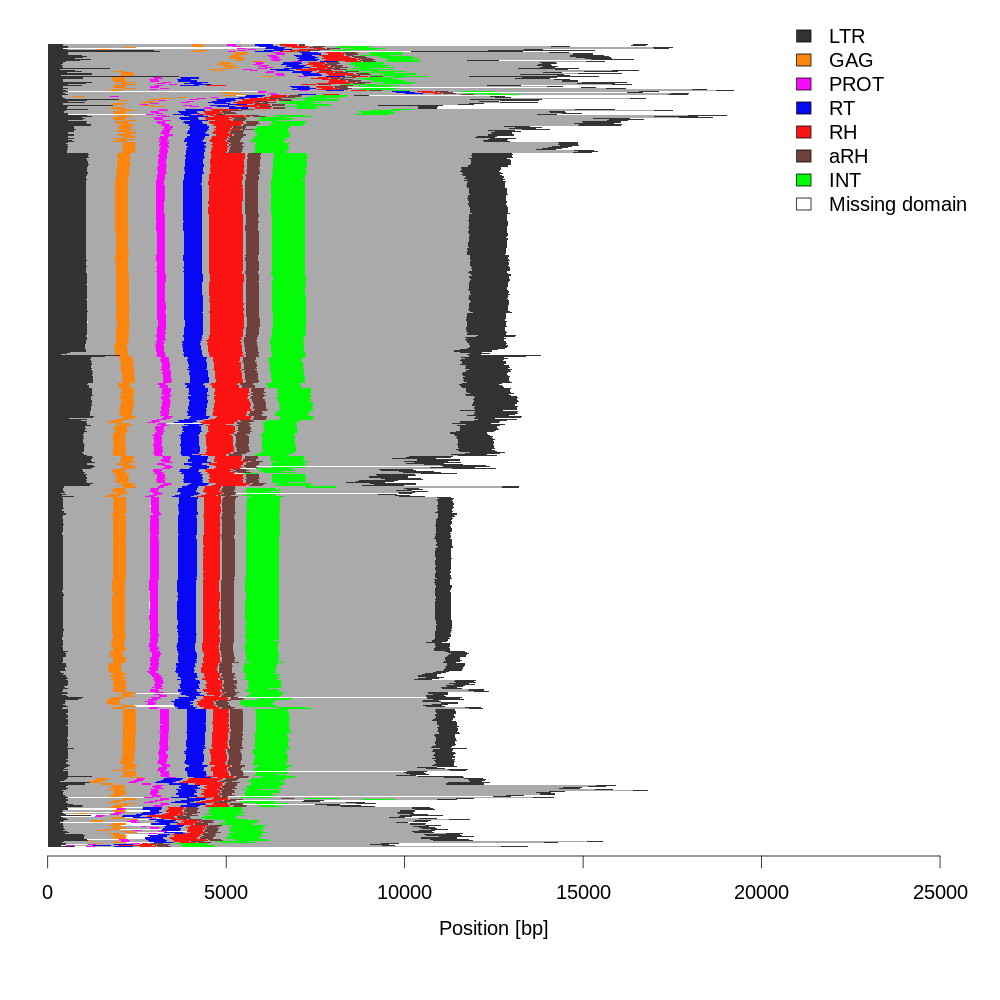

### Class_I_LTR_Ty3_gypsy_non_chromovirus_OTA_Tat_Retand_summary.png

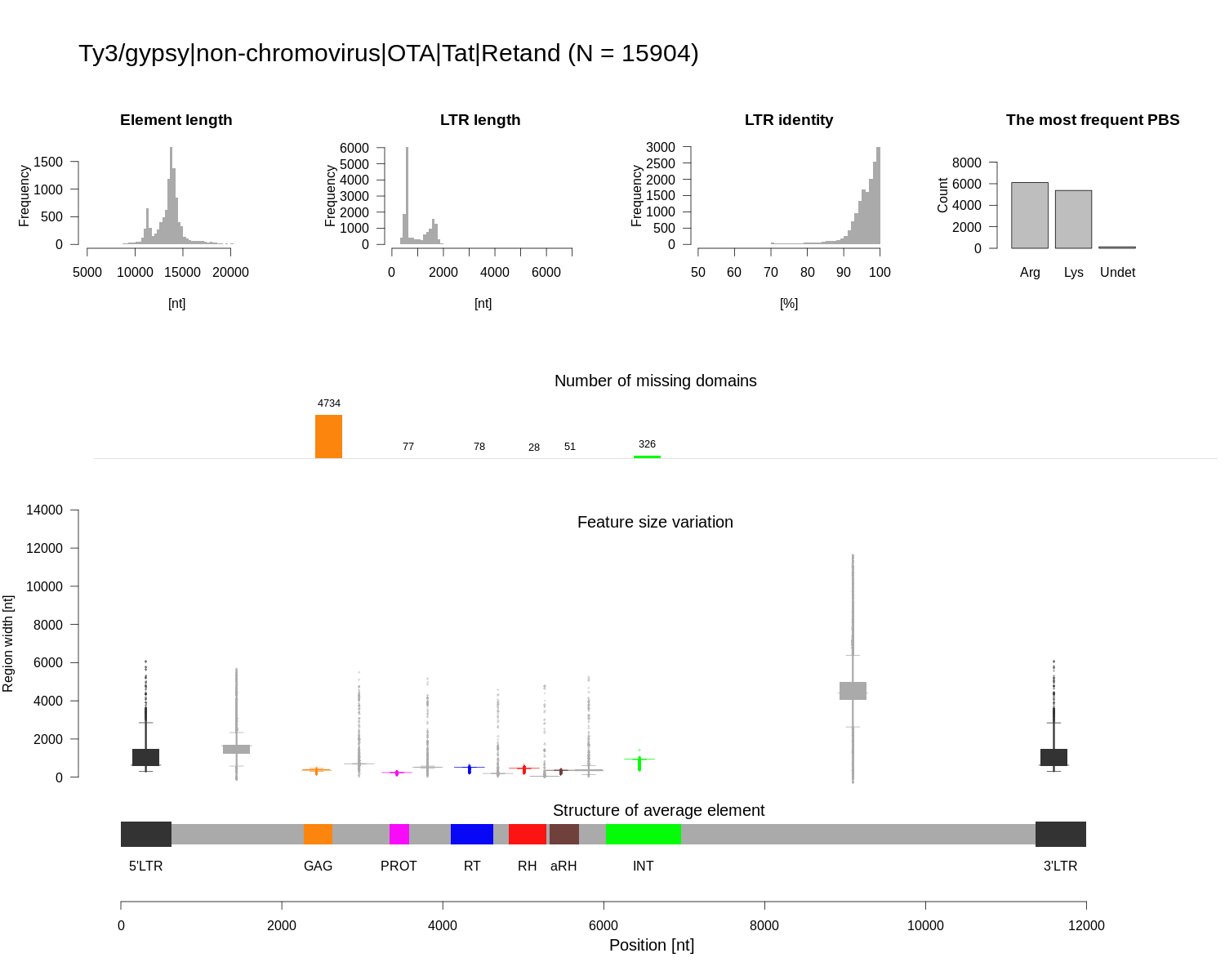

### Class_I_LTR_Ty3_gypsy_non_chromovirus_OTA_Tat_Retand_summary.png

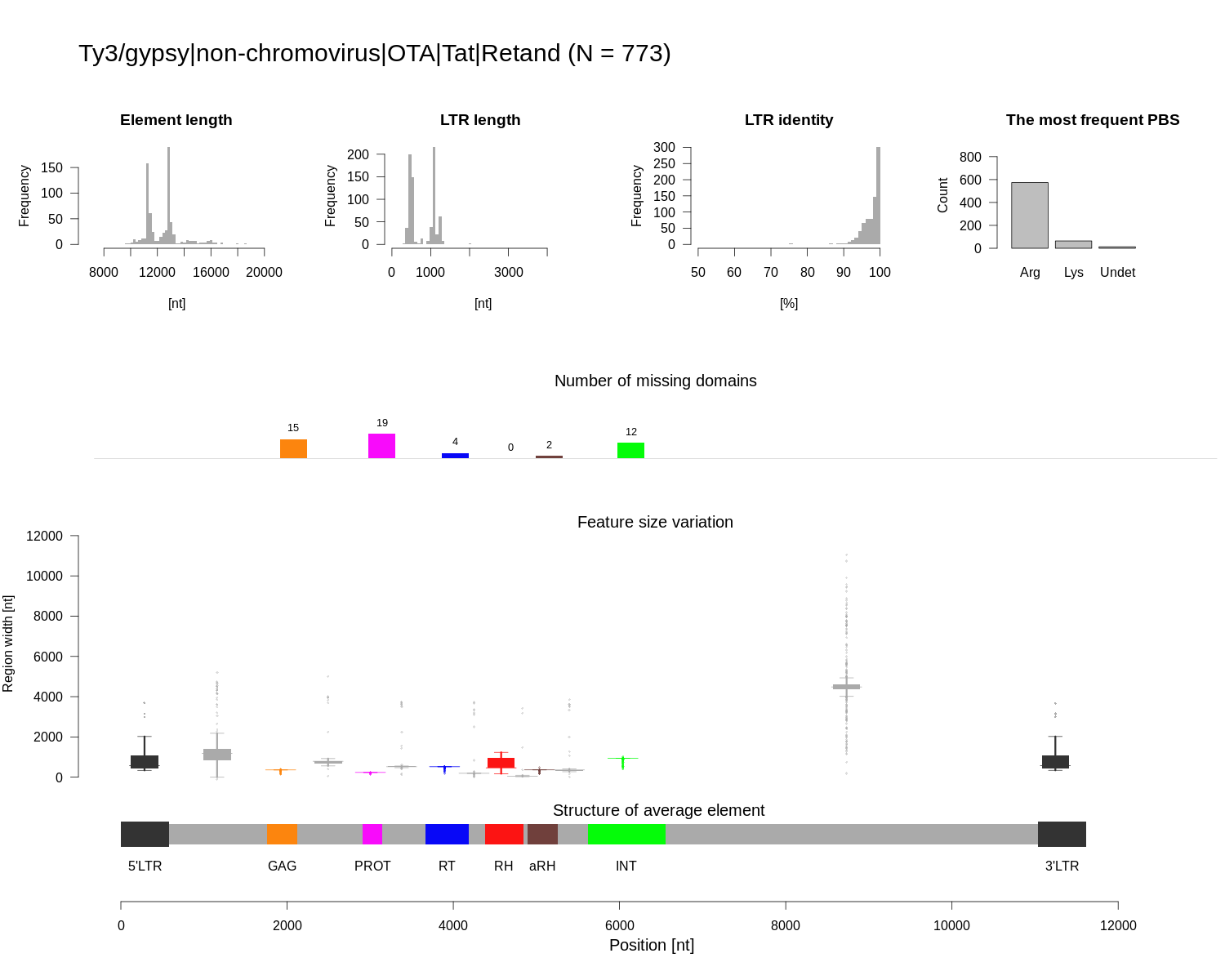

### number_of_elements.png

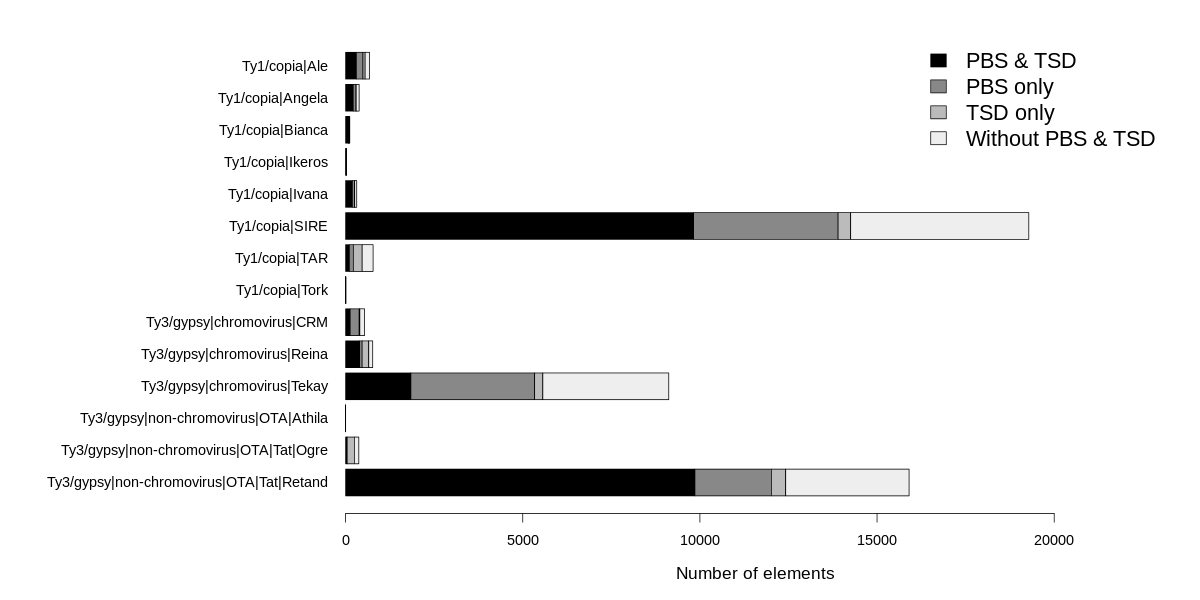

### number_of_elements.png

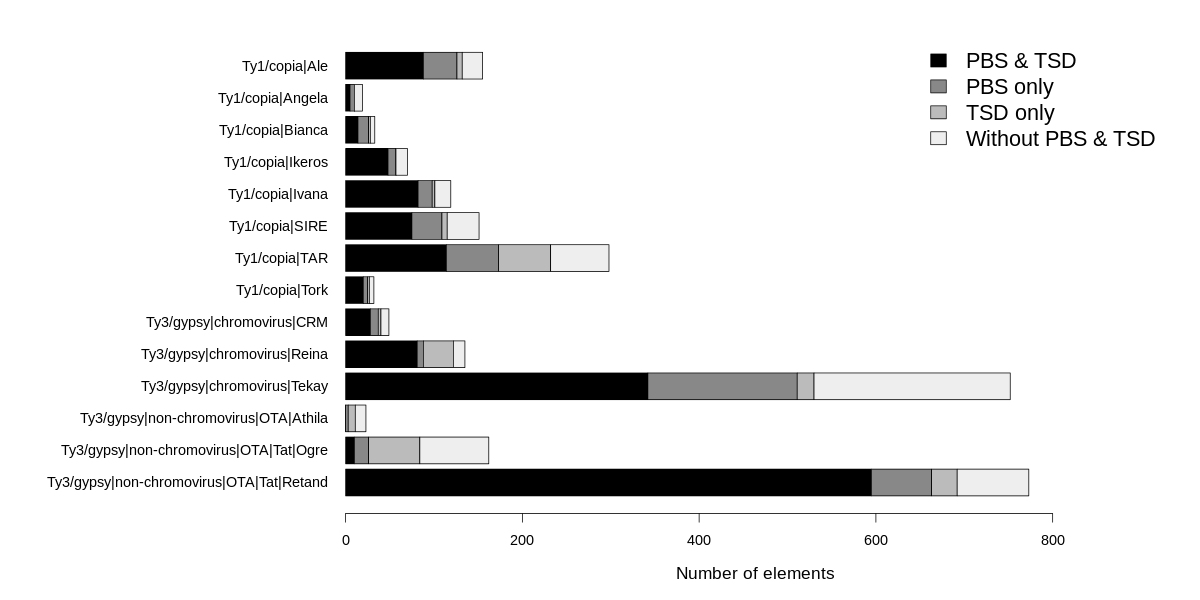
